# Genetic diversity landscapes in outcrossing and selfing *Caenorhabditis* nematodes

**DOI:** 10.1101/2022.12.13.520296

**Authors:** Anastasia A. Teterina, John H. Willis, Matt Lukac, Richard Jovelin, Asher D. Cutter, Patrick C. Phillips

## Abstract

*Caenorhabditis* nematodes form an excellent model for studying how the mode of reproduction affects genetic diversity, as some species reproduce via outcrossing whereas others can self-fertilize. Currently, chromosome-level patterns of diversity and recombination are only available for self-reproducing *Caenorhabditis*, making the generality of genomic patterns across the genus unclear given the profound potential influence of reproductive mode. Here we present a whole-genome diversity landscape, coupled with a new genetic map, for the outcrossing nematode *C. remanei*. We demonstrate that the genomic distribution of recombination in *C. remanei*, like the model nematode *C. elegans*, shows high recombination rates on chromosome arms and low rates toward the central regions. Patterns of genetic variation across the genome are also similar between these species, but differ dramatically in scale, being 10-fold greater for C*. remanei*. Historical reconstructions of variation in effective population size over the past million generations echo this difference in polymorphism. Evolutionary simulations demonstrate how selection, recombination, mutation, and selfing shape variation along the genome, and that multiple drivers can produce patterns similar to those observed in natural populations. Convolutional neural networks demonstrate the potential for classifying distinct evolutionary scenarios in simulated populations. Distinguishing these forces confidently with the empirical data, however, will benefit from larger population genomic samples from multiple populations and consideration of an even more extensive training set of simulations.

## Introduction

Population genomics aims to infer the evolutionary forces and historical processes that have shaped genetic variation within species while considering location-dependent effects generated by chromosomal structure and other genome-wide influences. Patterns of variation along the genome are driven by a range of factors such as natural selection, patterns of reproduction, genome functional organization, mutational and recombinational landscapes, as well as spatial and temporal population dynamics and demographic history. Because most of these factors act and constantly interact, inferring evolutionary scenarios using DNA sequence information within natural populations can be challenging. While many population genetic models incorporate several of these factors ([1], [2], [3], [4]), the enormous complexity of the problem means that there is no single analytical model that encompasses all of these interconnected processes. Even as the scale and quality of individual-level genomic data within natural populations continues to mount, a great deal of work remains to fully integrate the ways that genetic processes manifest patterns at the whole-genome level in combination with evolutionary processes operating within and between populations and species.

Given this challenge, one promising and integrative approach is to use current knowledge about the genetics of a given species alongside evolutionary simulations of a variety of evolutionary scenarios ([5], [6], [7], [8], [9], [10]) so as to generate a series of null models for hypothesis testing of empirical data. Given recent progress in molecular genetic methods, it is possible to obtain high-quality genetic information on genome properties by assembling chromosome-level genomes ([11], [12], [13]), analyzing genome-wide variation in the rate of mutation ([14], [15], [16]) and recombination ([17], [18], [19], [20]), and measuring functional-genomic patterns of activity ([21], [22]), and to then match these features with population-level whole-genome sequence data ([23], [24]). An important emerging approach is deep learning to use such massive and heterogeneous data to capture and classify hidden dependencies and patterns, a machine learning technique that is already finding applications in functional genomics ([25], [26], [22]), medicine ([27], [28]), and population genetics ([29], [30]). Taken together, high-quality genomic data, population theory, individualized null hypotheses from evolutionary simulations, and approaches to learning to identify signatures of evolutionary forces promise to be a powerful tool in population genetics to tease apart the genetic and evolutionary forces that govern genetic variation within and between species.

Variation in the mating system provides one crucial species-specific factor that influences traits, ecology, and population genetic parameters. For instance, self-fertilization as an extreme form of inbreeding acts to reduce the effective population size and the effective recombination rate ([31], [32], [33], [34], [35]), thereby leading to a reduction in heterozygosity, increased linkage disequilibrium, reduced influence of dominance, and increased variability in evolutionary trajectories due to the enhanced influence of drift and a concomitant reduction in efficiency of selection ([36], [37], [38], [39], [40], [41], [42], [43]). Within animals, the transition from outcrossing to selfing is often accompanied by accelerated reproductive incompatibility and isolation, relaxation of sexual selection and sexual conflict, degradation of mating ability, and the generation of outbreeding depression ([44], [45]). In the context of population genomic analysis, it is the influence of self-fertilization on linkage disequilibrium and the way that it expands the genomic footprint of natural selection that is of particular interest. So the contrast between extreme linkage disequilibrium in self-fertilizing species and natural variability in recombination rate across the genome in outcrossing species provides a unique opportunity to critically examine the interaction between population genetic and transmission genetic processes in shaping molecular variation within species.

*Caenorhabditis* nematodes are primarily outcrossing species with males and females ([46]), with the exception of three predominantly self-fertilizing hermaphroditic species: *C. elegans*, *C. briggsae*, and *C. tropicalis*. The self-fertilizing mode of reproduction appears to have evolved independently within each species and to have done so from an outcrossing ancestor fairly recently ([47], [48]). Sex in *Caenorhabditis* species is determined by sex chromosome dosage (X), with females and hermaphrodites having two copies of the X (XX) and males only one (X0) [49]. Because sex is determined by the absence of the X, males can arise spontaneously from nondisjunction of the sex chromosome during meiosis (∼0.1% cases for *C. elegans* [50], [51]), allowing rare outcrossing to occur. *C. elegans*, a model species for behavior genetics ([52], [53], [54], [55]), development biology ([56], [57], [58]), and experimental evolution ([59], [51], [60], [61]), has a very low rate of effective outcrossing in natural populations ([62], [63], [64], [65], [66], [67], [68]) which makes it an especially good model for investigating the consequences of reproductive mode ([69], [70] [71], [51], [72]). From a population genomic point of view, the genomic landscapes of diversity in all three selfing *Caenorhabditis* species are similar, with a consistent pattern among all chromosomes of higher genetic diversity on the peripheral “arm” regions and lower diversity on the central regions of chromosomes ([65], [66], [73], [74]). This pattern closely mirrors the chromosome-level pattern of recombination rate, which is high in chromosome arms and low in centers, with the latter occupying about half of the chromosome length ([65], [75], [76], [74]). To date, however, no comprehensive population genomic data are available for outcrossing *Caenorhabditis* species.

How might outcrossing be expected to affect the distribution of diversity across the genome? In this study, we use a previously constructed chromosome-level assembly ([77]) and a newly generated high-density genetic map to examine whole-genome DNA sequence diversity for *Caenorhabditis remanei*. *C. remanei* is an obligate outcrossing nematode that has a significantly larger effective population size and levels of molecular polymorphism than selfing species ([78], [79], [80], [81], [82]). To provide a uniform basis for comparison across species, we also reanalyzed a local sample of *C. elegans* from Hawaii (from [83]) to compare it with *C. remanei* using the same set of diversity statistics. In addition to inferring demographic histories, patterns of linkage disequilibrium, and genome-wide patterns of divergence, selection, and the spectrum of nucleotide substitutions, we performed evolutionary simulations under different evolutionary, mutational, and recombinational scenarios to compare patterns of diversity in *C. remanei* and *C. elegans* with theoretical expectations. We then employed convolutional neural networks to classify evolutionary scenarios in simulated and empirical data and discuss the caveats and perspectives of this approach. We find that overall levels of diversity are strongly determined by the mode of reproduction and that finer chromosome-level differences in polymorphism are governed by the interaction of selection, mutation, and recombination, indicating that a full understanding of evolution, demography, and genetic transmission are needed to understand whole-genome evolution.

## Results

### Recombination landscape of *C. remanei*

We reconstructed the first genetic map for an outcrossing species of *Caenorhabditis*, *C. remanei*, using 341 F2 individuals from crosses of two inbred lines (PX506 and PX553) generated from isolates collected near Toronto, Canada ([77], Fig 1, S1 Table). Of the 1,399,638 polymorphic sites among the parental strains, on average 106,071 markers were covered by bestRAD-sequencing. Full filtration for informative markers yielded 7,512 total sites across the genome, which in turn were used to construct the genetic map. The total length of the genetic map was 288.72 cM. Of this, 40.99 cM is the X chromosome determined via the female parent, and the remainder is the result of sex-averaged maps for each autosome (see details in S1 Table). The X chromosome has lower rates of recombination than the sex-averaged map of autosomes, which is likely driven by differential control of recombination on the sex chromosome (for example [84], [85], reviewed in [86]), and/or by unequal recombination rates in males and females ([87], [88], [89], also reviewed in [90]). Our crossing approach precludes the construction of sex-specific maps, although examining any potential differences in recombination between males and females could be of interest to future studies.

**Fig 1.**
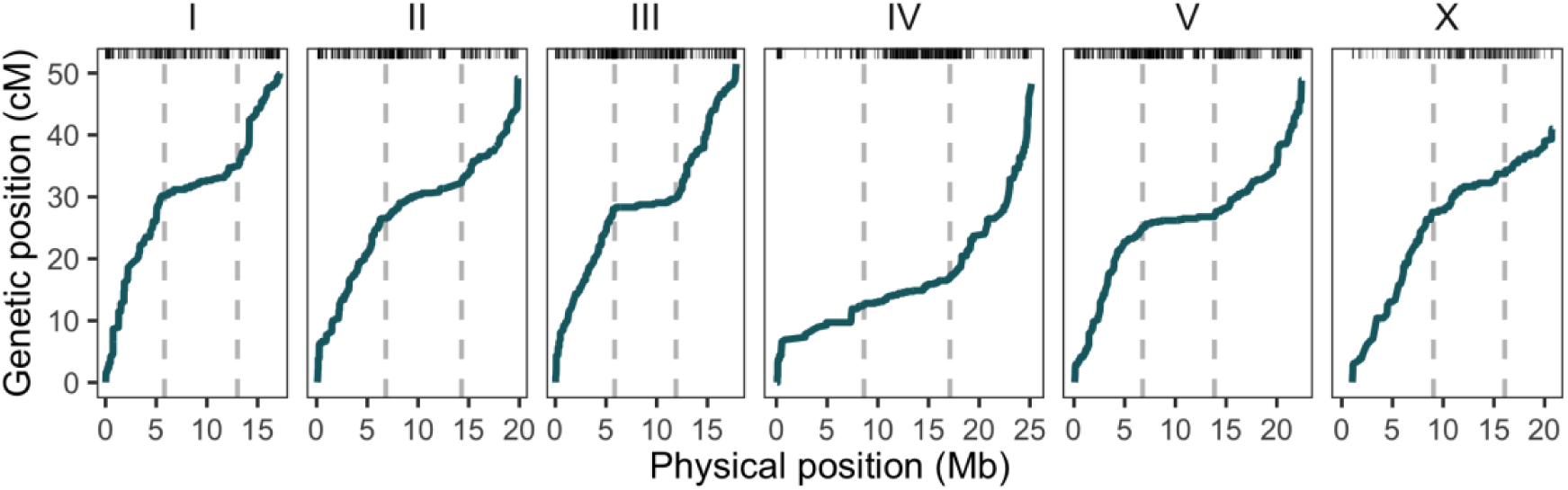
Marey map of genome-wide patterns of *C. remanei* recombination. Rugged marks on the top reflect the density of markers, while the dashed lines show the boundaries of the central domains (see Table 1). The recombination landscape of *C. remanei* resembles that of other Caenorhabditis species, *C. elegans* and *C. briggsae*, with core domains (“centers”) of low recombination and peripheral domains (“arms”) of uniform and high recombination.

**Table 1.** Positions of low recombination domains on chromosomes of the C. remanei genome obtained from crosses of the PX506 and PX553 strains.

| Chromosome | Left arm ends, Kb (95% CI) | Right arm starts, Kb (95% CI) | Recombination rate, cM/Mb (left, central, and right domains) | Chromosome size, bp |
| --- | --- | --- | --- | --- |
| I | 5,803 (5,753 – 5,853) | 13,007 (12,942 – 13,071) | 5.197, 0.673, 3.529 | 17,247,545 |
| II | 6,828 (6,779 – 6,878) | 14,292 (14,060 – 14,524) | 3.883, 0.808, 2.938 | 19,935,723 |
| III | 5,849 (5,828 – 5,870) | 11,888 (11,863 – 11,914) | 4.791, 0.316, 3.601 | 17,877,849 |
| **IV** | 8,639<br>(8,397 – 8,881) | 17,108<br>(17,072 – 17,145) | 1.447, 0.520, 3.615 | 25,790,997 |
| **V** | 6,767<br>(6,717 – 6,817) | 13,852<br>(13,737 – 13,966) | 3.697, 0.270, 2.531 | 22,502,457 |
| **X** | 9,040<br>(8,960 – 9,120) | 16,070<br>(15,498 – 16,642) | 3.047, 0.879, 1.339 | 21,501,900 |

For the purposes of the population genomic analysis, we are particularly interested in the shape of the recombination landscape across the whole genome. We found the chromosomal recombination landscape to be non-uniform in *C. remanei* in a fashion that is superficially similar to that seen in other *Caenorhabditis* species such as *C. elegans*, *C. briggsae*, and *C. tropicalis* with ends of the *C. remanei* chromosomes having elevated recombination rates and for rates of recombination within a given chromosomal domain (arm or center region) being fairly uniform (Fig 1, [65], [74], [75], [76], [91]). We defined the boundaries of the central domains of lower recombination using stepwise regression (Table 1) and refer to the central domain of lower recombination as “Center” and the peripheral regions of higher recombination as “Arms” in the following analyses. Regions of low recombination in *C. elegans* tend to be somewhat larger in relative size than in *C. remanei*. Specifically, the central domains of chromosomes I, II, III, IV, V, and X of *C. elegans* represent, correspondingly, 48%, 48%, 48%, 52%, 51%, and 36% of the total chromosome (estimated from Table 1 in [65]), while in *C. remanei* these represent 42%, 37%, 34%, 33%, 31%, and 33% of chromosomal length.

### Genetic diversity of *C. elegans* and *C. remanei*

In order to compare the landscape of genomic diversity in the outcrossing *C. remanei* to those of primarily self-fertilizing species, we sequenced 14 diploid genomes of *C. remanei* individuals collected in a forest area near Toronto, Canada (S1 Table). To our knowledge, this work represents the first comprehensive analysis of chromosome-scale patterns of diversity for outcrossing species from *Caenorhabditis*. Consistent with patterns observed for selfing species of *Caenorhabditis*, nucleotide diversity is as much as 40% higher in the regions of high recombination than in the central domains on all chromosomes (*C. remanei* mean *π* ± SD in arms, centers, and total: 1.7 x 10^-2^ ± 4.9 x 10^-3^, 1.2 x 10^-2^ ± 4.2 x 10^-3^, and 1.5 x 10^-2^ ± 5.2 x 10^-3^, see Fig 2). Average levels of polymorphism agree qualitatively with previous estimates from this species based on the analysis of individual genes ([92], [81], [82]).

**Fig 2.**
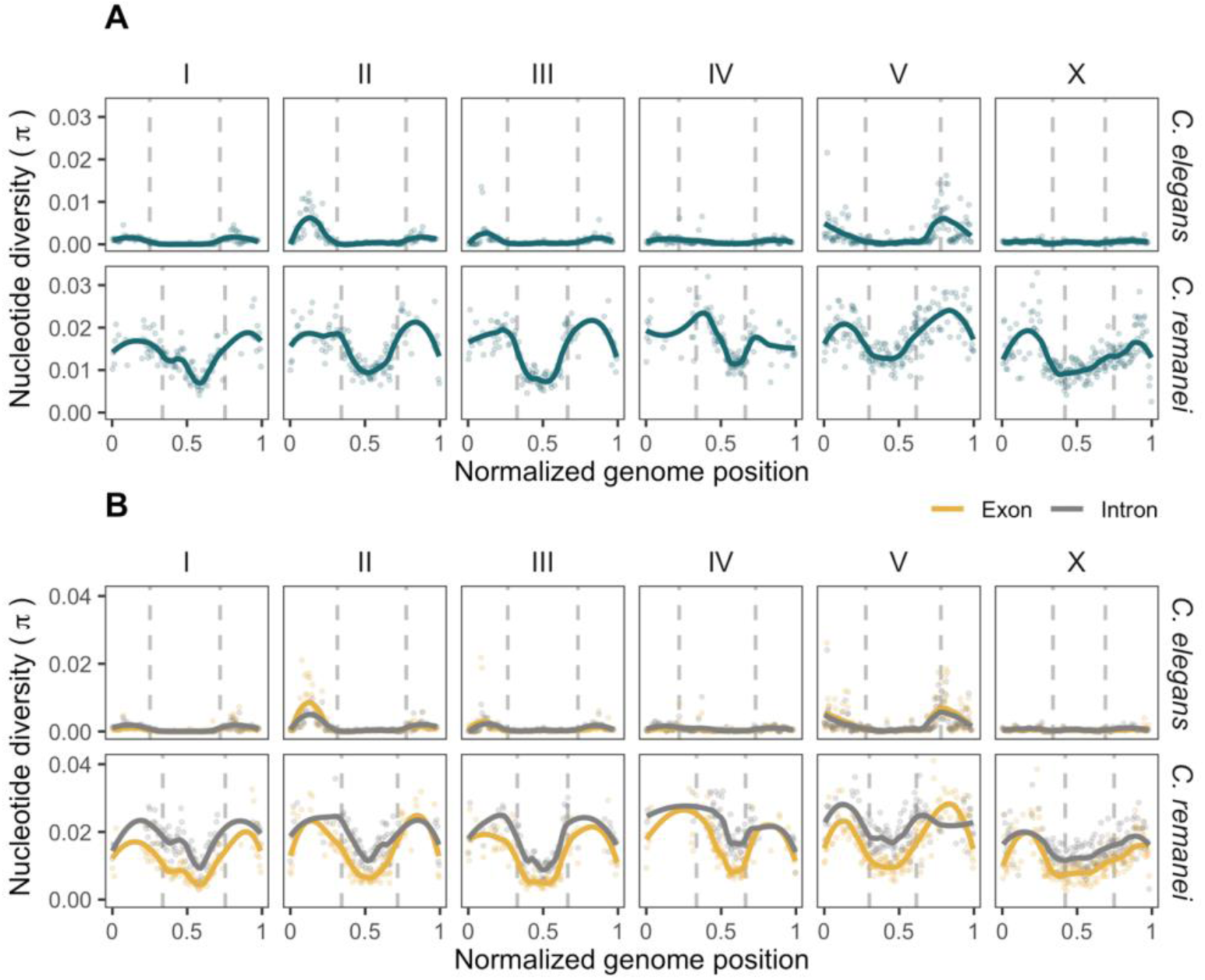
Diversity landscapes of *C. elegans* and *C. remanei*. Dots represent nucleotide diversity per 100-kb window, lines show locally weighted smoothing of these values, and the vertical dashed lines are the boundaries of regions of low recombination central domain for *C. elegans* from [65] and for *C. remanei* from this study. **(A)** Nucleotide diversity per 100-kb sliding windows, windows with less than 10% of coverage were removed. Quantitatively, outcrossing *C. remanei* has one order of magnitude greater nucleotide diversity than partially selfing *C. elegans*. However, qualitatively, both species have significantly greater diversity in the regions of high recombination. **(B)** Nucleotide diversity within exons (yellow) and introns (grey) of protein-coding genes summarized per 100-kb windows, windows with less than 5% of coverage were removed. *C. remanei* shows a large and substantial difference in diversity among exons and introns along the genome, unlike *C. elegans*. This likely indicates that all diversity in *C. elegans* is structured by second-order effects like selective sweeps and/or background selection since both functional and more weakly functional parts of the genome have very similar patterns of variation whereas similar functional elements in *C. remanei* show substantially different patterns of variation.

To directly compare equivalent samples of *C. remanei* and *C. elegans*, we reanalyzed 28 wild *C. elegans* isolates collected at a single location in Hawaii (data from [83], S1 Table) and calculated the diversity landscape across the genome using the same analysis pipeline that we applied to *C. remanei*. Diversity for *C. elegans* was assessed for individual genotypes rather than isotypes, as was performed in the original study, so as to properly retain information regarding genotype frequencies within the population. Consistent with previous reports, this analysis shows that *C. elegans*, like *C. remanei*, has higher diversity levels on the arms compared to the centers ([93], [94], [95], [66], [83]), with mean values (*π* ± SD) in arms, centers, and total of 1.8 x 10^-3^ ± 2.4 x 10^-3^, 5.4 x 10^-4^ ± 1.2 x 10^-3^, and 1.2 x 10^-3^ ± 2.1 x 10^-3^.

The patterns of nucleotide diversity in *C. remanei* are qualitatively similar to *C. elegans* in distribution across the genome but quantitatively different from *C. elegans* in scale by being higher by roughly one order of magnitude. This is consistent with previous observations of substantial reduction of diversity in selfing *vs.* outcrossing species of *Caenorhabditis* ([92], [78], [96], [79], [80], [97], [81], [82]). Highlighting this point, the number of SNVs that we used in the analysis (after filtering, masking of repeats, regions with low mappability, indels, and their flanking regions) was 243,456 variants for the *C. elegans* sample and almost 10x more, 2,365,750, for *C. remanei*.

When comparing patterns of polymorphism between domains of high and low recombination, we find that, as measured by *π*, chromosome arms are significantly more diverse in general than chromosome centers for both species (*C. elegans*: *d* = 0.62, *Z* = 9.28, *P*-value < 1e-04; *C. remanei*: *d* = 1.08, *Z* = 13.96, *P*-value < 1e-04), which also holds when looking specifically within exons and introns (*C. elegans* exons and introns: *d* = 0.52, *Z* = 7.94, *P*-value < 1e-04 and *d* = 0.65, *Z* = 9.73, *P*-value < 1e-04; *C. remanei*: *d* = 1.36, *Z* = 16.16, *P*-value < 1e-04 and *d* = 0.70, *Z* = 8.33, *P*-value < 1e-04). Within a given domain, we find that introns are much more diverse than exons within *C. remanei* (Arm: *d* = 0.43, *Z* = -5.34, *P*-value < 1e-04; Center: *d* = 1.36, *Z* = -15.86, *P*-value < 1e-04) but not significantly different in *C. elegans* (Arm: *d* = 0.06, *Z* = 1.03, *P*-value = 0.8456; Center: *d* = 0.04, *Z* = -0.56, *P*-value = 0.2927). As described below, this difference across functional groups is undoubtedly caused by the fact that the reduced effective recombination rate is *C. elegans* makes these domains highly susceptible to selective sweeps and background selection, which homogenizes diversity across linked genetic elements ([81]). In addition, hyper-divergent haplotypes, located mostly in the regions of high recombination, contribute to the difference in diversity among domains ([98]). Consistent with this idea, variance in *π* is nearly twice as large in chromosome arms as in central domains within *C. elegans* but fairly similar across the chromosome in *C. remanei*. Genomic patterns of other diversity statistics such as θ, Tajima’s D, variance, skew, kurtosis, the number of haplotypes, H1, H12, H2.H1, ZnS, omega (see for details on statistics), and β ([99]) are shown in S2 Fig.

### Divergence and the spectrum of substitutions in *C. remanei*

To facilitate examining the landscape of rates of divergence across the *C. remanei* genome, we reconstructed ancestral states for the *C. remanei* genome by using several genomes of *C. remanei* and a reference genome of *C. latens* ([82], [46], [100]). In so doing, we imputed ancestral states for 56% of the PX506 reference genome (60%, 56%, 58%, 42%, 59%, 68% from the lengths of the chromosome I - X), which included 3,553,584 nucleotide substitutions. All chromosomes had comparable fractions of substitutions from the ancestor (5 ± 0.24%). Calculating the Tamura distance [101] of the PX506 reference genome from the reconstituted ancestral genome shows that overall divergence is 1.6 times greater on the arms than in the central domains (Fig 3A). Divergence tends to be positively correlated with recombination ([102]). While the divergence landscape largely resembles the pattern of nucleotide diversity for the Toronto population of *C. remanei* examined here, chromosome regions showing low divergence tend to be wider than those seen for diversity measures in the regions of low recombination. The divergence time between *C. latens* and *C. remanei* allowed the resolution of population genetic processes (fixation) that are still ongoing among lineages within the Toronto population (segregating variation, Fig 3A, 3C and Fig 2), which provides a more clear definition of the boundaries of the domains. The fluctuations in the ratio of nucleotide diversity over divergence on the boundaries of recombination domains are especially noticeable on chromosomes I, II, III, and V (Fig 3B). These diversity/divergence ratio fluxes could result from selective processes and demographic or non-uniformity in the mutation landscape (Figures S3A and S3B, S3D). The divergence is noticeably lower in exons than in introns, 2.6 times less in the central domains and 2.7 in the arms, and significantly lower in domains of lower recombination (Exons: *d* = 1.13, *Z* = 54.40, *P*-value < 1e-04; Introns: *d* = 1.63, *Z* = 69.26, *P*-value < 1e-04; Total: *d* = 0.28, *Z* = 13.55, *P*-value < 1e-04) indicating strong selection.

**Fig 3.**
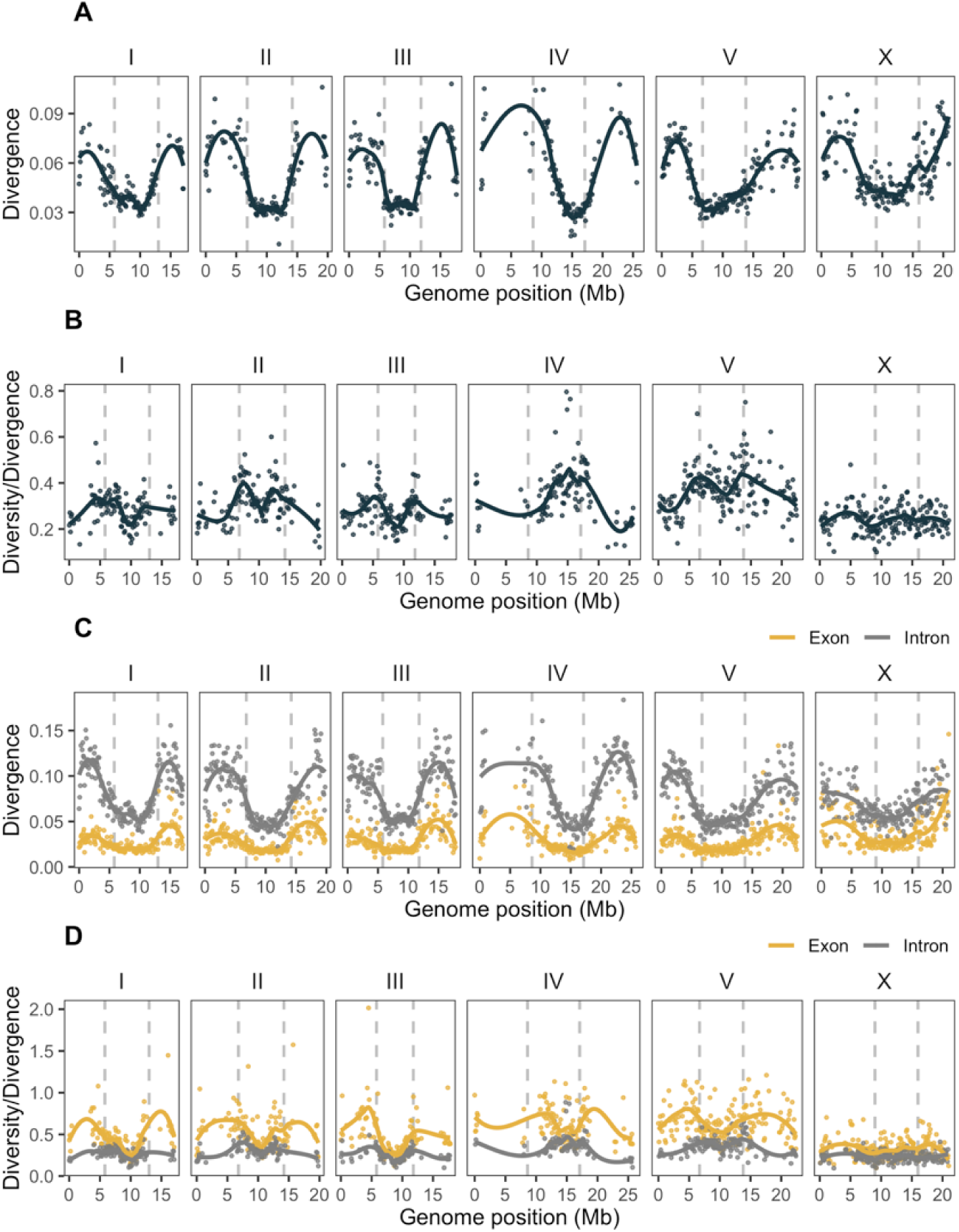
Divergence of the *C. remanei* PX506 strain from ancestral states inferred from the *C. latens* genome. Each point represents the Tamura 1992 distance on 100-kb filtered windows, with at least 30 kb of the window length containing ancestral states, with lines representing the locally weighted smoothing of these values. **(A)** Divergence of the reference genome (strain PX506) from the reconstructed ancestral genome. Divergence in central domains is 1.6 times lower than in the arms, and several highly divergent regions are located on the arms of chromosomes II, III, IV, and X. **(B)** Ratio of nucleotide diversity within the Toronto population of *C. remanei* over divergence shown above. The non-uniform ratio across the genome most likely reflects deviations driven by local variation in natural selection and mutation landscape. **(C)** Divergence of the reference *C. remanei* genome from the ancestral states in exons (yellow) and introns (grey). Only windows with more than 5 kb of ancestral states within exons or introns are shown. Divergence in exons and introns is, respectively, 1.7 and 1.6 times less in the central domains than in the arms. **(D)** The ratio of nucleotide diversity in the Toronto population of *C. remanei* over the divergence estimated in exons (yellow) and introns (grey).

We also used the inferred ancestor to estimate the rate of nucleotide substitution in *C. remanei*, finding that all types of substitutions have a similar pattern of fixation along the chromosomes (S3B Fig). Transition over transversion bias in this comparison is 1.16:1, which is smaller than the 1.5:1 ratio observed within the *C. remanei* population sample from Toronto. Transition substitutions from C→T and G→A were the most common, consistent with previous observations of the *C. elegans* mutation patterns ([103], [104], [105], [106], [98]). The genomic landscape of recombination also has an important effect on the nature of the substitutions, with more C→T and G→A substitutions in the central domain than in the arms, and more C→G and G→C in the arms (S3A Fig, S2 Table). In the *C. remanei* population, the distribution of segregating substitutions along the genome follows comparable patterns to the inferred ancestral patterns (Figures S3B and S3C).

Genome-scale variation in the nucleotide substitution pattern within *C. remanei* could be a consequence of the interaction of recombination, selection, and demographic processes but could also be generated by variation in the mutation process itself. Experiments with mutation accumulation lines in *C. elegans* have shown a 1.2-1.6 times higher rate of base substitutions on the arms relative to the centers of chromosomes ([103]), probably due to mutagenic effects of recombination through double-strand breaks ([107], [108], [109]). Other differences between these genomic regions also might contribute, such as chromatin organization, gene density and their expression, and transposon activity. Accurately inferring the mutation landscape of *C. remanei* will require further research, including the use of mutation accumulation lines and/or extensive sequencing of parent-offspring trios.

### Population structure, outcrossing, inbreeding, and effective recombination

Sampling worms from a single narrow geographic location helps to describe fine-scale population organization. Most of the worms in the *C. remanei* sample formed one cluster of related individuals, with a few more genetically distant worms (S4 Fig). This cluster was most likely formed from individuals from a single-family lineage displaying intensive inbreeding. This view is supported by the relatively high F_is_ values along the *C. remanei* genome (0.38 ± 0.15, S2M Fig).

The inbreeding coefficient was even higher in the *C. elegans* population (0.86 ± 0.22), as expected for selfing or partially selfing species. This F_is_ value corresponds to a 92.5% selfing rate ([110]; s=2F_is_/(1 + F_is_) under the assumption of the equilibrium), which falls within the range of values estimated for other *C. elegans* samples (Table 3). The effective selfing rate estimated from interchromosomal linkage disequilibrium (LD) is three orders of magnitude lower, 99.998% ±0.005% (Table 3). This *C. elegans* sample consisted primarily of individuals derived from several distinct genetic lineages, which have been combined into “isotypes” in [66] and [83] and the CeNDR database (https://www.elegansvariation.org, S4 Fig).

**Table 3.** Outcrossing rate of C. elegans samples reported in different studies. The Method column specifies the approach used to estimate the outcrossing rate.

| Outcrossing rate | Method | Study |
| --- | --- | --- |
| $1.3 \times 10^{-2}$ | Heterozygosity | [94] |
| $2 \times 10^{-1}$ | Heterozygosity | [111] |
| $1.7 \times 10^{-2}$ | Heterozygosity | [62] |
| $7.5 \times 10^{-2}$ | Heterozygosity | This study |
| $1.3 \times 10^{-4}$, $5 \times 10^{-5}$ | Linkage disequilibrium | [94] |
| $1.6 \times 10^{-5}$ to $2.2 \times 10^{-3}$ | Linkage disequilibrium | [63] |
| $< 1.1 \times 10^{-4}$ | Linkage disequilibrium | [73] |
| $2.4 \times 10^{-5}$ | Linkage disequilibrium | This study |

Patterns of genome-wide linkage disequilibrium were drastically different in the *C. elegans* and *C. remanei* samples. *C. elegans* had very large blocks of LD both within and across chromosomes, and LD decays slowly along the entire length of the chromosome (Fig 4 and S5 Fig), which agrees with theory ([41]) and previously reported results ([73]). The degree of LD also varies significantly across chromosomes in *C. elegans* (*d* = 0.58, *Z* = -151.1, *P*-value < 1e-04), which is also consistent with previous observations ([62], [63]). In contrast, LD within the *C. remanei* population decayed rapidly, within a few hundred base pairs on autosomes and a bit more gradually on the X chromosome (Fig 4), consistent with previous sub-genomic observations for this species ([80]). We observe a background inter- and intrachromosomal LD of 0.1, which is more than a null expectation (an inverse of the sample size, 0.04), that is probably related to the high inbreeding or another demographic process in the collected sample (S5 Fig). Consistent with these observations, the inferred genome-wide effective recombination rate in *C. elegans* was on average 8.4 times lower than in *C. remanei* (3.8 x 10^-4^ ± 2.7 x 10^-4^ versus 3.2 x 10^-3^ ± 7.7 x 10^-3^; S6 Fig). However, the effective recombination in *C. elegans* did not follow the recombination domain structure. Here, the difference in actual and effective recombination rates among species is clearly driven by the increase in linkage disequilibrium caused by self-fertilization.

**Fig 4.**
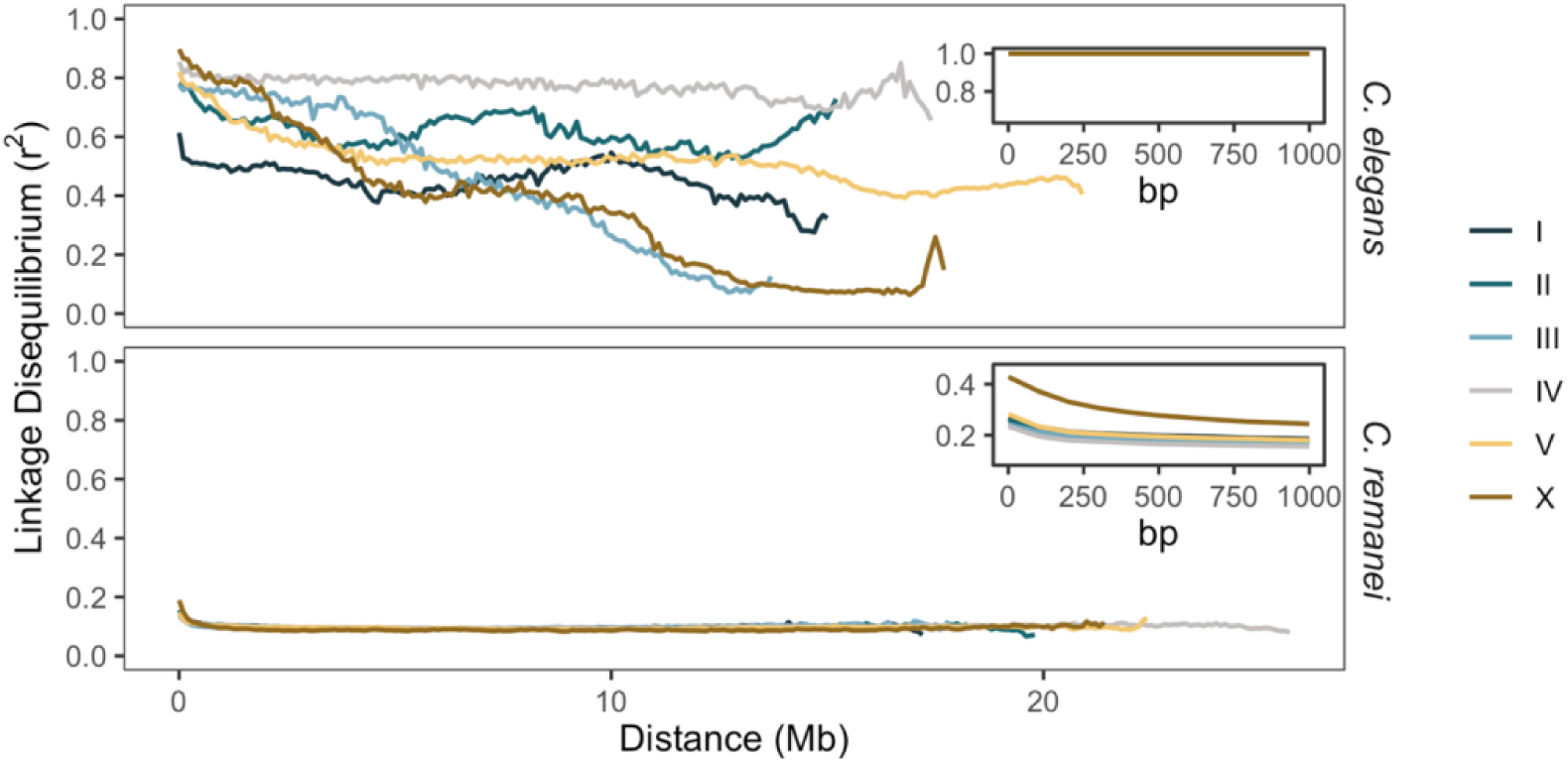
The decay of linkage disequilibrium (LD) along the chromosomes of *C. elegans* and *C. remanei* populations. The median values were estimated from LD between all biallelic polymorphic sites in *C. elegans* and every tenth site for *C. remanei* within either 100-kb windows (overall plots) or 100-bp windows (*C. remanei* inset plot). Each line represents a chromosome.

### Demographic history of *C. remanei* and *C. elegans* samples

Inference of demographic dynamics within both species reveal dynamic changes in population size over time for each (Fig 5). The first striking difference is that the estimated effective population size of our *C. remanei* sample is approximately two orders of magnitude higher than that of *C. elegans*, spanning thousands of generations (Table 4). The size of the *C. elegans* sample from Hawaii has changed dramatically in recent generations, which likely can be attributed to the intricate metapopulation structure ([94], [111]). From a historical point of view, the *C. elegans* sample shows a pattern of a precipitous decline in the effective population size, as previously noted ([73]). In contrast, the Toronto population of *C. remanei* studied here has maintained a consistent, relatively large effective population size but also displays notable fluctuations in size over time both in the past and recently (Fig 5).

**Fig 5.**
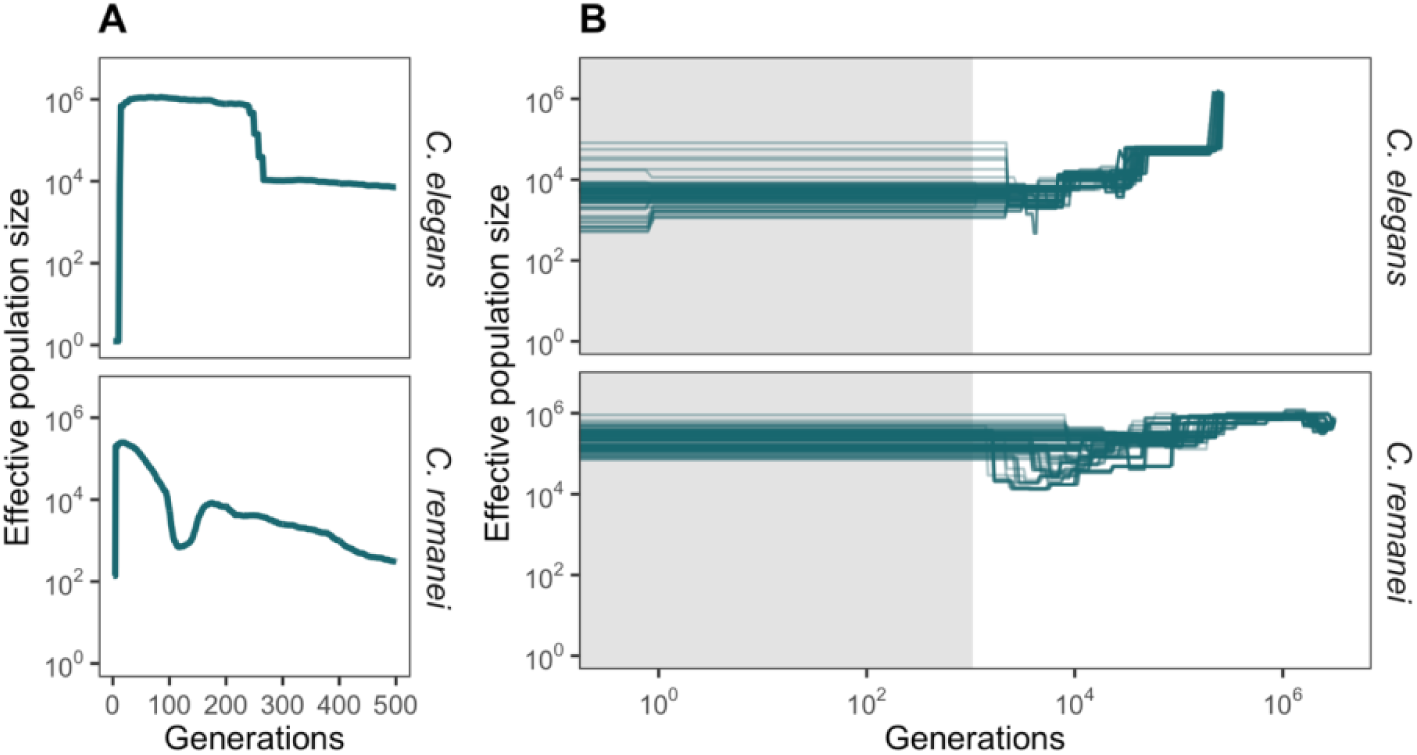
Reconstructed demographic history of *C. elegans* and *C. remanei* populations. **(A)** Recent demographic history inferred from linkage disequilibrium data ([112]). The lines represent the geometric mean of population size from 100 replicates based on autosomes. For *C. elegans*, generation time was doubled, and population size was divided by two. Both species had significant population declines in recent generations. **(B)** Ancient demographic history based on SFS and LD information ([113]). Calculations are based on 100 bootstrapped replicates using eight individuals from each species, with each line representing one replicate. The grey shadow shows the region of recent demographic history, where estimations are less reliable. In this analysis, we used one generation per year, and scaling of the mutation rate (x0.5) and coalescent time (x2) for *C. elegans*. Demography derived from individual chromosomes is depicted in S7 Fig. Population sizes of both species fluctuate significantly over time, which is likely to influence estimates of the coalescent population size. Consistent with this, coalescent population size estimates tend to be one order of magnitude less than the population sizes calculated from polymorphism data (Table 4). The long-term population size of *C. remanei* sample is around two orders of magnitude larger than those of *C. elegans*.

**Table 4.** Effective population size (N_e_) for C. elegans and C. remanei samples from different studies. The Method column indicates the approach used to estimate N_e_. For our study, the 95% confidence intervals for the mean N_e_ are shown in parentheses.

| Species | $N_e$ | Method | Study |
| --- | --- | --- | --- |
| *C. elegans* | $2 \times 10^2 - 9.5 \times 10^3$ | Allele frequencies | [94] |
| | $10^2 - 10^4$ | Allele frequencies | [93] |
| | $8 \times 10^4$ | Allele frequencies | [63] |
| | $10^1 - 10^4$ | Allele frequencies | [111] |
| | $6.5 \times 10^3$,<br>($4.2 \times 10^3$, $8.3 \times 10^3$) | Coalescent, low<br>accuracy | This study |
| | $3.4 \times 10^4$,<br>($3.1 \times 10^4$, $3.7 \times 10^4$) | Allele frequencies | This study |
| *C. remanei* | $1.6 \times 10^5$ | Allele frequencies | [80] |
| | $2.5 \times 10^5$,<br>($2.2 \times 10^5$, $2.8 \times 10^5$) | Coalescent, low<br>accuracy | This study |
| | $1.4 \times 10^6$,<br>($1.4 \times 10^6$, $1.5 \times 10^6$) | Allele frequencies | This study |

To look for site-specific changes in diversity that might be indicative of the action of natural selection, we also inferred population history for the *C. remanei* population using reconstructed ancestral states and the Relate framework ([114]). Overall, the pattern of demographic history using this approach was very similar to that reported above, as well as being concordant across all six chromosomes (see S7 Fig and S8A Fig). However, after adjusting for multiple comparisons, these estimates did not identify any genomic region that had been subjected to substantial positive selection in the past (S8C Fig). Nevertheless, estimated p-values deviated from the expectation of a uniform distribution in a fashion that was more pronounced on the arms than on the centers, which could potentially serve as a signal of non-neutral processes (S8D Fig). This also is consistent with significant and large differences across diversity and divergence in exons vs. introns in *C. remanei* due to selection discussed above (Fig 2B and Fig 3C). We attempted a similar analysis for *C. elegans* but could not reconstruct a sufficient number of ancestral sites from several strains of *C. elegans* and *C. inopinata* (data not shown) to allow the analysis to proceed.

### Evolutionary simulations

The empirical data shows two major features. First, that total polymorphism is lower in the partial-selfing *C. elegans* relative to the outcrossing *C. remanei*, and second, that the genomic landscape of genetic polymorphism is structured and appears to be strongly correlated with domains of high and low recombination. Because many interacting factors can potentially influence these observations, we conducted individual-based simulations to better understand the separate and combined effects of positive selection, background selection, recombination, variation in mutation rates, variation in demography, and variation in rates of partial selfing on population genomic signatures within these species. We looked at three separate scenarios on a wide array of population genetic statistics using evolutionary simulations in SLiM ([9]): the interaction of selfing rate, selection, mutation landscape, and recombination domains; the decay of the ancestral diversity; the effects of fluctuations in population size.

As predicted by theory ([31], [32], [33], [34], [117], [43]), selfing dramatically reduces the diversity in the population, especially when combined with either positive or negative selection (Fig 6A, far right panels). Because recombination has little impact on within-lineage diversity under self-fertilization, any form of selection tends to eliminate the variation at linked sites, often at the scale of the whole genome. As a corollary of this, variation in recombination rate across the genome exerts little influence on the genomic landscape of polymorphism when the selfing rate is high because the genetically effective recombination rate becomes very low. Similarly, the expected variance in evolutionary outcomes is also very low when selfing and selection combine because selection consistently eliminates variation irrespective of when and where new mutations arise within the genome (Fig 6B).

**Fig 6.**
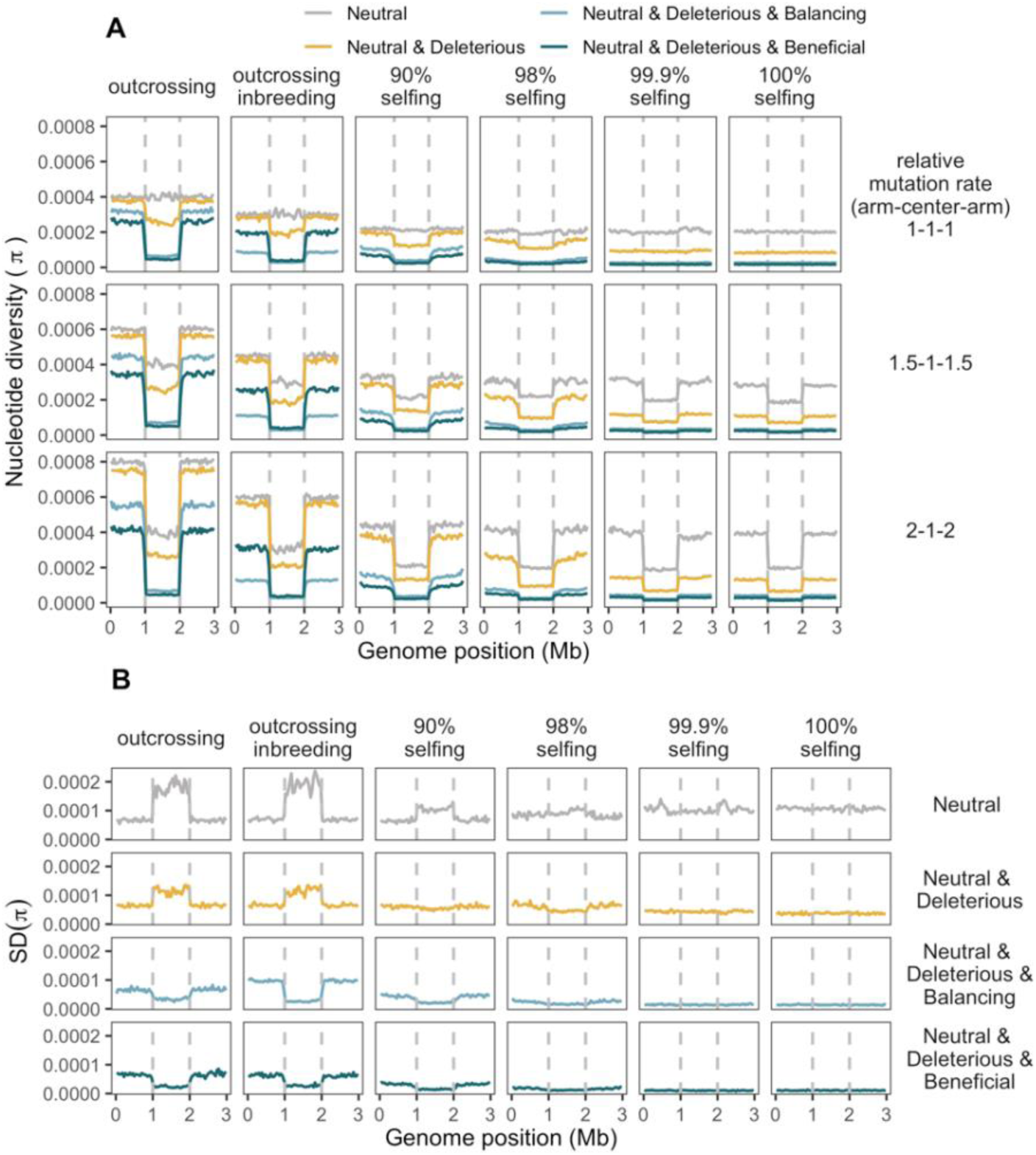
Nucleotide diversity landscapes in simulated populations. Lines represent nucleotide diversity per 40-kb window, the vertical dashed lines are the boundaries of regions of low recombination central domain. Columns show the outcrossing rate, where “outcrossing” means completely outcrossing populations and, in other columns, % specify the percentage of selfing in population; “outcrossing inbreeding” corresponds to scenarios with outcrossing populations that underwent the bottleneck at the very end of simulations (see Methods). Rows represent domain-specific differences in mutation rate, with 1-1-1 as the uniform mutation landscape, 1.5-1-1.5 as 50% more mutations in domains of high recombination, and 2-1-2 meaning two times more mutations in domains of high recombination. Colors show the selection regime (see Methods). **(A)** Mean nucleotide diversity per 40-kb sliding non-overlapping windows. The mean values are lower in regions of low recombination in scenarios with selection and a non-uniform mutation landscape. Higher selfing rate and selection pressure reduce the coalescent time and, consequently, nucleotide diversity. **(B)** The standard deviation of nucleotide diversity in scenarios with uniform mutation landscapes. The variance in diversity gets lower with selection and reduction of effective recombination in non-neutral scenarios and becomes higher with increased selfing in the case of neutrality ([115], [116]).

As even minor amounts of outcrossing enter the population ([118], [119], [120], [121]), however, the situation changes dramatically. In the neutral case, the genomic landscape of polymorphism remains flat regardless of outcrossing rate, as predicted (Fig 6A, [33], [35], [41]). But when selection is introduced, regardless of whether it is positive, negative, or balancing, then regions of high recombination maintain substantially more variation than regions of low recombination (Fig 6A, left most panels). These simulations, in particular, do a very good job of recapitulating the empirical patterns seen in both *C. elegans* and *C. remanei*. Recent inbreeding, as we see in our *C. remanei* samples, does not strongly disrupt this pattern, although the influence of recombination on the genomic distribution in polymorphism is reduced under balancing selection in face of inbreeding, as might be expected. Importantly, however, balancing selection does not generally lead to qualitatively different genomic patterns of polymorphism, nor does it change expectations of the influence of selfing on regions of high and low recombination (see also Figures S10, S11). The variance in outcomes among simulations also displays a recombination-dependent pattern (Fig 6B). Specifically, neutral scenarios have greater variance in regions of lower recombination ([116]). Whereas the variance of π in those domains in non-neutral simulations is lower than in neutral ones and extremely low in scenarios with positive selection.

Although variation in recombination does an excellent job of capturing the genomic differences in polymorphism observed in our empirical examples, it is also possible that these patterns could be caused by domain-specific variation in mutation rate. Indeed, there is evidence that mutation rates are different among recombination domains in *C. elegans* ([106], [103]). If mutation is in fact elevated on the chromosome arms, then polymorphism also increases on those arms, with the difference between arms and centers increasing with the disparity in mutation rate (see Fig 6). As long as there is natural selection and a sufficient level of outcrossing, then changing the mutational landscape does not qualitatively alter expectations of the pattern of genomic variation with respect to the influence of domains of high and low recombination. The situation is more complex under neutrality and/or with a high degree of selfing, in which case the disparity in mutation rate can mimic the pattern of polymorphism expected under the combination of selection and recombination. Thus, while the pattern of variation in *C. elegans* appears to be consistent with some level of outcrossing combined with selection, it is also consistent with selfing and variation in mutation rate correlated with recombination (Fig 6). As outlined below, distinguishing these cases, therefore, depends on the combined effects of all of these factors on additional haplotype and site-based statistics, such as theta, the number of haplotypes, variance, kurtosis, H12, H1.H1, H1, β-statistics, F_is_, omega, that are affected by the mutation landscape and other statistics, such as Tajima’s D and LD based statistics (ZnS), that tend not to be (Figures S10, S11, S12).

Most of the above-mentioned diversity statistics reflect recent evolutionary processes in a population. With simulations, we can explore more distant evolutionary past. One of the statistics that reflects the ancestral relatedness of individuals is the time to the most recent common ancestor (TMRCA). For outcrossing populations, TMRCA is smaller in the regions of lower recombination and reduces in scenarios with selection, with a greater reduction in regimes with beneficial and deleterious mutations. The TMRCA landscape inferred for the Toronto sample of *C. remanei* (S8B Fig) has a distinct shape which may be connected to the population structure and history as well as potential errors in the inference of old tree branches. Selfing, as expected from the theory ([33], [35]), drastically decreases TMRCA due to a reduction of the population size and effective recombination. The variation in the mutation landscape, naturally, has no effect on the TMRCA landscape.

To go further deep in time, we can examine the patterns of divergence from the ancestral genomic state in simulated populations by sampling a random individual at the end of a simulation and comparing its genome to the ancestral “blank” genome (S10O Fig). The divergence is higher in the regions of higher recombination in outcrossing simulated populations and populations with partial selfing, and elevated mutation rate naturally adds on to the divergence, but the reproduction mode has no effect on the divergence pattern. Populations with non-neutral selection regimes show non-uniform ratios of diversity/divergence along the genome. We implemented only simple scenarios with one population to roughly cover the subject of the relationship of diversity and divergence using a single genome comparison and diversity in one population analogous to what we did with empirical data (Fig 3), to investigate nuances of the relationship of diversity and divergence simulations with several populations are required.

Another formal possibility for the genomic pattern observed within *C. elegans* is that, while we might expect this species to exhibit a pattern consistent with selfing, there might be residual, *C. remanei-*like unresolved ancestral variation on the arms due to its transition from outcrossing to selfing (see [98]). To examine this possibility, we used evolutionary simulations to explore the decay rate of ancestral polymorphism from an outcrossing ancestor to a population experiencing either 98% or 100% selfing. For complete selfing and purely neutral variation, the ancestral pattern of variation can indeed persist even after 6N_e_ generations (Fig 7). However, the addition of any form of natural selection and allowing for a small fraction of outcrossing individuals leads to the rapid decay of ancestral diversity, within 1N_e_ generations for 100% selfing and 2N_e_ for 98% selfing. These results are consistent with the expected average coalescent time for the scenario (∼1/N, more specifically (1+F)/2N, where F is the selfing rate; [31], [32], [33], [35], [41], [122]). Here, the variance in the nucleotide diversity is considerably higher in the populations that transition to obligate selfing, which also agrees with previous studies ([123], [35]).

**Fig 7.**
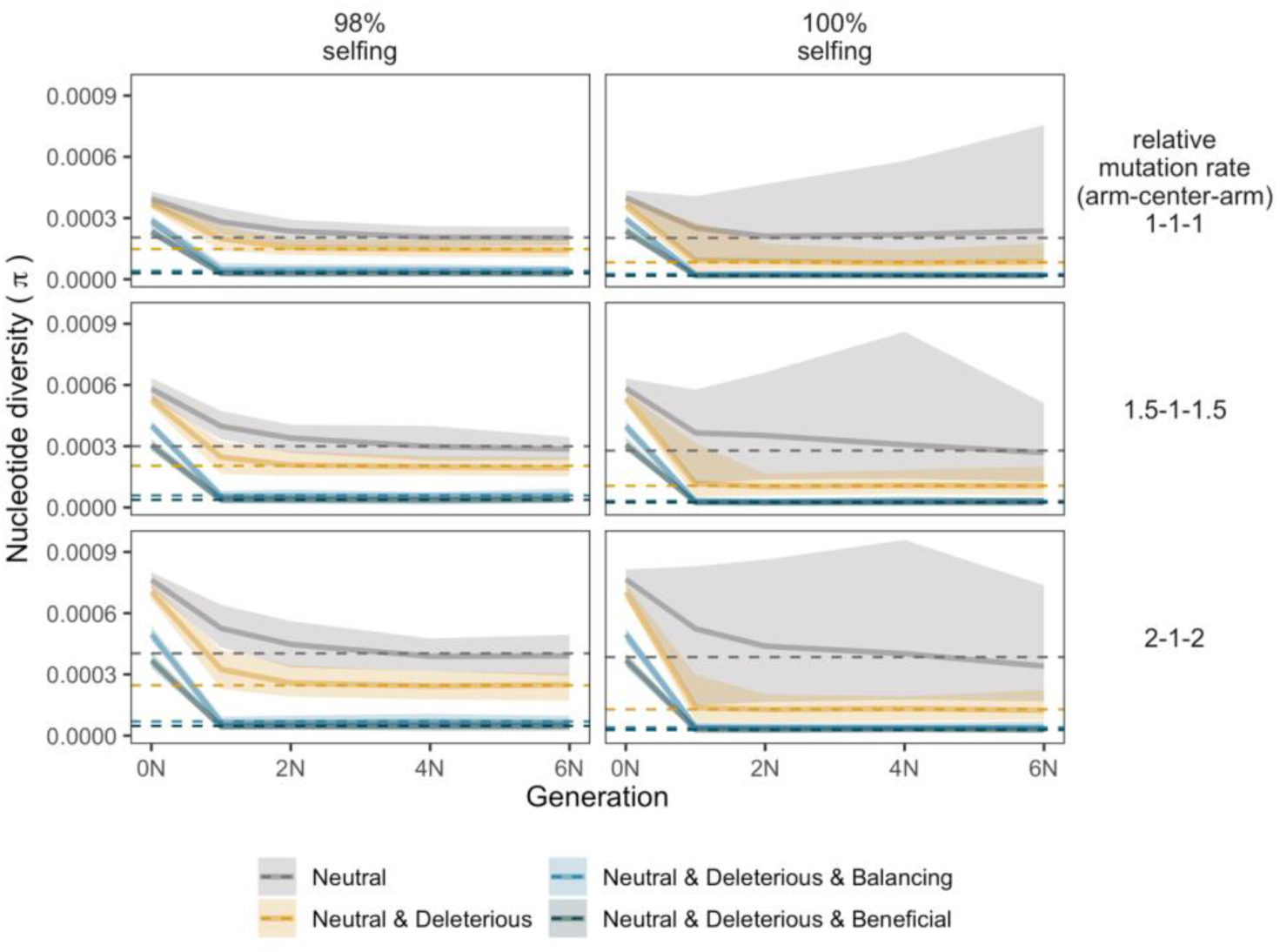
Decay of ancestral polymorphism in simulated populations that transitioned from outcrossing to selfing. The columns indicate populations that switched to 98% selfing (left column) and 100% selfing (right column) after obligate outcrossing. Lines represent the mean nucleotide diversity values estimated from 50 replicates in domains of high recombination, with the shaded areas representing the standard deviation among these replicates. Generations on the ordinate axis show the number of generations after the burn-in. Colors show the selection regime. Each row represents the mutation landscape (see Methods), and the dashed lines represent the nucleotide diversity in the populations without outcrossing ancestors. The average diversity in all scenarios with selection decline rapidly (within ∼N generations) towards levels of diversity in selfing or partially selfing populations. Neutral scenarios in obligate selfing populations have greater variance, which increases over time because these populations are, basically, composed of independent lineages.

The above simulations assume populations of constant size, but we know that variable sizes are the reality for most populations, especially those that might experience more of a “boom and bust” life cycle, such as that experienced by these species ([124]). We, therefore, also conducted a third type of simulation that examined two types of recent change in the population size under neutral scenarios: 1) change from 15K to 5K every five generations for 100 generations after the burn-in; 2) exponential growth by 3% every generation for 100 generations. We then evaluated the diversity statistics and compared them for each simulation after 100 generations of changes in the population size and at the burn-in. S13 Fig illustrates the fold change in a broad range of population genetic statistics in neutral simulations, with large effects in different directions in many of the statistics used in the study, but especially notable in Tajima’s D, haplotype-based statistics, skew, and kurtosis. Importantly, these statistics are sensitive to perturbations in population sizes, which occur naturally in wild worm populations. Likely, the impact of recent demographic shifts on diversity might be less pronounced in the regions of low recombination with selection as the variance in those scenarios is lower (e.g., Fig 6B). Thus, understanding population structure and local dynamics in a population is particularly valuable for interpreting observed patterns of diversity.

### Classifications

Because multiple genetic and evolutionary processes can lead to similar outcomes in terms of genomic diversity (e.g., mutation vs. recombination), we sought a comprehensive approach toward distinguishing major factors structuring genetic variation. We, therefore, trained a convolutional neural network on nine diversity statistics (π, θ, Tajima’s D, variance, skew, kurtosis, ω, and β -statistics) with the intent to classify population characteristics such as selfing rate, mutation landscape, and selection, and then apply this network to our empirical samples for *C. elegans* and *C. remanei*. The network was trained on the first set of simulations described above, with statistics estimated on 40-kb genomic sliding windows and normalized for each simulation. Unfortunately, haplotype-based statistics had to be excluded because they could not be scaled using fixed window size approaches given the drastically different estimated effective population sizes of *C. elegans* and *C. remanei*. Additionally, we excluded simulations with balancing selection as they were mostly classified as outcomes including deleterious and beneficial mutations.

Overall, the network did remarkably well classifying simulation results under most parameter combinations (S14 Fig). Matthews correlation coefficients that show the accuracy of predictions of unbalanced classes for joint-feature predictions was 51.8%, with specific predictions for mutation landscape being 67.7%, selfing rate being 85.2%, and selection regime being 71.7%. The confusion matrix is in S15 Fig. The largest point of error for the network was generated by subtle differences in mutation rate across chromosomal regions, with the network confusing the uniform mutation rate with the 15% increase in mutation rate on the arms (S15 Fig). Many of the population genetic statistics are quite similar for these scenarios (S10 Fig). Other cases that confused the network are distinguishing between simulations with neutral mutations and a combination of neutral and deleterious mutations. This is undoubtedly because even in the deleterious mutation case, many of the mutations are drawn from a distribution in which about a third of the selection coefficients have Ns <1 (S9 Fig). Also, many deleterious mutations tend to have short persistence times, so those present in a given population tend to be recessive and effectively neutral when rare.

Then we classified the normalized diversity patterns from the empirical data from *C. elegans* and *C. remanei* populations using the network trained on the simulations (Fig 8). For *C. elegans*, predictions tended to differ per chromosome. In each case, the selfing rate was predicted to be less than 100%, and some amount of effective recombination was detected, which is consistent with our estimation of outcrossing rate directly from linkage disequilibrium. For whatever reason, Chromosome I from this population of *C. elegans* has a qualitatively different pattern of variation that is more consistent with frequent historical outcrossing and more recent self-reproduction. One of the many possibilities is that it may be connected to the *zeel-1*/*peel-1* system, located on Chromosome I, which is responsible for gametic incompatibility and known to be under balancing selection ([125]). For *C. remanei*, all chromosomes were unexpectedly classified as adhering to one of the selfing scenarios. This is surprising because the network performed very well in distinguishing among outcrossing rates in the simulations.

**Fig 8.**
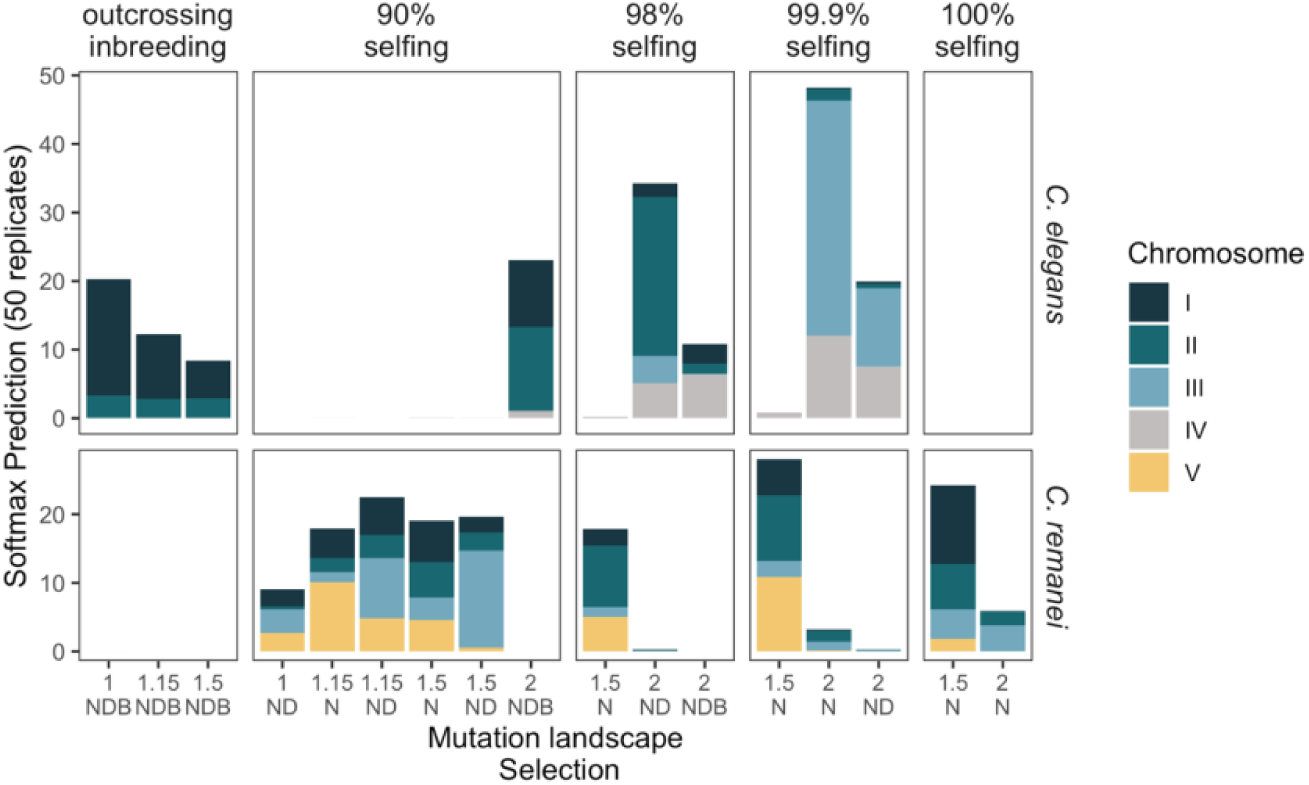
Classification of the empirical landscapes of *C. elegans* and *C. remanei* by evolutionary scenarios. The predictions show the probability (softmax) of each bootstrapped replicate of a chromosome to belong to a specific class of evolutionary scenarios. Only classes with more than 5 in total are shown. Colors represent the bootstrapped replicates of chromosomes; for each nematode, we used only 4 autosomes. Columns correspond to the selfing rate, where “outcrossing inbreeding is outcrossing populations that underwent a bottleneck, and the percentage of selfing defines the selfing rate in simulated populations. The classes represent mutation landscape (“1” – uniform; “1.15” – 15% more mutations on the arms; “1.5” – 50% more mutations on the arms; “2” – 2 times more mutations on the arms. “N”, “ND”, “NDB” mean neutral, neutral, and deleterious, and neutral, deleterious, and beneficial selection regimes. All of the predictions indicate some amount of outcrossing in *C. elegans*. *C. remanei* was not classified as an outcrossing species, which it undoubtedly is. Misclassification might be caused by the complex demographies of nematode populations and/or incompleteness of genomic data, especially on the arms.

To explore these differences a bit more fully, we analyzed the distribution of the simulation and empirical sample statistics using principal component analysis (S16 Fig). In keeping with the above, positions of the *C. elegans* chromosomes in the principal component space tended to be dispersed but within the range of simulated outcomes. In contrast, *C. remanei* tended to be located close together, indicating the similarity of population genetic processes but were at the edge of the scenarios captured by the simulations (S17 Fig). This would seem to indicate other demographic processes not included here have influenced the subtle signals being captured by the population genetic statistics. This conclusion is reinforced by the fact that saliency maps for the network (S18 Fig) show that even small perturbations in theta, kurtosis, and skew in the central domain are enough to change the predictions for outcrossing simulations. That means that the neural network is sensitive to small differences in the ratio of diversity among domains of recombination.

The evolutionary simulations examining recent fluctuations of population size discussed above show that these fluctuations can in principle generate large changes in most of the diverse statistics used in the network classification. It is therefore likely that the main confounder for the *C. remanei* network analysis is demography (see S13 Fig and Fig 5). Unfortunately, there is an effectively infinite number of possible demographic scenarios involving population size fluctuations, and we did have enough computational resources for full exploration of these models at full population size, as such simulations are extremely memory and time-consuming. Refinement of the network model thus awaits additional empirical information and/or an expanded analytical framework.

Finally, at the whole genome level, choices made during bioinformatic processing can also potentially subtly affect diversity statistics in a way that the network might not see. For instance, when filtering the sequence data, even small changes in filtering parameters, like changing the threshold for invariant sites can shift most of the diversity statistic estimations for *C. remanei* (S19 Fig).

## Discussion

The transition from traditional population genetics to molecular population genetics shifted the general analytical framework from alleles at a locus to the nucleotide sequence at a particular site in the genome. The progressive expansion of this framework requires scaling these approaches in the context of broader genome-wide factors such as linkage, recombination, and localized variation in mutation rates, as well as how the impacts of these processes are amplified by population history and structure, and species biology. Attempting to understand the separate and combined impacts of these factors requires comprehensive information about molecular diversity across the genome and a theoretical context in which different alternatives can be rigorously tested. *Caenorhabditis* nematodes provide a natural experiment in which phylogenetically close species have drastically different lifestyles, demography, and genetic processes. With this in mind, we assessed the genomic landscape of various population diversity statistics of populations of *C. remanei* and *C. elegans*. We find that the *level* of diversity is dramatically different across species, with partially selfing *C. elegans* having order of magnitude lower diversity than the outcrossing *C. remanei*, yet that the pattern of genetic diversity is strikingly similar across both species’ genomes, being positively correlated with large-scale transitions in recombination rate between chromosome centers and arms.

### Mating systems and global patterns of diversity

Focusing first on global differences among species, here we used local samples of individuals to facilitate analysis of population history and structure and to allow comparison of features of diversity distributions among outcrossing and selfing species with similar genome organizations. Previous analysis of diversity of *C. elegans* at a global scale ([83], [98]) has demonstrated that even with an overall low genetic diversity expected for a nearly selfing species, a large number of hyper-divergent haplotypes— covering one-fifth of the genome and located mostly on the chromosome arms—have been maintained in that species, possibly because they are involved in adaptation to specific environmental niches. These regions also could result from allelic sequence divergence, analogous to the Meselson effect ([126]) demonstrated for asexual species, expected from coalescent theory when the fraction of outcrossing is considerably lower than the reciprocal of the population size ([118], [119], [122], [127]). Overall, our estimations of the outcrossing rate and population size for the *C. elegans* sample overlap with previously reported values (Tables 3 and 4). It remains an open question if such hyper-divergent regions are present in populations of outcrossing *Caenorhabditis* species, because short-read data, such as that used here for *C. remanei* tends to be difficult to align in hyperdivergent sections of regions of high recombination rate. Long-read data will be required to close these gaps in the coverage.

Looking broadly at haplotype structure across the genome, the rate of decay in linkage disequilibrium (LD) is drastically different in *C. remanei* and *C. elegans*. As predicted by theory ([128], [41]), the outcrossing *C. remanei* shows rapid LD decay, within hundreds of base pairs, and low levels of inter and intrachromosomal correlations. These observations and our estimations of genome-wide effective recombination rate are consistent with previous studies ([80]). However, slightly elevated background LD could indicate a recent inbreeding, consistent with the inferred demographic history (Fig 5) and high F_is_ values (S2M Fig). The “Scottish tartan” pattern of genomic linkage disequilibrium in *C. elegans*, in which associations across chromosomes can often be nearly as strong as associations within chromosomes (Fig. S5), is particularly striking and is consistent with the previous observations of high inter- and intrachromosomal linkage disequilibrium in *C. elegans* ([80]). Because effective recombination rate is a major driver of diversity and linkage along the genome, nematodes provide an especially useful model for studying the evolutionary effects of this factor.

### Genomic landscapes of diversity and factors that affect them in *C. elegans* and *C. remanei* populations

Moving from a global genome perspective to patterns of diversity observed along specific chromosomes requires teasing apart a large number of potentially influential factors (including population history, structure, and demography, natural selection, species biology and development, and reproduction mode, location-specific variation in recombination and mutation, and additional genomic properties such as genome activity and positions of various genetic elements like transposons, genes, regulatory elements, etc.). Using the empirical data from populations of *C. elegans* and *C. remanei* and evolutionary simulations derived here, we step through below how some of these factors contribute to the landscape of diversity.

### Recombination landscape and genomic organization

We constructed the first comprehensive genetic map of *C. remanei*, the first for an outcrossing species in this genus (Fig 2), to guide the understanding of genomic patterns of diversity. The genetic map has a similar structure to other maps constructed for *Caenorhabditis* species ([65], [75], [74]) with large central parts of chromosomes of suppressed recombination ([65], [75], [91]). The recombinational landscape deduced from the population data was also consistent with this pattern. Notably, the X chromosome exhibits the same recombination pattern as autosomes in *C. remanei*, as in other *Caenorhabditis* species ([65], [75], [91]), but with a noticeably shorter genetic map. More research is needed to determine if this is due to distinct recombination regulation on the sex chromosome ([84], [129], [130], [90]) or to averaging of recombination rates between sexes, which might have different recombination regulation, as has been shown for *C. elegans* ([131], [90]). *C. elegans* and *C. remanei*, and some other *Caenorhabditis* species, share other similarities in chromosome organization: central parts of chromosomes display lower recombination and have higher gene density, lower repetitive content, lower GC-content, higher gene expression, and a higher level of inter-chromosomal interactions than the peripheral parts of chromosomes ([132], [133], [77]). As recombination across the genome is one of the critical factors that might affect the shape of diversity landscape, and the genetic map is necessary to model and interpret diversity along the genome, we used the domain-like structure of *Caenorhabditis* chromosomes as the basis for our simulations to allow comparison of diversity landscapes between domains in empirical and simulated populations.

### Diversity landscapes and linkage-disequilibrium

Self-reproduction leads to increased homozygosity and linkage disequilibrium, which is known from theory ([31], [32], [33], [35], [134], [135], [136], [122]) and which has been frequently shown empirically (reviewed in [137]). For *Caenorhabditis* nematodes, genomic landscapes of diversity have already been derived for three selfing species ([66], [73], [74]). In order to compare the diversity patterns of nematodes with different reproduction modes, we obtained the first genome-wide diversity landscape for an outcrossing *Caenorhabditis* species, *C. remanei*, and contrasted its patterns with the selfing *C. elegans*. *C. remanei* has one order of magnitude higher nucleotide diversity than *C. elegans*, due to larger population size, higher effective recombination, and outcrossing, which is consistent with the previous gene-based estimates ([92], [78]). The diversity landscapes in both species follow the domain-like organization of chromosomes with higher diversity levels in the peripheral regions of higher recombination and low diversity in the “central” domains of lower recombination.

In the absence of selection, evolutionary simulations show that nonuniform recombination rates by themselves cannot generate the structure of diversity observed in the empirical data, although it does of course have a very large impact on the majority of population genetic statistics (Fig 6 and S10 Fig, neutral scenarios in the top rows compared to Fig 4 and S2 Fig). In general, these patterns agree with previous predictions expected for a number of these statistics (see Table 1 in [43]).

### Interaction of selection and recombination

Genetic variation and recombination rate tend to be correlated due to the reduction of diversity of linked sites both by positive and background selection ([36], [138], [139], [38], [140], [141], [4]). Positive correlations of meiotic recombination rate and nucleotide diversity have been shown for many species ([142], [143], [144], [145], [108], [146], [147], [148], [149]), however, this pattern is not universal and can be influenced by a variety of factors ([107], [150]). Selfing generates a distinct reduction in effective recombination, thereby reducing the efficiency of both background and positive selection ([151], [152], [32], [39], [122], [42]). Both *C. elegans* and *C. remanei* display higher diversity in regions of higher recombination, which may be a signature of linked selection. This is consistent with prior research showing that *Caenorhabditis* nematodes exhibit substantial effects of background ([144], [147], [153], [154]), positive selection ([66]), and perhaps balancing selection on some loci ([125], [98]). Our observation that exons and introns show very strong differences in diversity in *C. remanei* whereas they are very similar in *C. elegans*, provides strong evidence that selection is much more efficient in *C. remanei* than *C. elegans*, almost certainly because reduced effective recombination rates in *C. elegans*.

The genome organization across *Caenorhabditis* as a whole appears to be likely the result of prolonged continuous selection, as chromosomes within all species are compact, densely packed with genes, and have similar patterns of recombination, and chromatin activity ([77]). They also tend to lack repetitive elements in the regions of lower recombination, which are likely to have been removed by background selection ([133]). Similarly, the central gene-dense regions in domains of lower recombination also tend to be more conserved. For example, within *C. remanei*, reconstructing ancestral states using *C. latens* genome as an outgroup reveals that the central chromosome domains are twice as conserved as the arms, a pattern also observed in *C. elegans* ([132]).

But exactly how do recombination and selection interact to generate these patterns? To address this question, we performed evolutionary simulations using the *Caenorhabditis* domain-like recombination landscape under conditions of different selfing rates, patterns of deleterious and beneficial mutations, as well as mutations under balancing selection. By including selection, we were able to mimic the shape of some of the diversity statistics found in empirical data (Fig 2, S2 and Fig 6, 10, S11, S12). Importantly for the interpretation of diversity within *C. elegans*, even a small proportion of outcrossing (>1/N) was enough to start observing the effects of linked selection, as is predicted by theory ([121], [122], [120]). So, at first glance, it would appear that we can do a very good job predicting patterns of genomic diversity within these species using the right balance of natural selection integrated across domains that differ dramatically in recombination structure. However, diversity is not shaped solely by the forces that either restructure or remove it from the population, but also by mutational forces that introduce it into the population in the first place.

### Mutation landscapes

Mutation is the initial source of genetic variation, and the mutation rate can fluctuate along the chromosome due to such factors as chromatin accessibility, methylation, transcription activity, recombination rate, genomic context, replication and reparation timing ([155], [156], [157], [158], [159], [160], [161], [162], [163], [164]). Nonuniform mutation landscapes have been observed in various species ([165], [166], [167], [168]), including *C. elegans* ([103], [106]), where the mutation rate on the arms is 1.2-1.6 higher than in the central regions. For these nematodes, these differences might be caused by the mutagenic properties of recombination itself, which varies strongly across the genome, or by a variety of other potential factors. To explore potential variation in mutation rates across the genome, we inferred the substitution spectrum for *C. remanei* using our inference of ancestral genomic states. This spectrum has the same dominant type of substitution, C>T|G>A, as *C. elegans* ([103], [169], [170]), and these are the most common polymorphic types of biallelic substitution both in *C. elegans* and *C. remanei* samples (resp. 58.4% and 60% from all types of biallelic substitutions). The fraction of C->T|G->A, A->T|T->A, and C->G|G->C substitutions are different within regions of higher and lower recombination in *C. remanei* (S3 Fig). The similarity of both chromosome organization and substitution spectrum of *C. remanei* and *C. elegans* implies that the mutation rate in *C. remanei* might be greater in regions of high recombination. However, direct mutation accumulation experiments and more detailed analysis of mutational signatures and subtypes are required to fully describe the features of the mutation landscape of *C. remanei*. Similarly, we know very little about the degree of gene convertion within *Caenorhabditis* species, which can lead to an increase in homozygosity in the regions of higher recombination ([127]). An indirect signature of this process is higher GC content in the regions of higher recombination, which we do indeed observe in these *Caenorhabditis* species ([132], [133], [77]). However, since we currently have no basis for parameterizing these potential effects, the potential for gene conversion was not included in our simulations.

Patterns of genomic diversity with varying patterns of mutational input are uncomfortably similar to those produced by an interaction of selection recombination alone. It naturally makes sense that regions with more mutation would also have more standing variation. If higher mutation rates are correlated with higher recombination rates, then at least at a superficial level, it would seem to be very difficult to distinguish among these evolutionary forces (Fig 6 and Figures S10, S11, S12). Our evolutionary simulations showed that variation in mutation rate along the genome affects most diversity statistics that we used in this study, except Tajima’s D, ω, ZnS, F_is_, and TMRCA. In reality, a complex combination of evolutionary forces influences the genetic variability in these nematodes, which suggests that a deeper understanding of the balance of these forces requires more subtle ways of distinguishing among them, as we attempt to do with our classification analysis below.

### History and structure of populations

Organisms live in dynamic environments that change in space and time, which naturally has the potential to dramatically affect population densities and therefore the context for evolutionary change. *C. elegans* populations locally undergo phases of exponential growth in localized areas of vegetative decay, followed by dispersal to habitats with new resources ([171], [124], [68]). Consistent with this, its global population structure suggests metapopulation dynamics of frequent local extinctions followed by recolonization ([172], [94], [111]).

The structure of the local collection of *C. elegans* used here, which is part of a much larger dataset from Hawaii ([83]), shows both multiple divergent lineages and resampling of a few closely related individuals, consistent with the emerging metapopulation paradigm. And indeed, the inferred demographic history of this population suggests massive reduction and fluctuations in population size over time (Fig 5). While the ancient pattern of population history and size is comparable to the previously reported dynamics in *C. elegans* described by Thomas et al. [73], these demographic histories cannot be directly compared on the recent time scale since we employed genomic data from nematodes isolated from one location, whereas they used pseudo-diploids from a “global” sample.

The global population structure of *C. remanei* similarly implies extensive migration across its range ([82], [173]), but local dynamics are still poorly understood. In the Toronto sample of *C. remanei*, we observed elevated inbreeding coefficient and background linkage disequilibrium along the genome, which is likely the result of within-family mating or similar demographic processes in recent generations. Moving back in time, the inferred evolutionary population history of this population displays major fluctuations in the population size of up to two orders of magnitude (Fig 5 and S8A Fig). Even so, the inferred sizes are larger than those predicted for *C. elegans*. One consequence of these estimates is that it means that the total size and overall scale of population dynamics make it particularly difficult to implement this specific demography in our simulations due to excessive computational demands. Nevertheless, our simulations show that continuous exponential population growth had much stronger effects on most diversity statistics than fluctuations in the population size. Similarly, a recent bottleneck in outcrossing populations, which seems to be the case for the *C. remanei* population, can have a significant impact on most of the diversity statistics used in our study (“outcrossing inbreeding” columns in Fig 6, and Figures S10, S11, S12). Overall, however, these demographic effects are minor relative to persistent differences in the mating system.

A potential explanation for the pattern of diversity within *C. elegans* is that the pattern of polymorphism across the genome is a residual echo of ancestral outcrossing, even in the expected homogenizing effects of self-reproduction. In particular, *C. elegans* is predicted to have switched to selfing reproduction within the last ∼4 mya ([47]). To test this hypothesis, we simulated populations exhibiting a discrete change in the mode of reproduction and then tracked how the diversity decays over time. We find that polymorphism from the outcrossing ancestor is predicted to decay very rapidly, so it is very unlikely that the pattern of diversity that we observe in *C. elegans* is the ghost of past outcrossing and instead that it is maintained by ongoing low levels of outcrossing within extant populations or as a result of a complex interaction of selection and metapopulation dynamics.

### Evolutionary simulations and approach to classify evolutionary scenarios

No complete theoretical framework exists to describe how all of the above factors — selection, drift, mutation, genome organization, mating system, and population history — interact to shape diversity at a genomic scale, despite much theory ([174], [42], [120], [122], [43], [175]). Forward-time evolutionary simulation provides an alternative approach to test hypotheses for complex scenarios designed specifically for the species of interest ([8], [9], [176]). Our simulated diversity landscapes compared with the empirical patterns in *C. elegans* and *C. remanei* confirmed the general conclusion that differences in effective population size are responsible for generating the scale and magnitude of genetic variation, whereas effective recombination and selection coupled with mutation and genome organization, shape the distribution of that diversity across the genome (the genomic landscape of diversity). Thus, changing modes of mating across *Caenorhabditis* has had a profound effect on both the scale and shape of the diversity landscape.

We successfully used neural networks to classify various potential evolutionary scenarios using known patterns generated by our forward-time evolutionary simulations that varied patterns of selection, selfing rate, and mutation landscape. Here, perhaps not surprisingly, selfing rate led to the most effective means of classifying potential outcomes. Overall, however, we were not particularly successful in classifying the specific empirical patterns observed in these species (Fig 8). Specifically, whereas patterns for *C. elegans* worked well from the viewpoint of our simulation results, those from *C. remanei* did not, as its patterns of genomic variation were frequently classified as being consistent with selfing rather than outcrossing. Particularly important here seems to be the fact that even small changes in the ratios of some statistics between domains can alter the predicted structure (see an example in S18 Fig; also, Figures S10, S11, S12).

Several factors could make our classification scheme particularly sensitive to these issues. First and foremost, we could not simulate all possible demographic histories of the population, because simulating populations of equivalent size to natural populations, as well as including large-scale changes in population size, turns out to be computationally infeasible. Nevertheless, it is clear that these effects can be substantial, as additional simulations of both recent size fluctuation and moderate exponential population growth showed that recent changes in the population size affect most of the statistics used for classification (S13 Fig). Further, even subtle factors, such as the quality of genomic coverage and bioinformatic assumptions, can lead to significant differences among the diversity statistics (S19 Fig). Long-read data may help close the gap in the coverage and improve the mapping quality for species with such high diversity as we see with these outcrossing nematodes. Lastly, it is possible that the precision of the classification could be improved by including additional genomic drivers such as gene conversion. For *C. remanei*, some of these challenges might be overcome by including broader sampling from across the entire species range so that local demographic differences do not unduly influence the core species-wide diversity patterns generated by conserved genomic processes such as patterns of recombination and mutation. It is equally likely that the sensitivity of this approach can be enhanced when we are able to add in additional complexities such as realistic demographic histories and local spatial and temporal dynamics to the simulations.

## Conclusion

*Caenorhabditis* nematodes provide a useful model to study evolutionary consequences of selfing, as they have highly divergent mating systems while maintaining an overall similarity in genome organization. We demonstrated the recombination landscape of outcrossing *C. remanei* to be similar to *C. elegans*, with extensive domains of lower recombination on the central parts of chromosomes that predict genomic landscape of the diversity in both species. The scale of genetic polymorphism within selfing *C. elegans* and outcrossing *C. remanei*, however, are dramatically different because of large differences in effective population size and demographic history. Although we were able to classify evolutionary forces that shaped diversity in simulated populations, applying this approach to empirical data will require more comprehensive simulations of realistic biological scenarios and additional high-quality empirical genomic data. These findings support the emerging perspective that understanding patterns of variation at any particular site in the genome requires a global perspective of the forces that shape variation across the genome as a whole.

## Materials and methods

### Genetic map for Caenorhabditis remanei

We constructed a genetic map for *C. remanei* from crosses of 2 inbred strains, PX506 and PX553. Initially, *C. remanei* isolates were derived from individuals living with terrestrial isopods (*Oniscidea*) taken from Koffler Scientific Reserve at Jokers Hill, King City, Toronto, Ontario, in October 2008 as described in [177]. Strain PX390 was created from one female mated with 3 males from an isopod. Strain PX393 is from one female and one male from an independent isopod. The strains were propagated for 2-3 generations before freezing. PX506 and PX553 are inbred strains generated from PX390 and PX393, respectively (the parental strain for PX506 was inadvertently specified as PX393 in [77]). To reduce residual heterozygosity, the lines were sib-mated for 28-30 generations before freezing. Nematodes were kept under the standard laboratory conditions according to Brenner [49].

The genetic map was constructed from 4 crosses of *C. remanei* strains PX506 and PX553, (2 crosses of ♀PX506 x ♂PX553 and 2 ♀PX553 x ♂PX506) using parental genotypes and 341 individually sequenced F2 worms, all females. Single L4 animals were digested in proteinase K, and the DNA content was linearly amplified with the phi-29 enzyme (GenomiPhiV3, GE life sciences), then normalized samples were processed for bestRAD sequencing [178] with EcoRI restriction site based adapters. Each multiplexed sample set was sequenced on the Illumina Hi-Seq 4000 platform with four lanes of 100bp paired-end reads (University of Oregon Sequencing Facility, Eugene, OR). Additionally, we sequenced the PX553 parental strain using the Nextera kit (Illumina) and Hi-Seq 4000 platform. The genome sequence of the PX506 strain was generated previously [77].

For parental strains, we checked the sequence quality of reads with FastQC v.0.11.5 [179] and MultiQC v.1.3 [180], trimmed and filtered reads with Skewer v.0.2.2 [181]. The filtered reads were mapped to the *C. remanei* genome (GCA_010183535.1 from the NCBI database) with BWA-MEM v.0.7.17 [182]. Then we filtered reads with SAMtools v.1.5 [183], removed duplicates by Picard tools v.2.17.6 [184], realigned indels, and called variants with GATK v.3.7 and v.4.1 [185]. Variants were filtered with standard GATK hard filters (see discussions in [186], [187] [188]), with only diallelic loci being used in the analysis. We also masked repetitive regions, as well as sites with too low or too high of coverage.

To process the bestRAD reads, we filtered reads without barcodes and flip forward and reverse reads when a barcode was found on the reverse read using Flip2BeRAD [189], demultiplexed reads with process_radtags from the Stacks package v.1.46 [190], followed by an additional adapter and quality trimming with Skewer. Then, we mapped the reads to the *C. remanei* reference genome (GCA_010183535.1) using bwa mem, marked duplicates and recalibrated alignment with variants from the parental, PX506 and PX553, strains with Picard and GATK BaseRecalibrator, filtered reads that do not cover the parental variants, have secondary alignments, or low mapping quality by SAMtools, and called variants with samtools mpileup. We generated the genetic map with Lep-Map3 [191], the order of markers was defined by their positions on the reference genome. The recombination rate per Mb was estimated in R, the boundaries of low and high recombination domains were determined with the pricewise regression by the segmented package [192] in R. For more details on this part of the analysis, see the scripts at https://github.com/phillips-lab/CR_CE_popgen/genetic_map/.

### *C. remanei* and *C. elegans* population genomic data

To study the genomic pattern of diversity in outcrossing nematodes, we sequenced 14 wild individuals of *C. remanei*. Isopods, a phoretic host carrier of *C. remanei* [193], were collected at the same station as the strains used for the genetic map, the Koffler Scientific Reserve in Ontario, Canada, in September 2013, sacrificed within a few hours following collection after having been placed on agar plates seeded with *Escherichia coli* OP50. From each of 14 single *C. remanei* individuals isolated the next day from these samples (S3 Table), genomic DNA was directly amplified using the Repli-G kit (Qiagen), and then sequenced with Illumina HiSeq from TruSeq gDNA libraries by GenomeQuebec. One pair of *C. remanei* individuals derived from a shared isopod host (NS50-1, NS50-2), whereas all other individuals were isolated from different isopods.

To compare the population diversity patterns of this outcrossing species with a selfing species using similar approaches, we re-analyzed genomic sequences of 28 wild isolates of *C. elegans* collected at one location on the Big Island, Hawaii, from Crombie et al. [83]. For more details on the *C. elegans* sample, see Source data 1 from [83]. Sample IDs and the NCBI Sequence Read Archive accession numbers for *C. elegans* and *C. remanei* are in S1 Table.

### Variant calling, diversity statistics, and demography

We filtered and mapped reads to the *C. elegans* genome (project PRJNA13758, from WormBase version WS245) or the *C. remanei* genome (GCA_010183535.1 from the NCBI database) as described for the genetic map parental strains above. Variants were filtered with the standard GATK hard, only diallelic loci were used in the analysis. Additionally, we masked some genomic regions, such as indels with 10bp flanking regions with GATK and BEDTools v.2.25 [194]; repetitive regions using the masked versions of genomes and a script to extract them from [195]; and regions with too low or high mappability (<5X or >100x coverage in half of the individuals), all masks were combined by BEDTools merge.

We estimated 12 population diversity statistics using diploS/HIC fvecVcf [196] with two minor modifications: allowing not to normalize statistics, and to have only one sub-window in a window (see diploSHIC_note.txt on the project GitHub), and, additionally, the β-statistic using BetaScan [197] for 100 kb windows in *C. elegans* and *C. remanei* samples. To estimate the β-statistic for empirical data, we first applied an imputation with beagle v.5.0 [198] and a data format conversion by glactools [199]. Nucleotide diversity within introns and exons was estimated using these features from corresponding genome annotations, VCFtools [200], and BEDTools. We compared nucleotide diversity between domains of recombination and different gene features using Cohen’s d from the lsr package [201] and the Fisher-Pitman Permutation Test (Z) from the coin package [202] in R [203].

Demographic history was inferred by SMC++ v1.15.1 [113] for 100 bootstrapped replicates of 8 individuals using data from chromosomes I, III, IV for *C. elegans* and I, II, III, V for *C. remanei*. Additionally, we performed this analysis for all chromosomes separately. The genomic regions with unfit mappability or repeats were masked as described above. We used the mutation rate of 2.3e-09 base substitution per generation [106], and, for *C. elegans*, we rescaled this mutation rate by 0.5 and, later, the obtained generation time by 2 due to selfing (as discussed in [204]). To compare estimates of the recent population sizes from SMC++, as well as with previous estimates within *C. elegans*, we also calculated effective population size from nucleotide diversity data using the Watterson estimator [205] assuming neutrality and complete selfing for *C. elegans* (see [32] and [206], Ne=N/2). The recent demographic histories based on the linkage disequilibrium information were inferred by GONE ([112]), using autosomes and recombination rate retrieved from the genetic maps of *C. remanei* and *C. elegans* (Table 1; and Table 1 from [65]); the population size was averaged by taking the geometric mean across 100 replicates. Generation time was scaled by 2, and population size by 0.5 for *C. elegans*, analogous to the approach outlined above.

The inbreeding coefficient (F_is_) along the genome was estimated for sites with a minor allele frequency of more than 0.05 using VCFtools v.0.1.15, BEDtools v.2.25, HTSlib v.1.6, plink v1.90b4.6 [207], and popStats v1.0.0 [208]. The sample structure was visualized in the two-dimension space using latent coordinates from popVAE [209]. The effective recombination rate along the genomes was inferred by ReLERNN [176], using one of the replicates of reconstructed demography from SMC++ and (x0.5) mutation and (x2) time scaling for the selfing species. Linkage disequilibrium within and across chromosomes and the LD decay were estimated with plink using all sites for *C. elegans* sample, and every 10^th^ site for *C. remanei* for all LD calculations except the fine-resolution LD where we used all sites.

Figures were plotted via R packages dichromat ([210]), ggplot2 [211], gridExtra [212], ggpubr ([213]), magick ([214]), and also boot ([215]), coin ([216]), lsr ([217]), reshape2 ([218]), scales ([219]) to estimate summary statistics. The scripts for variant calling and masking, diversity statistics estimation, demography reconstructions, populations structure, effective recombination inference, LD decay, and corresponding figures and statistics are in https://github.com/phillips-lab/CR_CE_popgen/diversity_stats/.

### Reconstruction of the ancestral states of *C. remanei*

We reconstructed the ancestral states of the *C. remanei* and *C. latens* reference genomes. We used genomes of four strains of C. remanei, PX506, PX356 (GCA_001643735.2), PB4641 (GCA_000149515.1), PX439 (GCA_002259225.1) and *C. latens* (GCA_002259235.1). Preliminary topology was obtained with progressiveMauve [220] with 1 Mb regions from each of 6 chromosomes. Next, the genomes were masked based on their mappability by GenMap [221], and aligned by Progressive Cactus [222] with the following species tree topology: ((((*C. remanei* PB4641, *C. remanei* PX356), *C. remanei* PX439), *C. remanei* PX506), *C. latens*). The ancestral states were re-estimated by ancestorsML tool in the HAL tools [223].

Then for each chromosome, we calculated the fractions of sites with ancestral states, GC content in the ancestor, various types of substitutions per 100 kb windows, and extracted ancestral states positions polymorphic in the *C. remanei* population by HAL tools, BEDtools, and bash. We used Relate [114] to infer the history of the *C. remanei* population based on the ancestral states and the recombination map described above. We also examined demographic tests of the signal of positive selection along the genome, adjusting reported p-values using the harmonic mean p-value within 3Mb windows (2 Mb overlap) from harmonicmeanp [224] R package. The time to the most recent common ancestor (TMRCA), the relative TMRCA half-life (RTH, following [225]), and terminal branch lengths were assessed by phangorn [226] and phytools [227] R packages from Relate generated trees. Statistical analysis and visualization was made with coin, data.table ([228]), dplyr ([229]), ggplot2, gridExtra, ggpubr, lsr, pals ([230]), and scales packages in R. All scripts for the ancestral reconstruction, demographic inference, and related analyses available at https://github.com/phillips-lab/CR_CE_popgen/ancestral/

### Evolutionary simulations

To understand how various factors affect the genomic landscape of diversity, we run forward-time individual-based evolutionary simulations in SLiM v.3.3 ([8], [9]) using the tree-sequence format ([10]). We performed three types of simulation: 1) effects of the selfing rate, selection, and mutation landscape; 2) the decay of the ancestral diversity; 3) effects of rapid changes in population size on estimated statistics. All simulations had a population size of 5,000 individuals, 1 chromosome of 3 Mb in size with three recombination domains: left and right arms (1 Mb) with a high recombination rate (2.5e-7), and a 1 Mb central domain of low recombination (1e-9) to mimic the recombination domains of lower and higher recombination in nematodes.

In the first type of simulations, we changed selfing, mutation landscape, and selection regime. The selfing rate was 0% (outcrossers), 90%, 98%, 99.9%, or 100% (selfers). The mutation landscape was uniform with the mutation rate of 2e-8 in all domains of recombination. While this is an order of magnitude higher than actual mutation rate estimates, higher rates greatly facilitate the simulation process and, since the emphasis is on the relative values of mutation on arms and centers, this difference should not affect the normalized statistics used above. We used several domain-specific patterns of mutation rate differences: a uniform landscape with no difference in the mutation rate on the arms and the central domain (denoted as 1-1-1), 15% more mutations on the arms than on the centers (1.15-1-1.15), 50% more (1.5-1-1.5), or 2 times more mutations on the arms (2-1-2), the 1x mutation rate was the same in all simulations. Slightly elevated mutation rates within regions of higher recombination have been shown for worms previously [32], [49]. We considered four main types of selection regimes: only neutral mutations; neutral and 10% deleterious mutations; neutral, 10% deleterious, and 1% beneficial mutations; neutral, 10% deleterious, and 1% balancing mutations. Non-neutral simulations utilized various distributions for selection, and simulations with balancing selection used only one of these distributions (see details in S4 Table). Distributions of dominance were different for deleterious and beneficial mutations, with a shift towards recessive for the former and more additive coefficients for the latter (see S20 Fig and the SLiM scripts). All simulations were run for 50,000 generations (10N, burn-in).

The second type of simulations was designed to explore the effects of a switch from ancestral outcrossing to selfing. We ran neutral and non-neutral simulations (see details in S4 Table) in which all populations were outcrossing during burn-in and then subsequently changed to either 98 or 100% selfing. To observe the decay of ancestral diversity we saved results every 5,000 generations for 30,000 (6N) generations and repeated each scenario 50 times.

The aim of the third type of simulations was to reveal the consequences of rapid change in population size on the diversity statistics used in our analyses. These simulations utilized only neutral mutations but allowed different selfing rates and mutational landscapes. First, we performed simulations with 100 generations of exponential growth of 3% following burn-in, generating a rapid population size increase from 5K to about 100K. Second, we investigated the effects of fluctuations in population size following burn-in, by setting the population size to 15,000 and then back to 5,000 for 10 generations and so on 5 iterations. For the third type of scenario with exponential growth or fluctuations of the population size, we compared diversity statistics for the same populations at the burn-in generation and at the end of the simulation for 50 replicates.

We added neutral mutations, “recapitated” trees, and converted tree sequences from SLiM simulations to the VCF file format with 100 individuals using tskit v.0.3.4 ([231]), msprime v.0.6.1 ([232]), pyslim ([233]), and estimated diversity statistics with diploS/HIC and BetaScan as for empirical data but for 40kb windows with and without normalization. We also estimated the divergence from the ancestral genome using bash and BEDtools, F_is_ statistics via popStats, and calculated tree heights (TMRCA) from tree-sequences using python modules argparse ([234]), msprime, statistics ([239]), and pyslim.

We plotted and analyzed simulation results using packages coin, corrplot, dichromat, dplyr, ggplot2, gridExtra, ggpubr, and lsr packages in R. SLiM scripts for simulations, and related R scripts for visualization and analysis are in https://github.com/phillips-lab/CR_CE_popgen/simulations/.

### Classification of empirical data

In order to classify mutation rate, selfing rate, and type of selection on empirical data we trained a convolutional neural network (CNN) on a variety of features from the simulated chromosomes. Each simulated chromosome was divided into 75 sub-windows and 13 population diversity statistics were calculated within each sub-window, resulting in 13 by 75-pixel images used as input features for the CNN. The targets for the network consisted of one-hot encodings on each combination of the three evolutionary scenarios.

PCA and saliency plots were employed to visualize the similarity between simulated and empirical features. We projected the empirical features to PC space fitted to the simulated features in order to see how close the simulations resemble the empirical features. Saliency plots allow us to glimpse the features through the lens of the CNN. That is, we can see how sensitive the network classification is to perturbations of the summary statistics in each sub-window.

The CNN architecture was composed of four convolutional layers and four dense, fully connected layers. To prevent overfitting, dropout was used in between all but the final two layers. ReLU activations were used on all but the final layer, which had linear activation to improve numerical stability for the categorical cross-entropy loss functional. Weight learning was controlled by the Adam optimization routine with an initial learning rate of 1.5e-5. To reduce overfitting, we stopped training after the validation loss improved no more than 1e-7 for 100 epochs and saved the weights that resulted in the highest validation accuracy. The CNN was implemented using the Keras module within TensorFlow ([235]). The details of the analysis are at https://github.com/phillips-lab/CR_CE_popgen/cnn_model/.

## Supporting information

Captions for Supplementary figures

## Acknowledgments

We thank Rose Reynolds, Timothy Ahearne, Chadwick Smith, Scott Scholz for *C. remanei* mapping strain generation, Larry Meng for DNA isolation of *C. remanei* worms from the Koffler Scientific Reserve used for population genomic sequencing. Many thanks go to Sean O’rourke and members of Mike Miller’s lab at UC Davis for help with bestRAD sequencing, as well as to Gavin Woodruff, Murillo Rodrigues, Peter Ralph, Andy Kern, and other former and current members of Phillips lab and Kern-Ralph co-lab at the University of Oregon for comments and discussions. We thank the Research Advanced Computing Services team for assistance with the computing cluster Talapas at the University of Oregon and the Genomic Core Facility at the University of Oregon for assistance with library preparation.

## References

1. Charlesworth B, Charlesworth D. Population genetics from 1966 to 2016. Heredity. 2017;118: 2–9. doi:10.1038/hdy.2016.55

2. Crow JF, Kimura M. An Introduction to Population Genetics Theory. Blackburn Press; 2009.

3. Felsenstein J. Theoretical Evolutionary Genetics. University of Washington, Seattle; 1978. Available: evolution.gs.washington.edu/pgbook/

4. Ellegren H, Galtier N. Determinants of genetic diversity. Nat Rev Genet. 2016;17: 422–433. doi:10.1038/nrg.2016.58

5. Hoban S, Bertorelle G, Gaggiotti OE. Computer simulations: tools for population and evolutionary genetics. Nat Rev Genet. 2012;13: 110–122. doi:10.1038/nrg3130

6. Arenas M. Simulation of Molecular Data under Diverse Evolutionary Scenarios. Lewitter F, editor. PLoS Comput Biol. 2012;8: e1002495. doi:10.1371/journal.pcbi.1002495

7. Kelleher J, Etheridge AM, McVean G. Efficient Coalescent Simulation and Genealogical Analysis for Large Sample Sizes. PLOS Computational Biology. 2016;12: e1004842. doi:10.1371/journal.pcbi.1004842

8. Messer PW. SLiM: Simulating Evolution with Selection and Linkage. Genetics. 2013;194: 1037–1039. doi:10.1534/genetics.113.152181

9. Haller BC, Messer PW. SLiM 3: Forward Genetic Simulations Beyond the Wright–Fisher Model. Hernandez R, editor. Molecular Biology and Evolution. 2019;36: 632–637. doi:10.1093/molbev/msy228

10. Haller BC, Galloway J, Kelleher J, Messer PW, Ralph PL. Tree-sequence recording in SLiM opens new horizons for forward-time simulation of whole genomes. Molecular Ecology Resources. 2019;19: 552–566. doi:10.1111/1755-0998.12968

11. Sohn J, Nam J-W. The present and future of *de novo* whole-genome assembly. Brief Bioinform. 2016; bbw096. doi:10.1093/bib/bbw096

12. Jung H, Winefield C, Bombarely A, Prentis P, Waterhouse P. Tools and Strategies for Long-Read Sequencing and De Novo Assembly of Plant Genomes. Trends in Plant Science. 2019;24: 700–724. doi:10.1016/j.tplants.2019.05.003

13. Hotaling S, Kelley JL, Frandsen PB. Toward a genome sequence for every animal: Where are we now? Proc Natl Acad Sci USA. 2021;118: e2109019118. doi:10.1073/pnas.2109019118

14. Ségurel L, Wyman MJ, Przeworski M. Determinants of Mutation Rate Variation in the Human Germline. Annu Rev Genom Hum Genet. 2014;15: 47–70. doi:10.1146/annurev-genom-031714-125740

15. Campbell CR, Tiley GP, Poelstra JW, Hunnicutt KE, Larsen PA, Lee H-J, et al. Pedigree-based and phylogenetic methods support surprising patterns of mutation rate and spectrum in the gray mouse lemur. Heredity. 2021;127: 233–244. doi:10.1038/s41437-021-00446-5

16. Doitsidou M, Jarriault S, Poole RJ. Next-Generation Sequencing-Based Approaches for Mutation Mapping and Identification in *Caenorhabditis elegans*. Genetics. 2016;204: 451–474. doi:10.1534/genetics.115.186197

17. Peñalba JV, Wolf JBW. From molecules to populations: appreciating and estimating recombination rate variation. Nat Rev Genet. 2020;21: 476–492. doi:10.1038/s41576-020-0240-1

18. Zhou Y, Browning BL, Browning SR. Population-Specific Recombination Maps from Segments of Identity by Descent. The American Journal of Human Genetics. 2020;107: 137–148. doi:10.1016/j.ajhg.2020.05.016

19. Hinch AG, Zhang G, Becker PW, Moralli D, Hinch R, Davies B, et al. Factors influencing meiotic recombination revealed by whole-genome sequencing of single sperm. Science. 2019;363: eaau8861. doi:10.1126/science.aau8861

20. Dréau A, Venu V, Avdievich E, Gaspar L, Jones FC. Genome-wide recombination map construction from single individuals using linked-read sequencing. Nat Commun. 2019;10: 4309. doi:10.1038/s41467-019-12210-9

21. Santiago-Rodriguez TM, Hollister EB. Multi ‘omic data integration: A review of concepts, considerations, and approaches. Seminars in Perinatology. 2021;45: 151456. doi:10.1016/j.semperi.2021.151456

22. Krassowski M, Das V, Sahu SK, Misra BB. State of the Field in Multi-Omics Research: From Computational Needs to Data Mining and Sharing. Front Genet. 2020;11: 610798. doi:10.3389/fgene.2020.610798

23. Goodwin S, McPherson JD, McCombie WR. Coming of age: ten years of next-generation sequencing technologies. Nat Rev Genet. 2016;17: 333–351. doi:10.1038/nrg.2016.49

24. Hoban S, Kelley JL, Lotterhos KE, Antolin MF, Bradburd G, Lowry DB, et al. Finding the Genomic Basis of Local Adaptation: Pitfalls, Practical Solutions, and Future Directions. The American Naturalist. 2016;188: 379–397. doi:10.1086/688018

25. Min S, Lee B, Yoon S. Deep learning in bioinformatics. Brief Bioinform. 2016; bbw068. doi:10.1093/bib/bbw068

26. Zhang Z, Zhao Y, Liao X, Shi W, Li K, Zou Q, et al. Deep learning in omics: a survey and guideline. Briefings in Functional Genomics. 2019;18: 41–57. doi:10.1093/bfgp/ely030

27. Ching T, Himmelstein DS, Beaulieu-Jones BK, Kalinin AA, Do BT, Way GP, et al. Opportunities and obstacles for deep learning in biology and medicine. Journal of The Royal Society Interface. 2018;15: 20170387. doi:10.1098/rsif.2017.0387

28. Piccialli F, Somma VD, Giampaolo F, Cuomo S, Fortino G. A survey on deep learning in medicine: Why, how and when? Information Fusion. 2021;66: 111–137. doi:10.1016/j.inffus.2020.09.006

29. Sheehan S, Song YS. Deep Learning for Population Genetic Inference. PLOS Computational Biology. 2016;12: e1004845. doi:10.1371/journal.pcbi.1004845

30. Schrider DR, Kern AD. Supervised Machine Learning for Population Genetics: A New Paradigm. Trends in Genetics. 2018;34: 301–312. doi:10.1016/j.tig.2017.12.005

31. Golding GB, Strobeck C. Linkage disequilibrium in a finite population that is partially selfing. Genetics. 1980;94: 777–789. doi:10.1093/genetics/94.3.777

32. Pollak E. On the Theory of Partially Inbreeding Finite Populations. I. Partial Selfing. Genetics. 1987;117: 353–360.

33. Nordborg M, Donnelly P. The Coalescent Process with Selfing. Genetics. 1997;146: 1185–1195.

34. Nordborg M. Structured Coalescent Processes on Different Time Scales. Genetics. 1997;146: 1501–1514.

35. Nordborg M. Linkage disequilibrium, gene trees and selfing: an ancestral recombination graph with partial self-fertilization. Genetics. 2000;154: 923–929.

36. Maynard Smith J, Haigh J. The hitch-hiking effect of a favourable gene. Genetics Research. 1974;23: 23–35. doi:10.1017/S0016672300014634

37. Barton NH. Genetic hitchhiking. Philos Trans R Soc Lond B Biol Sci. 2000;355: 1553–1562. doi:10.1098/rstb.2000.0716

38. Charlesworth B, Morgan MT, Charlesworth D. The effect of deleterious mutations on neutral molecular variation. Genetics. 1993;134: 1289–1303. doi:10.1093/genetics/134.4.1289

39. Charlesworth B. Evolutionary rates in partially self-fertilizing species. Am Nat. 1992;140: 126–148. doi:10.1086/285406

40. Glémin S. Extinction and fixation times with dominance and inbreeding. Theor Popul Biol. 2012;81: 310–316. doi:10.1016/j.tpb.2012.02.006

41. Hartfield M, Wright SI, Agrawal AF. Coalescence and Linkage Disequilibrium in Facultatively Sexual Diploids. Genetics. 2018;210: 683–701. doi:10.1534/genetics.118.301244

42. Hartfield M, Glémin S. Limits to Adaptation in Partially Selfing Species. Genetics. 2016;203: 959–974. doi:10.1534/genetics.116.188821

43. Hartfield M, Bataillon T, Glémin S. The Evolutionary Interplay between Adaptation and Self-Fertilization. Trends Genet. 2017;33: 420–431. doi:10.1016/j.tig.2017.04.002

44. Thomas CG, Woodruff GC, Haag ES. Causes and consequences of the evolution of reproductive mode in Caenorhabditis nematodes. Trends in Genetics. 2012;28: 213–220. doi:10.1016/j.tig.2012.02.007

45. Cutter AD. Reproductive transitions in plants and animals: selfing syndrome, sexual selection and speciation. New Phytologist. 2019;224: 1080–1094. doi:10.1111/nph.16075

46. Félix M-A, Braendle C, Cutter AD. A Streamlined System for Species Diagnosis in Caenorhabditis (Nematoda: Rhabditidae) with Name Designations for 15 Distinct Biological Species. PLOS ONE. 2014;9: e94723. doi:10.1371/journal.pone.0094723

47. Cutter AD, Wasmuth JD, Washington NL. Patterns of Molecular Evolution in Caenorhabditis Preclude Ancient Origins of Selfing. Genetics. 2008;178: 2093–2104. doi:10.1534/genetics.107.085787

48. Kiontke KC, Félix M-A, Ailion M, Rockman MV, Braendle C, Pénigault J-B, et al. A phylogeny and molecular barcodes for Caenorhabditis, with numerous new species from rotting fruits. BMC Evolutionary Biology. 2011;11: 339. doi:10.1186/1471-2148-11-339

49. Brenner S. The Genetics of Caenorhabditis elegans. Genetics. 1974;77: 71–94.

50. Hodgkin J, Doniach T. Natural Variation and Copulatory Plug Formation in Caenorhabditis Elegans. Genetics. 1997;146: 149–164.

51. Teotónio H, Estes S, Phillips PC, Baer CF. Experimental Evolution with Caenorhabditis Nematodes. Genetics. 2017;206: 691–716. doi:10.1534/genetics.115.186288

52. Brenner S. The genetics of behaviour. Br Med Bull. 1973;29: 269–271. doi:10.1093/oxfordjournals.bmb.a071019

53. Wolinsky E, Way J. The behavioral genetics of Caenorhabditis elegans. Behav Genet. 1990;20: 169–189. doi:10.1007/BF01067789

54. Albrecht DR, Bargmann CI. High-content behavioral analysis of Caenorhabditis elegans in precise spatiotemporal chemical environments. Nat Methods. 2011;8: 599–605. doi:10.1038/nmeth.1630

55. Kwon N, Hwang AB, You Y-J, V. Lee S-J, Ho Je J. Dissection of C. elegans behavioral genetics in 3-D environments. Sci Rep. 2015;5: 9564. doi:10.1038/srep09564

56. The Nematode Caenorhabditis elegans. Wood W B. New York: Cold Spring Harbor Laboratory; 1988.

57. Corsi AK, Wightman B, Chalfie M. A Transparent Window into Biology: A Primer on Caenorhabditis elegans. Genetics. 2015;200: 387–407. doi:10.1534/genetics.115.176099

58. Meneely PM, Dahlberg CL, Rose JK. Working with Worms: Caenorhabditis elegans as a Model Organism. Current Protocols Essential Laboratory Techniques. 2019;19: e35. doi:10.1002/cpet.35

59. Gray JC, Cutter AD. Mainstreaming Caenorhabditis elegans in experimental evolution. Proc Biol Sci. 2014;281: 20133055. doi:10.1098/rspb.2013.3055

60. Nigon VM, Félix M-A. History of research on C. elegans and other free-living nematodes as model organisms. WormBook; 2018. Available: https://www.ncbi.nlm.nih.gov/books/NBK453431/

61. Andersen EC, Rockman MV. Natural genetic variation as a tool for discovery in *Caenorhabditis* nematodes. Félix M-A, editor. Genetics. 2022;220: iyab156. doi:10.1093/genetics/iyab156

62. Barrière A, Félix M-A. Temporal Dynamics and Linkage Disequilibrium in Natural *Caenorhabditis elegans* Populations. Genetics. 2007;176: 999–1011. doi:10.1534/genetics.106.067223

63. Cutter AD. Nucleotide Polymorphism and Linkage Disequilibrium in Wild Populations of the Partial Selfer *Caenorhabditis elegans*. Genetics. 2006;172: 171–184. doi:10.1534/genetics.105.048207

64. Cutter AD, Dey A, Murray RL. Evolution of the Caenorhabditis elegans Genome. Molecular Biology and Evolution. 2009;26: 1199–1234. doi:10.1093/molbev/msp048

65. Rockman MV, Kruglyak L. Recombinational Landscape and Population Genomics of Caenorhabditis elegans. PLOS Genetics. 2009;5: e1000419. doi:10.1371/journal.pgen.1000419

66. Andersen EC, Gerke JP, Shapiro JA, Crissman JR, Ghosh R, Bloom JS, et al. Chromosome-scale selective sweeps shape Caenorhabditis elegans genomic diversity. Nat Genet. 2012;44: 285–290. doi:10.1038/ng.1050

67. Félix M-A, Duveau F. Population dynamics and habitat sharing of natural populations of Caenorhabditis elegans and C. briggsae. BMC Biol. 2012;10: 59. doi:10.1186/1741-7007-10-59

68. Richaud A, Zhang G, Lee D, Lee J, Félix M-A. The Local Coexistence Pattern of Selfing Genotypes in *Caenorhabditis elegans* Natural Metapopulations. Genetics. 2018;208: 807–821. doi:10.1534/genetics.117.300564

69. Anderson JL, Morran LT, Phillips PC. Outcrossing and the Maintenance of Males within C. elegans Populations. J Hered. 2010;101: S62–S74. doi:10.1093/jhered/esq003

70. Teotonio H, Carvalho S, Manoel D, Roque M, Chelo IM. Evolution of Outcrossing in Experimental Populations of Caenorhabditis elegans. PLOS ONE. 2012;7: e35811. doi:10.1371/journal.pone.0035811

71. Carvalho S, Phillips PC, Teotónio H. Hermaphrodite life history and the maintenance of partial selfing in experimental populations of Caenorhabditis elegans. BMC Evolutionary Biology. 2014;14: 117. doi:10.1186/1471-2148-14-117

72. Cutter AD, Morran LT, Phillips PC. Males, Outcrossing, and Sexual Selection in *Caenorhabditis* Nematodes. Genetics. 2019;213: 27–57. doi:10.1534/genetics.119.300244

73. Thomas CG, Wang W, Jovelin R, Ghosh R, Lomasko T, Trinh Q, et al. Full-genome evolutionary histories of selfing, splitting, and selection in Caenorhabditis. Genome Res. 2015;25: 667–678. doi:10.1101/gr.187237.114

74. Noble LM, Yuen J, Stevens L, Moya N, Persaud R, Moscatelli M, et al. Selfing is the safest sex for Caenorhabditis tropicalis. Castric V, Weigel D, Castric V, editors. eLife. 2021;10: e62587. doi:10.7554/eLife.62587

75. Snoek BL, Volkers RJM, Nijveen H, Petersen C, Dirksen P, Sterken MG, et al. A multi-parent recombinant inbred line population of C. elegans allows identification of novel QTLs for complex life history traits. BMC Biology. 2019;17: 24. doi:10.1186/s12915-019-0642-8

76. Ross JA, Koboldt DC, Staisch JE, Chamberlin HM, Gupta BP, Miller RD, et al. Caenorhabditis briggsae Recombinant Inbred Line Genotypes Reveal Inter-Strain Incompatibility and the Evolution of Recombination. PLoS Genet. 2011;7: e1002174. doi:10.1371/journal.pgen.1002174

77. Teterina AA, Willis JH, Phillips PC. Chromosome-Level Assembly of the Caenorhabditis remanei Genome Reveals Conserved Patterns of Nematode Genome Organization. Genetics. 2020;214: 769–780. doi:10.1534/genetics.119.303018

78. Jovelin R, Ajie BC, Phillips PC. Molecular evolution and quantitative variation for chemosensory behaviour in the nematode genus Caenorhabditis. Mol Ecol. 2003;12: 1325–1337. doi:10.1046/j.1365-294x.2003.01805.x

79. Cutter AD, Charlesworth B. Selection Intensity on Preferred Codons Correlates with Overall Codon Usage Bias in Caenorhabditis remanei. Current Biology. 2006;16: 2053–2057. doi:10.1016/j.cub.2006.08.067

80. Cutter AD, Baird SE, Charlesworth D. High Nucleotide Polymorphism and Rapid Decay of Linkage Disequilibrium in Wild Populations of Caenorhabditis remanei. Genetics. 2006;174: 901–913. doi:10.1534/genetics.106.061879

81. Jovelin R, Dunham JP, Sung FS, Phillips PC. High Nucleotide Divergence in Developmental Regulatory Genes Contrasts With the Structural Elements of Olfactory Pathways in Caenorhabditis. Genetics. 2009;181: 1387–1397. doi:10.1534/genetics.107.082651

82. Dey A, Jeon Y, Wang G-X, Cutter AD. Global Population Genetic Structure of Caenorhabditis remanei Reveals Incipient Speciation. Genetics. 2012;191: 1257–1269. doi:10.1534/genetics.112.140418

83. Crombie TA, Zdraljevic S, Cook DE, Tanny RE, Brady SC, Wang Y, et al. Deep sampling of Hawaiian Caenorhabditis elegans reveals high genetic diversity and admixture with global populations. eLife. 2019;8: e50465. doi:10.7554/eLife.50465

84. Wagner CR, Kuervers L, Baillie D, Yanowitz JL. xnd-1 Regulates the Global Recombination Landscape in C. elegans. Nature. 2010;467: 839–843. doi:10.1038/nature09429

85. Meneely PM, McGovern OL, Heinis FI, Yanowitz JL. Crossover Distribution and Frequency Are Regulated by him-5 in Caenorhabditis elegans. Genetics. 2012;190: 1251–1266. doi:10.1534/genetics.111.137463

86. Yu Z, Kim Y, Dernburg AF. Meiotic Recombination and the Crossover Assurance Checkpoint in Caenorhabditis elegans. Semin Cell Dev Biol. 2016;54: 106–116. doi:10.1016/j.semcdb.2016.03.014

87. Zetka MC, Rose AM. Sex-Related Differences in Crossing over in Caenorhabditis Elegans. Genetics. 1990;126: 355–363.

88. Meneely PM, Farago AF, Kauffman TM. Crossover distribution and high interference for both the X chromosome and an autosome during oogenesis and spermatogenesis in Caenorhabditis elegans. Genetics. 2002;162: 1169–1177.

89. Lim JGY, Stine RRW, Yanowitz JL. Domain-Specific Regulation of Recombination in Caenorhabditis elegans in Response to Temperature, Age and Sex. Genetics. 2008;180: 715–726. doi:10.1534/genetics.108.090142

90. Saito TT, Colaiácovo MP. Regulation of Crossover Frequency and Distribution during Meiotic Recombination. Cold Spring Harb Symp Quant Biol. 2017;82: 223–234. doi:10.1101/sqb.2017.82.034132

91. Hillier LW, Miller RD, Baird SE, Chinwalla A, Fulton LA, Koboldt DC, et al. Comparison of C. elegans and C. briggsae Genome Sequences Reveals Extensive Conservation of Chromosome Organization and Synteny. PLOS Biology. 2007;5: e167. doi:10.1371/journal.pbio.0050167

92. Graustein A, Gaspar JM, Walters JR, Palopoli MF. Levels of DNA Polymorphism Vary With Mating System in the Nematode Genus Caenorhabditis. Genetics. 2002;161: 99–107.

93. Sivasundar A, Hey J. Population Genetics of Caenorhabditis elegans: The Paradox of Low Polymorphism in a Widespread Species. Genetics. 2003;163: 147–157.

94. Barrière A, Félix M-A. High local genetic diversity and low outcrossing rate in Caenorhabditis elegans natural populations. Curr Biol. 2005;15: 1176–1184. doi:10.1016/j.cub.2005.06.022

95. Haber M, Schüngel M, Putz A, Müller S, Hasert B, Schulenburg H. Evolutionary history of Caenorhabditis elegans inferred from microsatellites: evidence for spatial and temporal genetic differentiation and the occurrence of outbreeding. Mol Biol Evol. 2005;22: 160–173. doi:10.1093/molbev/msh264

96. Haag ES, Ackerman AD. Intraspecific variation in fem-3 and tra-2, two rapidly coevolving nematode sex-determining genes. Gene. 2005;349: 35–42. doi:10.1016/j.gene.2004.12.051

97. Dolgin ES, Charlesworth B, Baird SE, Cutter AD. Inbreeding and Outbreeding Depression in Caenorhabditis Nematodes. Evolution. 2007;61: 1339–1352. doi:10.1111/j.1558-5646.2007.00118.x

98. Lee D, Zdraljevic S, Stevens L, Wang Y, Tanny RE, Crombie TA, et al. Balancing selection maintains hyper-divergent haplotypes in Caenorhabditis elegans. Nat Ecol Evol. 2021;5: 794–807. doi:10.1038/s41559-021-01435-x

99. Siewert KM, Voight BF. BetaScan2: Standardized Statistics to Detect Balancing Selection Utilizing Substitution Data. Genome Biology and Evolution. 2020;12: 3873–3877. doi:10.1093/gbe/evaa013

100. Fierst JL, Willis JH, Thomas CG, Wang W, Reynolds RM, Ahearne TE, et al. Reproductive Mode and the Evolution of Genome Size and Structure in Caenorhabditis Nematodes. PLOS Genetics. 2015;11: e1005323. doi:10.1371/journal.pgen.1005323

101. Tamura K. Estimation of the number of nucleotide substitutions when there are strong transition-transversion and G+C-content biases. Mol Biol Evol. 1992;9: 678–687. doi:10.1093/oxfordjournals.molbev.a040752

102. Phung TN, Huber CD, Lohmueller KE. Determining the Effect of Natural Selection on Linked Neutral Divergence across Species. Akey JM, editor. PLoS Genet. 2016;12: e1006199. doi:10.1371/journal.pgen.1006199

103. Konrad A, Brady MJ, Bergthorsson U, Katju V. Mutational Landscape of Spontaneous Base Substitutions and Small Indels in Experimental Caenorhabditis elegans Populations of Differing Size. Genetics. 2019;212: 837–854. doi:10.1534/genetics.119.302054

104. Denver DR, Dolan PC, Wilhelm LJ, Sung W, Lucas-Lledó JI, Howe DK, et al. A genome-wide view of Caenorhabditis elegans base-substitution mutation processes. Proc Natl Acad Sci U S A. 2009;106: 16310–16314. doi:10.1073/pnas.0904895106

105. Chu JS-C, Johnsen RC, Chua SY, Tu D, Dennison M, Marra M, et al. Allelic Ratios and the Mutational Landscape Reveal Biologically Significant Heterozygous SNVs. Genetics. 2012;190: 1225–1233. doi:10.1534/genetics.111.137208

106. Saxena AS, Salomon MP, Matsuba C, Yeh S-D, Baer CF. Evolution of the Mutational Process under Relaxed Selection in Caenorhabditis elegans. Mol Biol Evol. 2019;36: 239–251. doi:10.1093/molbev/msy213

107. Cutter AD, Payseur BA. Genomic signatures of selection at linked sites: unifying the disparity among species. Nat Rev Genet. 2013;14: 262–274. doi:10.1038/nrg3425

108. Spencer CCA, Deloukas P, Hunt S, Mullikin J, Myers S, Silverman B, et al. The Influence of Recombination on Human Genetic Diversity. PLOS Genetics. 2006;2: e148. doi:10.1371/journal.pgen.0020148

109. Halldorsson BV, Palsson G, Stefansson OA, Jonsson H, Hardarson MT, Eggertsson HP, et al. Characterizing mutagenic effects of recombination through a sequence-level genetic map. Science. 2019;363. doi:10.1126/science.aau1043

110. David P, Pujol B, Viard F, Castella V, Goudet J. Reliable selfing rate estimates from imperfect population genetic data. Mol Ecol. 2007;16: 2474–2487. doi:10.1111/j.1365-294X.2007.03330.x

111. Sivasundar A, Hey J. Sampling from natural populations with RNAI reveals high outcrossing and population structure in Caenorhabditis elegans. Curr Biol. 2005;15: 1598–1602. doi:10.1016/j.cub.2005.08.034

112. Santiago E, Novo I, Pardiñas AF, Saura M, Wang J, Caballero A. Recent Demographic History Inferred by High-Resolution Analysis of Linkage Disequilibrium. Kim Y, editor. Molecular Biology and Evolution. 2020;37: 3642–3653. doi:10.1093/molbev/msaa169

113. Terhorst J, Kamm JA, Song YS. Robust and scalable inference of population history from hundreds of unphased whole genomes. Nature Genetics. 2017;49: 303–309. doi:10.1038/ng.3748

114. Speidel L, Forest M, Shi S, Myers SR. A method for genome-wide genealogy estimation for thousands of samples. Nature Genetics. 2019;51: 1321–1329. doi:10.1038/s41588-019-0484-x

115. Wright S. The genetical structure of populations. Ann Eugen. 1951;15: 323–354. doi:10.1111/j.1469-1809.1949.tb02451.x

116. Hudson RR. Properties of a neutral allele model with intragenic recombination. Theoretical Population Biology. 1983;23: 183–201. doi:10.1016/0040-5809(83)90013-8

117. Charlesworth D, Wright SI. Breeding systems and genome evolution. Curr Opin Genet Dev. 2001;11: 685–690. doi:10.1016/s0959-437x(00)00254-9

118. Bengtsson BO. Genetic variation in organisms with sexual and asexual reproduction. J Evol Biol. 2003;16: 189–199. doi:10.1046/j.1420-9101.2003.00523.x

119. Ceplitis A. Coalescence times and the Meselson effect in asexual eukaryotes. Genet Res. 2003;82: 183–190. doi:10.1017/s0016672303006487

120. Roze D. Background Selection in Partially Selfing Populations. Genetics. 2016;203: 937–957. doi:10.1534/genetics.116.187955

121. Agrawal AF, Hartfield M. Coalescence with Background and Balancing Selection in Systems with Bi- and Uniparental Reproduction: Contrasting Partial Asexuality and Selfing. Genetics. 2016;202: 313–326. doi:10.1534/genetics.115.181024

122. Hartfield M. Evolutionary genetic consequences of facultative sex and outcrossing. J Evol Biol. 2016;29: 5–22. doi:10.1111/jeb.12770

123. Hudson, Richard R. Gene genealogies and the coalescent process. Oxford surveys in evolutionary biology. 1990;7: 44.

124. Frézal L, Félix M-A. C. elegans outside the Petri dish. eLife. 2015;4: e05849. doi:10.7554/eLife.05849

125. Seidel HS, Rockman MV, Kruglyak L. Widespread Genetic Incompatibility in *C. Elegans* Maintained by Balancing Selection. Science. 2008;319: 589–594. doi:10.1126/science.1151107

126. Mark Welch DB, Meselson M. Evidence for the Evolution of Bdelloid Rotifers Without Sexual Reproduction or Genetic Exchange. Science. 2000;288: 1211–1215. doi:10.1126/science.288.5469.1211

127. Hartfield M, Wright SI, Agrawal AF. Coalescent Times and Patterns of Genetic Diversity in Species with Facultative Sex: Effects of Gene Conversion, Population Structure, and Heterogeneity. Genetics. 2016;202: 297–312. doi:10.1534/genetics.115.178004

128. McVean GAT. A Genealogical Interpretation of Linkage Disequilibrium. Genetics. 2002;162: 987–991. doi:10.1093/genetics/162.2.987

129. Gao J, Kim H-M, Elia AE, Elledge SJ, Colaiácovo MP. NatB Domain-Containing CRA-1 Antagonizes Hydrolase ACER-1 Linking Acetyl-CoA Metabolism to the Initiation of Recombination during C. elegans Meiosis. PLOS Genetics. 2015;11: e1005029. doi:10.1371/journal.pgen.1005029

130. McClendon TB, Mainpal R, Amrit FRG, Krause MW, Ghazi A, Yanowitz JL. X Chromosome Crossover Formation and Genome Stability in *Caenorhabditis elegans* Are Independently Regulated by *xnd-1*. G3 Genes|Genomes|Genetics. 2016;6: 3913–3925. doi:10.1534/g3.116.035725

131. Jaramillo-Lambert A, Harigaya Y, Vitt J, Villeneuve A, Engebrecht J. Meiotic Errors Activate Checkpoints that Improve Gamete Quality without Triggering Apoptosis in Male Germ Cells. Curr Biol. 2010;20: 2078–2089. doi:10.1016/j.cub.2010.10.008

132. The C. elegans Sequencing Consortium*. Genome Sequence of the Nematode *C. elegans* : A Platform for Investigating Biology. Science. 1998;282: 2012–2018. doi:10.1126/science.282.5396.2012

133. Woodruff GC, Teterina AA. Degradation of the Repetitive Genomic Landscape in a Close Relative of Caenorhabditis elegans. Lu J, editor. Molecular Biology and Evolution. 2020;37: 2549–2567. doi:10.1093/molbev/msaa107

134. Charlesworth D. Effects of inbreeding on the genetic diversity of populations. Philos Trans R Soc Lond B Biol Sci. 2003;358: 1051–1070. doi:10.1098/rstb.2003.1296

135. Glémin S, Bazin E, Charlesworth D. Impact of mating systems on patterns of sequence polymorphism in flowering plants. Proceedings of the Royal Society B: Biological Sciences. 2006;273: 3011–3019. doi:10.1098/rspb.2006.3657

136. Phillips PC. Self fertilization sweeps up variation in the worm genome. Nat Genet. 2012;44: 237–238. doi:10.1038/ng.2201

137. Glémin S, François CM, Galtier N. Genome Evolution in Outcrossing vs. Selfing vs. Asexual Species. In: Anisimova M, editor. Evolutionary Genomics: Statistical and Computational Methods. New York, NY: Springer; 2019. pp. 331–369. doi:10.1007/978-1-4939-9074-0_11

138. Kaplan NL, Hudson RR, Langley CH. The “hitchhiking effect” revisited. Genetics. 1989;123: 887–899. doi:10.1093/genetics/123.4.887

139. Stephan W, Wiehe THE, Lenz MW. The effect of strongly selected substitutions on neutral polymorphism: Analytical results based on diffusion theory. Theoretical Population Biology. 1992;41: 237–254. doi:10.1016/0040-5809(92)90045-U

140. Nordborg M, Charlesworth B, Charlesworth D. The effect of recombination on background selection. Genet Res. 1996;67: 159–174. doi:10.1017/s0016672300033619

141. Coop G, Ralph P. Patterns of Neutral Diversity Under General Models of Selective Sweeps. Genetics. 2012;192: 205–224. doi:10.1534/genetics.112.141861

142. Aguade M, Miyashita N, Langley CH. Reduced variation in the yellow-achaete-scute region in natural populations of Drosophila melanogaster. Genetics. 1989;122: 607–615. doi:10.1093/genetics/122.3.607

143. Begun DJ, Aquadro CF. Levels of naturally occurring DNA polymorphism correlate with recombination rates in D. melanogaster. Nature. 1992;356: 519–520. doi:10.1038/356519a0

144. Cutter AD, Payseur BA. Selection at Linked Sites in the Partial Selfer Caenorhabditis elegans. Molecular Biology and Evolution. 2003;20: 665–673. doi:10.1093/molbev/msg072

145. Nordborg M, Hu TT, Ishino Y, Jhaveri J, Toomajian C, Zheng H, et al. The Pattern of Polymorphism in Arabidopsis thaliana. PLOS Biology. 2005;3: e196. doi:10.1371/journal.pbio.0030196

146. Begun DJ, Holloway AK, Stevens K, Hillier LW, Poh Y-P, Hahn MW, et al. Population Genomics: Whole-Genome Analysis of Polymorphism and Divergence in Drosophila simulans. PLOS Biology. 2007;5: e310. doi:10.1371/journal.pbio.0050310

147. Cutter AD, Choi JY. Natural selection shapes nucleotide polymorphism across the genome of the nematode Caenorhabditis briggsae. Genome Res. 2010;20: 1103–1111. doi:10.1101/gr.104331.109

148. Cutter AD, Moses AM. Polymorphism, divergence, and the role of recombination in Saccharomyces cerevisiae genome evolution. Mol Biol Evol. 2011;28: 1745–1754. doi:10.1093/molbev/msq356

149. Stankowski S, Chase MA, Fuiten AM, Rodrigues MF, Ralph PL, Streisfeld MA. Widespread selection and gene flow shape the genomic landscape during a radiation of monkeyflowers. PLOS Biology. 2019;17: e3000391. doi:10.1371/journal.pbio.3000391

150. Coop G. Does linked selection explain the narrow range of genetic diversity across species? Evolutionary Biology; 2016 Mar. doi:10.1101/042598

151. Haldane. A mathematical theory of natural and artificial selection, part V: selection and mutation. Mathematical Proceedings of the Cambridge Philosophical Society. 23: 838--844.

152. Stebbins GL. Self Fertilization and Population Variability in the Higher Plants. The American Naturalist. 1957;91: 337–354. doi:10.1086/281999

153. Rockman MV, Skrovanek SS, Kruglyak L. Selection at Linked Sites Shapes Heritable Phenotypic Variation in *C. elegans*. Science. 2010;330: 372–376. doi:10.1126/science.1194208

154. Cutter AD. Molecular evolution inferences from the C. elegans genome. WormBook; 2018. Available: https://www.ncbi.nlm.nih.gov/books/NBK116079/

155. Wolfe KH, Sharp PM, Li W-H. Mutation rates differ among regions of the mammalian genome. Nature. 1989;337: 283–285. doi:10.1038/337283a0

156. Lercher MJ, Hurst LD. Human SNP variability and mutation rate are higher in regions of high recombination. Trends in Genetics. 2002;18: 337–340. doi:10.1016/S0168-9525(02)02669-0

157. Stamatoyannopoulos JA, Adzhubei I, Thurman RE, Kryukov GV, Mirkin SM, Sunyaev SR. Human mutation rate associated with DNA replication timing. Nat Genet. 2009;41: 393–395. doi:10.1038/ng.363

158. Ananda G, Chiaromonte F, Makova KD. A genome-wide view of mutation rate co-variation using multivariate analyses. Genome Biol. 2011;12: R27. doi:10.1186/gb-2011-12-3-r27

159. Schuster-Böckler B, Lehner B. Chromatin organization is a major influence on regional mutation rates in human cancer cells. Nature. 2012;488: 504–507. doi:10.1038/nature11273

160. Woo YH, Li W-H. DNA replication timing and selection shape the landscape of nucleotide variation in cancer genomes. Nat Commun. 2012;3: 1004. doi:10.1038/ncomms1982

161. Arbeithuber B, Betancourt AJ, Ebner T, Tiemann-Boege I. Crossovers are associated with mutation and biased gene conversion at recombination hotspots. Proc Natl Acad Sci USA. 2015;112: 2109–2114. doi:10.1073/pnas.1416622112

162. Polak P, Karlić R, Koren A, Thurman R, Sandstrom R, Lawrence M, et al. Cell-of-origin chromatin organization shapes the mutational landscape of cancer. Nature. 2015;518: 360–364. doi:10.1038/nature14221

163. Supek F, Lehner B. Differential DNA mismatch repair underlies mutation rate variation across the human genome. Nature. 2015;521: 81–84. doi:10.1038/nature14173

164. Terekhanova NV, Seplyarskiy VB, Soldatov RA, Bazykin GA. Evolution of Local Mutation Rate and Its Determinants. Mol Biol Evol. 2017;34: 1100–1109. doi:10.1093/molbev/msx060

165. Gaffney DJ, Keightley PD. The scale of mutational variation in the murid genome. Genome Res. 2005;15: 1086–1094. doi:10.1101/gr.3895005

166. Gonzalez-Perez A, Sabarinathan R, Lopez-Bigas N. Local Determinants of the Mutational Landscape of the Human Genome. Cell. 2019;177: 101–114. doi:10.1016/j.cell.2019.02.051

167. Serero A, Jubin C, Loeillet S, Legoix-Né P, Nicolas AG. Mutational landscape of yeast mutator strains. Proc Natl Acad Sci U S A. 2014;111: 1897–1902. doi:10.1073/pnas.1314423111

168. Monroe JG, Srikant T, Carbonell-Bejerano P, Becker C, Lensink M, Exposito-Alonso M, et al. Mutation bias reflects natural selection in Arabidopsis thaliana. Nature. 2022;602: 101–105. doi:10.1038/s41586-021-04269-6

169. Rajaei M, Saxena AS, Johnson LM, Snyder MC, Crombie TA, Tanny RE, et al. Mutability of mononucleotide repeats, not oxidative stress, explains the discrepancy between laboratory-accumulated mutations and the natural allele-frequency spectrum in C. elegans. Genome Res. 2021;31: 1602–1613. doi:10.1101/gr.275372.121

170. Katju V, Konrad A, Deiss TC, Bergthorsson U. Mutation rate and spectrum in obligately outcrossing *Caenorhabditis elegans* mutation accumulation lines subjected to RNAi-induced knockdown of the mismatch repair gene *msh-2*. Félix M-A, editor. G3 Genes|Genomes|Genetics. 2022;12: jkab364. doi:10.1093/g3journal/jkab364

171. Cutter AD. *Caenorhabditis* evolution in the wild. BioEssays. 2015;37: 983–995. doi:10.1002/bies.201500053

172. Hanski I. Metapopulation dynamics. Nature. 1998;396: 41–49. doi:10.1038/23876

173. Jovelin R, Comstock JS, Cutter AD, Phillips PC. A recent global selective sweep on the age-1 phosphatidylinositol 3-OH kinase regulator of the insulin-like signaling pathway within Caenorhabditis remanei. G3 (Bethesda). 2014;4: 1123–1133. doi:10.1534/g3.114.010629

174. Hartfield M, Glémin S. Hitchhiking of Deleterious Alleles and the Cost of Adaptation in Partially Selfing Species. Genetics. 2014;196: 281–293. doi:10.1534/genetics.113.158196

175. Glémin S. Balancing selection in self-fertilizing populations. Evolution. 2021;75: 1011–1029. doi:10.1111/evo.14194

176. Adrion JR, Galloway JG, Kern AD. Predicting the Landscape of Recombination Using Deep Learning. Molecular Biology and Evolution. 2020;37: 1790–1808. doi:10.1093/molbev/msaa038

177. Sikkink KL, Reynolds RM, Ituarte CM, Cresko WA, Phillips PC. Rapid evolution of phenotypic plasticity and shifting thresholds of genetic assimilation in the nematode Caenorhabditis remanei. G3: Genes, Genomes, Genetics. 2014;4: 1103–1112. doi:10.1534/g3.114.010553

178. Ali OA, O’Rourke SM, Amish SJ, Meek MH, Luikart G, Jeffres C, et al. RAD Capture (Rapture): Flexible and Efficient Sequence-Based Genotyping. Genetics. 2016;202: 389–400. doi:10.1534/genetics.115.183665

179. Andrews S. FastQC: a quality control tool for high throughput sequence data. In: Babraham Bioinformatics [Internet]. 2010 [cited 17 Nov 2020]. Available: https://www.bioinformatics.babraham.ac.uk/projects/fastqc/

180. Ewels P, Magnusson M, Lundin S, Käller M. MultiQC: summarize analysis results for multiple tools and samples in a single report. Bioinformatics. 2016;32: 3047–3048. doi:10.1093/bioinformatics/btw354

181. Jiang H, Lei R, Ding S-W, Zhu S. Skewer: a fast and accurate adapter trimmer for next-generation sequencing paired-end reads. BMC Bioinformatics. 2014;15: 182. doi:10.1186/1471-2105-15-182

182. Li H. Aligning sequence reads, clone sequences and assembly contigs with BWA-MEM. arXiv:13033997 [q-bio]. 2013 [cited 17 Nov 2020]. Available: http://arxiv.org/abs/1303.3997

183. Li H, Handsaker B, Wysoker A, Fennell T, Ruan J, Homer N, et al. The Sequence Alignment/Map format and SAMtools. Bioinformatics. 2009;25: 2078–2079. doi:10.1093/bioinformatics/btp352

184. Broad Institute. Picard Tools. [cited 17 Nov 2020]. Available: http://broadinstitute.github.io/picard/

185. McKenna A, Hanna M, Banks E, Sivachenko A, Cibulskis K, Kernytsky A, et al. The Genome Analysis Toolkit: A MapReduce framework for analyzing next-generation DNA sequencing data. Genome Res. 2010;20: 1297–1303. doi:10.1101/gr.107524.110

186. how to Apply hard filters to a call set - Legacy GATK Forum. [cited 17 Nov 2020]. Available: https://sites.google.com/a/broadinstitute.org/legacy-gatk-forum-discussions/tutorials/2806-how-to-apply-hard-filters-to-a-call-set

187. Van der Auwera GA, Carneiro MO, Hartl C, Poplin R, Angel G del, Levy-Moonshine A, et al. From FastQ Data to High-Confidence Variant Calls: The Genome Analysis Toolkit Best Practices Pipeline. Current Protocols in Bioinformatics. 2013;43: 11.10.1–11.10.33. doi:https://doi.org/10.1002/0471250953.bi1110s43

188. DePristo MA, Banks E, Poplin R, Garimella KV, Maguire JR, Hartl C, et al. A framework for variation discovery and genotyping using next-generation DNA sequencing data. Nature Genetics. 2011;43: 491–498. doi:10.1038/ng.806

189. Hether T. Flip2BeRAD: Python and C++ utilities for flipping RADseq reads. 2017. Available: https://github.com/tylerhether/Flip2BeRAD

190. Catchen J, Hohenlohe PA, Bassham S, Amores A, Cresko WA. Stacks: an analysis tool set for population genomics. Molecular Ecology. 2013;22: 3124–3140. doi:https://doi.org/10.1111/mec.12354

191. Rastas P. Lep-MAP3: robust linkage mapping even for low-coverage whole genome sequencing data. Bioinformatics. 2017;33: 3726–3732. doi:10.1093/bioinformatics/btx494

192. Muggeo VM. Segmented: an R package to fit regression models with broken-line relationships. R News 8 (1): 20–25. R Foundation for Statistical Computing Vienna, Austria; 2008.

193. Baird SE. Natural and experimental associations of Caenorhabditis remanei with Trachelipus rathkii and other terrestrial isopods. Nematology. 1999;1: 471–475. doi:10.1163/156854199508478

194. Quinlan AR, Hall IM. BEDTools: a flexible suite of utilities for comparing genomic features. Bioinformatics. 2010;26: 841–842. doi:10.1093/bioinformatics/btq033

195. Cook DE. Generates the masked ranges within a fasta file. In: Gist [Internet]. [cited 23 Nov 2020]. Available: https://gist.github.com/danielecook/cfaa5c359d99bcad3200

196. Kern AD, Schrider DR. diploS/HIC: An Updated Approach to Classifying Selective Sweeps. G3 (Bethesda). 2018;8: 1959–1970. doi:10.1534/g3.118.200262

197. Siewert KM, Voight BF. Detecting Long-Term Balancing Selection Using Allele Frequency Correlation. Mol Biol Evol. 2017;34: 2996–3005. doi:10.1093/molbev/msx209

198. Browning BL, Zhou Y, Browning SR. A One-Penny Imputed Genome from Next-Generation Reference Panels. The American Journal of Human Genetics. 2018;103: 338–348. doi:10.1016/j.ajhg.2018.07.015

199. Renaud G. glactools: a command-line toolset for the management of genotype likelihoods and allele counts. Bioinformatics. 2018;34: 1398–1400. doi:10.1093/bioinformatics/btx749

200. Danecek P, Auton A, Abecasis G, Albers CA, Banks E, DePristo MA, et al. The variant call format and VCFtools. Bioinformatics. 2011;27: 2156–2158. doi:10.1093/bioinformatics/btr330

201. Navarro D. Learning Statistics with R: a tutorial for psychology students and other beginners. 2015 [cited 23 Nov 2020]. Available: https://learningstatisticswithr.com

202. Hothorn T, Hornik K, Wiel MA van de, Zeileis A. Implementing a Class of Permutation Tests: The coin Package. Journal of Statistical Software. 2008;28: 1–23. doi:10.18637/jss.v028.i08

203. R Core Team. A language and environment for statistical computing R Foundation for Statistical Computing Department of Agronomy, Faculty of Agriculture of the University of the Free State. Vienna, Austria www R-project org. 2017.

204. Partially selfing organisms · Issue #82 · popgenmethods/smcpp. In: GitHub [Internet]. [cited 23 Nov 2020]. Available: https://github.com/popgenmethods/smcpp/issues/82

205. Watterson GA. On the number of segregating sites in genetical models without recombination. Theoretical Population Biology. 1975;7: 256–276. doi:10.1016/0040-5809(75)90020-9

206. Wang J, Santiago E, Caballero A. Prediction and estimation of effective population size. Heredity. 2016;117: 193–206. doi:10.1038/hdy.2016.43

207. Weeks JP. plink: An R Package for Linking Mixed-Format Tests Using IRT-Based Methods. Journal of Statistical Software. 2010;35: 1–33. doi:10.18637/jss.v035.i12

208. Skoglund P, Mallick S, Bortolini MC, Chennagiri N, Hünemeier T, Petzl-Erler ML, et al. Genetic evidence for two founding populations of the Americas. Nature. 2015;525: 104–108. doi:10.1038/nature14895

209. Battey CJ, Coffing GC, Kern AD. Visualizing population structure with variational autoencoders. G3 Genes|Genomes|Genetics. 2021;11. doi:10.1093/g3journal/jkaa036

210. Lumley T, Knoblauch K, Waichler S, Zeileis A. dichromat: Color Schemes for Dichromats. 2022. Available: https://CRAN.R-project.org/package=dichromat

211. Wickham H. ggplot2: elegant graphics for data analysis. Springer; 2016.

212. Auguie B, Antonov A. gridExtra: Miscellaneous Functions for “Grid” Graphics. 2017. Available: https://CRAN.R-project.org/package=gridExtra

213. Kassambara A. ggpubr: “ggplot2” Based Publication Ready Plots. 2020. Available: https://CRAN.R-project.org/package=ggpubr

214. Ooms J. magick: Advanced Graphics and Image-Processing in R. 2021. Available: https://CRAN.R-project.org/package=magick

215. Canty A, Ripley B. boot: Bootstrap Functions (Originally by Angelo Canty for S). 2021. Available: https://CRAN.R-project.org/package=boot

216. Hothorn T, Winell H, Hornik K, van de Wiel MA, Zeileis A. coin: Conditional Inference Procedures in a Permutation Test Framework. 2021. Available: https://CRAN.R-project.org/package=coin

217. Navarro D. lsr: Companion to “Learning Statistics with R.” 2021. Available: https://CRAN.R-project.org/package=lsr

218. Wickham H. reshape2: Flexibly Reshape Data: A Reboot of the Reshape Package. 2020. Available: https://CRAN.R-project.org/package=reshape2

219. Wickham H, Seidel D, RStudio. scales: Scale Functions for Visualization. 2022. Available: https://CRAN.R-project.org/package=scales

220. Darling AE, Mau B, Perna NT. progressiveMauve: Multiple Genome Alignment with Gene Gain, Loss and Rearrangement. PLOS ONE. 2010;5: e11147. doi:10.1371/journal.pone.0011147

221. Pockrandt C, Alzamel M, Iliopoulos CS, Reinert K. GenMap: ultra-fast computation of genome mappability. Bioinformatics. 2020;36: 3687–3692. doi:10.1093/bioinformatics/btaa222

222. Armstrong J, Hickey G, Diekhans M, Fiddes IT, Novak AM, Deran A, et al. Progressive Cactus is a multiple-genome aligner for the thousand-genome era. Nature. 2020;587: 246–251. doi:10.1038/s41586-020-2871-y

223. Hickey G, Paten B, Earl D, Zerbino D, Haussler D. HAL: a hierarchical format for storing and analyzing multiple genome alignments. Bioinformatics. 2013;29: 1341–1342. doi:10.1093/bioinformatics/btt128

224. Wilson DJ. The harmonic mean p-value for combining dependent tests. PNAS. 2019;116: 1195–1200. doi:10.1073/pnas.1814092116

225. Rasmussen MD, Hubisz MJ, Gronau I, Siepel A. Genome-Wide Inference of Ancestral Recombination Graphs. PLoS Genet. 2014;10: e1004342. doi:10.1371/journal.pgen.1004342

226. Schliep KP. phangorn: phylogenetic analysis in R. Bioinformatics. 2011;27: 592–593. doi:10.1093/bioinformatics/btq706

227. Revell LJ. phytools: an R package for phylogenetic comparative biology (and other things). Methods in Ecology and Evolution. 2012;3: 217–223. doi:10.1111/j.2041-210X.2011.00169.x

228. Dowle M, Srinivasan A, Gorecki J, Chirico M, Stetsenko P, Short T, et al. data.table: Extension of “data.frame.” 2021. Available: https://CRAN.R-project.org/package=data.table

229. Wickham H, François R, Henry L, Müller K, RStudio. dplyr: A Grammar of Data Manipulation. 2022. Available: https://CRAN.R-project.org/package=dplyr

230. Wright K. pals: Color Palettes, Colormaps, and Tools to Evaluate Them. 2021. Available: https://CRAN.R-project.org/package=pals

231. Kelleher J, Thornton KR, Ashander J, Ralph PL. Efficient pedigree recording for fast population genetics simulation. PLOS Computational Biology. 2018;14: e1006581. doi:10.1371/journal.pcbi.1006581

232. Kelleher J, Lohse K. Coalescent Simulation with msprime. In: Dutheil JY, editor. Statistical Population Genomics. New York, NY: Springer US; 2020. pp. 191–230. doi:10.1007/978-1-0716-0199-0_9

233. Introduction — PySLiM manual. [cited 3 Oct 2022]. Available: https://tskit.dev/pyslim/docs/stable/introduction.html

234. argparse — Parser for command-line options, arguments and sub-commands — Python 3.10.7 documentation. [cited 3 Oct 2022]. Available: https://docs.python.org/3/library/argparse.html

235. Martín Abadi, Ashish Agarwal, Paul Barham, Eugene Brevdo, Zhifeng Chen, Craig Citro, et al. TensorFlow: Large-Scale Machine Learning on Heterogeneous Systems. 2015. Available: https://www.tensorflow.org/

