## Supplementary material for "Genetic diversity landscapes in outcrossing and selfing *Caenorhabditis* nematodes": Captions for Supplementary figures

**S1 Fig. The number of crossover events in F2 individuals from 4 crosses of *C. remanei* inbred lines.**

A1 and A2 are crosses of ♀ PX506 ♂ PX553; whereas B1 and B2 are crosses of ♀ PX553 ♂ PX506. We observed similar distributions of the number of crossover events in all crosses. Autosomes had more recombination events than the sex chromosome (X).

**S2 Fig. Genomic diversity statistics in *C. elegans* and *C. remanei* populations.**

Dots represent the diversity statistics estimated in 100-Kb non-overlapping windows, whereas lines show locally weighted smoothing of these values. Windows with less than 10% covered positions were removed from the analysis. The vertical dashed lines indicate the boundaries of regions of low recombination central domain. The x-axis represents the normalized genome position. See the description of statistics in the Methods section. In almost all statistics, *C. remanei* and *C. elegans* exhibit distinct patterns and scales.

**S3 Fig. Landscape of nucleotide substitutions in the *C. remanei* PX506 reference genome, as estimated from inferred ancestral states.**

The first two lines (dark teal and teal) are transitions and the other lines are various forms of transversions. **(A)** Percent of substitutions for each class as estimated from ancestral GC content as a fraction of coverage of a 1 Mb genomic window. **(B)** Relative fraction of each substitution type at a given genomic location. Overall, relative proportions for three of the substitution types are homogeneous along the genome, while the three other types (C→T|G→A, A→T|T→A, and C→G|G→C, marked with asterisks) show small but significant differences between domains of recombination (see S2 Table).

**S4 Fig. Two-dimensional representation of relatedness and structure in *C. remanei* and *C. elegans* population.**

LD1 and LD2 show two latent dimensions. Some individuals in *C. remanei* population are closely related. In the *C. elegans* population, there are few lines with several individuals that are almost genetically identical and were combined to isotypes in previous studies ([83], see S3 Table).

**S5 Fig. Genome wide patterns of linkage disequilibrium.**

The panels show linkage disequilibrium ($r^2$) in the *C. elegans* **(A)** and *C. remanei* **(B)** populations. The linkage between and within chromosomes is highly similar in *C. elegans*, but significantly different (see the main text). *C. remanei* shows the fast decay of linkage disequilibrium (Fig 4) and low interchromosomal LD.

**S6 Fig. Genome-wide landscape of recombination inferred from population diversity data of *C. elegans* and *C. remanei*.**
The x-axis shows the normalized genome position. The vertical dashed lines indicate the boundaries of central regions of low recombination obtained from genetic maps.

**S7 Fig. Demographic history of populations of *C. elegans* and *C. remanei* inferred from each chromosome.**

The color represents chromosomes. We ran 100 bootstrapped replicates using eight individuals from each species, each line represents one replicate. The grey shadow indicates the region of recent demographic history, where estimations are less accurate. We used one generation per year in this analysis and scaled of the mutation rate (x0.5) and coalescent time (x2) for *C. elegans*.

**S8 Fig. Analysis of inferred genome-wide genealogies of the *C. remanei* population.** **(A)** Demographic history of the *C. remanei* population estimated for each chromosome. **(B)** Tree statistics calculated from the genealogies averaged for 100kb windows; TMRCA (time to the most recent common ancestor), RTH (relative TMRCA half-time), and the lengths of terminal branches of the trees. **(C)** Signatures of positive selection along the genome, the y-axis shows the p-values after the correction on multiple comparisons using the harmonic mean approach (see the Methods). **(D)** Quantile-quantile plot displays p-values from the tests for positive selection (y-axis) versus the expected uniform distribution of p-values (x-axis). The yellow color shows sites on the arms, and the black color indicates sites on the central parts of chromosomes.

**S9 Fig. The distribution of selection and dominance coefficients of beneficial and deleterious mutations in outcrossing simulated populations.**
This picture depicts mutations from the SD&SB-SD&SB class described in S4 Table with the uniform mutation landscape. **(A)** Percentage of mutation classes with allele frequency more than 0.5 at the beginning of the simulation (“Initial”) and the end on the arms and centers. Colors display the class of dominance coefficient (h), and the columns represent the strengths of selection (absolute values of Ns, where N is the population size of 5,000 and s is the selection coefficient). **(B)** The percent of corresponding mutation classes of mutations with allele frequency less than 0.5.

**S10 Fig. Distributions of diversity statistics in simulated populations**.

Lines represent locally weighted smoothing of the values estimated per 40-kb sliding non-overlapping windows, the vertical dashed lines indicate the boundaries of central domain with low recombination rate. Columns show the outcrossing rate, where “outcrossing” means completely outcrossing populations and, in other columns, % specify the percentage of selfing in population; "outcrossing inbreeding" corresponds to scenarios with outcrossing populations that underwent the bottleneck at the very end of simulations (see Methods). Rows represent domain-specific differences in mutation rate, with 1-1-1 is the uniform mutation landscape, 1.15-1-1.15 means 15% more mutations on the arms, 1.5-1-1.5 is 50% more mutations in domains of high recombination, and 2-1-2 means two times more mutations in domains of high recombination. Colors show the selection regime (see details in Methods). On this figure, shown only 4 selection regimes that are specified with asterixis in S4 Table.

**S11 Fig. Distributions of diversity statistics in simulated populations for scenarios with neutral and deleterious and beneficial mutations.**
See description to S10 Fig and parameters in S4 Table.

**S12 Fig. Distributions of diversity statistics in simulated populations for scenarios with neutral and deleterious mutations.**
See description to S10 Fig and parameters in S4 Table.

**S13 Fig. Differences in diversity statistics between simulations with shifts in the population size compared to the values of the statistics of the corresponding simulation before the changes in neutral scenarios.**
The colors represent the fold change in statistics at the end of the simulation versus before changes in size. **(A)** Fluctuation in population size for 100 generations, where every five generations, the population size went from 5,000 to 15.000 and then back. **(B)** Exponential growth of 3 % for 100 generations.

**S14 Fig. Model accuracy and cross-entropy loss curves for the convolutional neural network.**
The orange color shows the validation accuracy or loss, and the teal color indicates values for training.

**S15 Fig**. **The confusion matrix for the convolutional neural network.**

The color represents the count of classified simulations. The values on the axes represent the simulated evolutionary scenario, where the first number denotes the mutation rate increase on the arms relative to the central domains, e.g., 1.5 is 50% more mutations on the arms. The second values show the selfing rate, where “0.0” stands for the outcrossing populations, “0.0i” are outcrossing with inbreeding at the end of simulations, and for instance, 0.9 is 90% selfing in the population. The third value indicated the selection regime, with “N” for neutral simulation, “ND” for neutral and deleterious, and “NDB” for neutral, deleterious, and beneficial mutations. Most values are located around the main diagonal, where the predictions match the simulated scenarios. The two off-diagonals indicate confusion between the adjacent mutation rates. For instance, when the mutation rate on the arms was 15% higher the network sometimes predicted either the uniform or 50% more mutations on the arms. Additionally, there is a confusion around some of the neutral and neutral/deleterious scenarios (as seen by 2x2 blocks on the diagonals), probably because the fraction of nearly-neutral mutations in some simulations is high (see S9 Fig and S20 Fig).

**S16 Fig. Principal component analysis of simulated population.**
PC1 and PC2 are the first and second principal components. Each dot shows one simulation from 41,840 featured here. Colors indicate different parameter values for simulation scenarios. **(A)** Mutation rate variation, where “1” is the uniform mutation rate, and “1.5” is 50% more mutations on the arms relative to the central domains. **(B)** Selfing rate in the simulated population, “0.0” denoted the outcrossing populations, “0.0i” outcrossing with inbreeding at the end of simulations, and for instance, “0.999” represents 99.9% of selfing in the population. **(C)** Selection regime. “N” shows neutral simulations, “ND” is neutral and deleterious mutations, and “NDB” is neutral, deleterious, and beneficial mutations.

**S17 Fig. Principal component analysis of simulated populations and position of chromosomes of *C. elegans* and *C. remanei* projected to the PCA space.**
For each chromosome, we did 50 bootstrapped replicates sampling statistics from domains of low and high recombination. Colors indicate the chromosomes of *C. elegans* **(A)** and *C. remanei* **(B)**. Sex chromosomes and some autosomes with long tracts of repetitive sequences were removed from the analysis.

**S18 Fig. Saliency plot for one of the bootstrapped replicates of chromosome I from the *C. remanei* population by the neural network.**

The teal color shows the original diversity landscapes of 9 statistics. That replicate was classified as (1.15, 0.9, ND), meaning 15% more mutations on the arms, 90% selfing, and neutral and deleterious mutations. The orange color shows perturbed statistics, slightly changing to the original ones. However, that population would be classified as (1.5, 0.0, ND), with 50% more mutations on the arms, outcrossing with neutral and deleterious mutations. This example demonstrates how sensitive the model is to slight changes in diversity statistics.

**S19 Fig. Effects of bioinformatic filtering of the empirical data to the estimated diversity statistics.**
The colors indicate different parameters of the filters. The first value of the filter specifies the number of *C. remanei* individuals, either 14 or 17. The following two values are, correspondingly, minimal and maximal coverage, and the last is the fraction of individuals with the specified coverage. In the analysis, we used the top filter indicated by black color.

**S20 Fig. Distributions of selection and dominance coefficients used in evolutionary simulations.**
The columns show the percentage of each class of selection coefficients drawn from gamma distributions with different parameters (see S4 Table). Dominance coefficients were chosen independently from a mixture of uniform and beta distributions with distinct parameters for deleterious mutations and beneficial mutations (see the Methods and SLiM scripts at https://github.com/phillips-lab/CR_CE_popgen/simulations).
