## Supplementary tables for "Genetic diversity landscapes in outcrossing and selfing *Caenorhabditis* nematodes"

**S1 Table**. **Supplementary information for the *C. remanei* genetic map.**
Lengths of genetic maps are from joint estimation using a pedigree for 4 families with grandparents.

| **Chromosome** | **I** | **II** | **III** | **IV** | **V** | **X** |
|---|---|---|---|---|---|---|
| **Number of informative markers** | 2227 | 2027 | 1998 | 2413 | 2542 | 1389 |
| **Number of markers after filtration** | 900 | 1091 | 1381 | 1413 | 1660 | 1067 |
| **Number of F2 individual with at least 1 recombination** | 222 | 216 | 226 | 219 | 218 | 129 |
| **Length of maternal map (cM)** | 49.973 | 49.126 | 51.502 | 48.301 | 48.817 | 40.998 (females only) |

**S2 Table. The difference in the relative fraction of substitutions from the *C. latens* and *C. remanei* common ancestor to the *C. remanei* strain PX506 between "arms" and "centers".**

Approximative General Independence Test (two sided), the p-values were adjusted with the Bonferroni correction.

| Substitution | Cohen's D | Z | P-value | Adjusted P-value |
|---|---|---|---|---|
| **C→T\|G→A** | **0.5508074** | **3.612359** | **0.0005** | **0.003** |
| A→G\|T→C | 0.3517458 | 2.314029 | 0.0211 | 0.0633 |
| C→A\|G→T | 0.3025061 | -1.991209 | 0.0484 | 0.0722 |
| **A→T\|T→A** | **0.424313** | **-2.788705** | **0.0078** | **0.0312** |
| **C→G\|G→C** | **0.4358583** | **-2.864092** | **0.0045** | **0.0225** |
| A→C\|T→G | 0.3251058 | -2.139442 | 0.0361 | 0.0722 |

**S3 Table. Individually sequences worms used in this study.**
* For details on *C. elegans* data see the Supplementary material from Crombie et al. 2019, doi:10.7554/eLife.50465

| Species | ID | Isotype | SRA Accession | Collected at (latitude, longitude) |
|---|---|---|---|---|
| *C. elegans** | ECA779 | ECA760 | SRR9321974 | * the Big Island, Hawaii, USA (20.039858, -155.441608), August 2017; all *C. elegans* sequences are from Crombie et al. 2019 |
| | ECA780 | ECA760 | SRR9321975 | |
| | ECA808 | ECA760 | SRR9322800 | |
| | ECA781 | ECA778 | SRR9322495 | |
| | ECA756 | ECA778 | SRR9322785 | |
| | ECA757 | ECA778 | SRR9322428 | |
| | ECA762 | ECA812 | SRR9322881 | |
| | ECA748 | ECA812 | SRR9322898 | |
| | ECA750 | ECA812 | SRR9322348 | |
| | ECA751 | ECA812 | SRR9322262 | |
| | ECA752 | ECA760 | SRR9322261 | |
| | ECA753 | ECA760 | SRR9322784 | |
| | ECA761 | ECA812 | SRR9322880 | |
| | ECA772 | ECA768 | SRR9322091 | |
| | ECA777 | ECA777 | SRR9322085 | |
| | ECA778 | ECA778 | SRR9322022 | |
| | ECA758 | ECA760 | SRR9322414 | |
| | ECA759 | ECA812 | SRR9322945 | |
| | ECA770 | ECA807 | SRR9322093 | |
| | ECA774 | ECA812 | SRR9322089 | |
| | ECA775 | ECA812 | SRR9322088 | |
| | ECA776 | ECA812 | SRR9322087 | |
| | ECA807 | ECA807 | SRR9322543 | |
| | ECA809 | ECA812 | SRR9322799 | |
| | ECA783 | ECA812 | SRR9321978 | |
| | ECA784 | ECA760 | SRR9321979 | |
| | ECA785 | ECA760 | SRR9321980 | |
| | ECA787 | ECA812 | SRR9322542 | |
| *C. remanei* | A24-1 | – | SRR17772309 | Koffler Scientific Reserve, Jokers Hill, Ontario, Canada (44.029608, -79.531142), September 2013 |
| | A31-1 | – | SRR17772314 | |
| | N15-2 | – | SRR17772305 | |
| | NS16-2 | – | SRR17772304 | |
| | NS26 | – | SRR17772312 | |
| | NS30 | – | SRR17772310 | |
| | NS33-2 | – | SRR17772315 | |
| | NS4 | – | SRR17772306 | |
| | NS50-1 | – | SRR17772303 | |
| | NS50-2 | – | SRR17772302 | |
| | NS52 | – | SRR17772311 | |
| | NS56 | – | SRR17772308 | |

| | | | | |
|---|---|---|---|---|
| | NS59-2 | – | SRR17772313 | |
| | R18-1 | – | SRR17772307 | |
| | PX553 | – | SRR17714788 | Inbred strain (28 generations of inbreeding) generated from strain PX393 collected at the Koffler Scientific Reserve, Jokers Hill, Ontario, Canada (44.029608, -79.531142), October 2008 |
| | PX553 X PX506 F2 | – | SRR17714789-SRR17714792 | Best-RAD data for F2 females from four crosses of PX506 and PX553 strains |

**S4 Table. Distribution of selection coefficients used in evolutionary simulations.**

The selection coefficients were drawn from a gamma distribution with a mean of Ns and a scale of 0.3. The right column is showing the number of replicates for each combination of parameters. * Decay simulations only for these scenarios (50 replicates).

| **Scenario** | **Arm-Center** | **Description, (mean selection parameter Ns)** | **Replicates** |
|---|---|---|---|
| Neutral | * | Only neutral mutations | 600 |
| Neutral + Deleterious | WD-WD | “Weak” (Ns=3) deleterious mutations in both recombination domains | 200 |
| | MD-SD | “Moderate” (Ns=7.5) on the arms, “strong” (Ns=15) on the centers | 200 |
| | SD-SD* | “Strong” (Ns=15) deleterious in both domains | 200 |
| Neutral + Deleterious + Beneficial | WD&SB-WD&SB | “Weak” (Ns=3) deleterious mutations and “strong” (Ns=15) beneficial in both recombination domains | 200 |
| | MD&MB-SD&SB | “Moderate” (Ns=7.5) deleterious and beneficial on the arms and “strong” (Ns=15) on the centers | 200 |
| | SD&SB-SD&SB* | “Strong” (Ns=15) for both types along the chromosome | 200 |
| Neutral + Deleterious + Balancing | SD&SBa-SD&SBa* | “Strong” (Ns=15) for both types along the chromosomes, selection for beneficial mutation is dependent on allele frequency in populations (max at 0.5) | 200 |
