## Supplementary figures and images for "Genetic diversity landscapes in outcrossing and selfing *Caenorhabditis* nematodes"

### Supplementary Figure S1

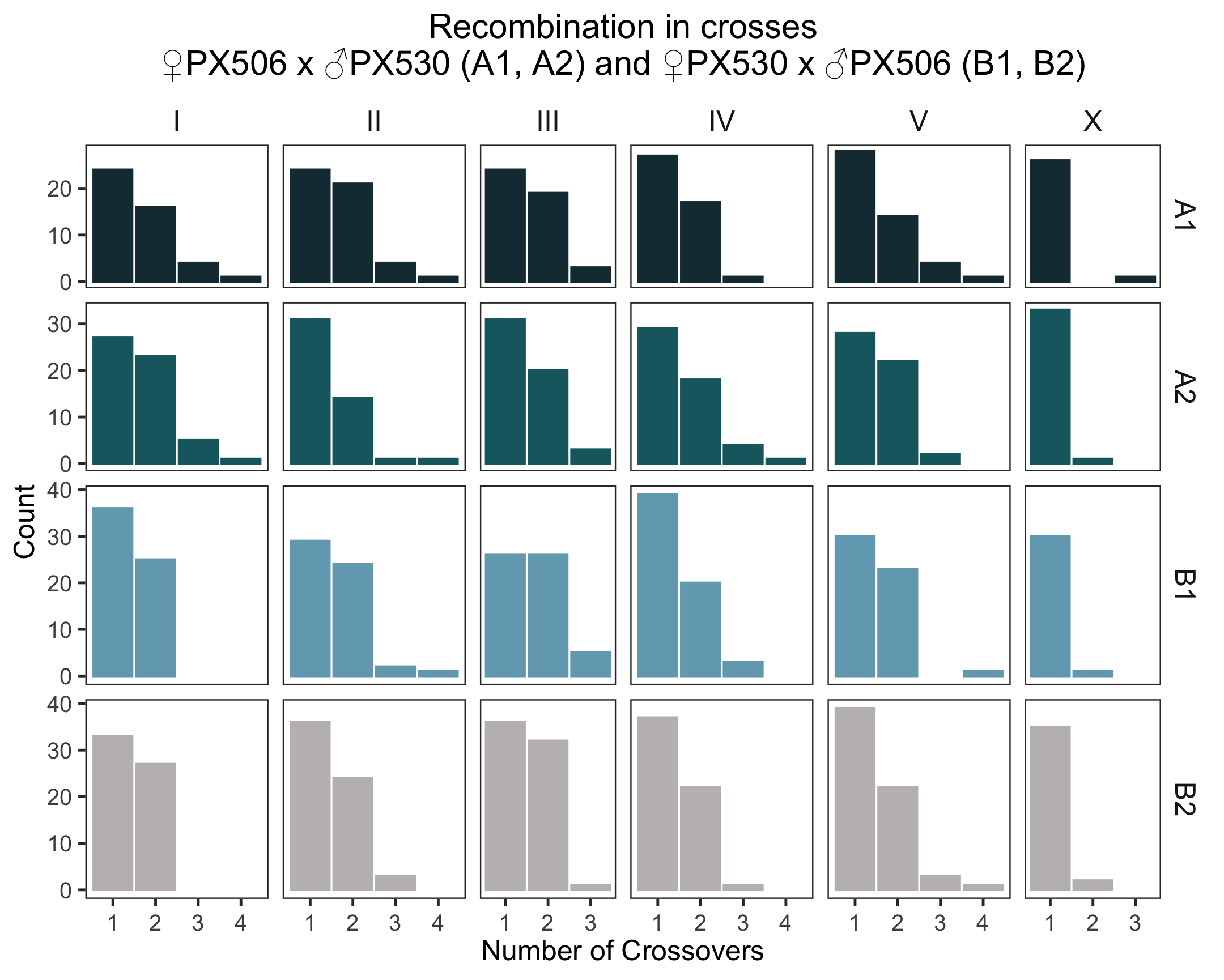

### Supplementary Figure S2

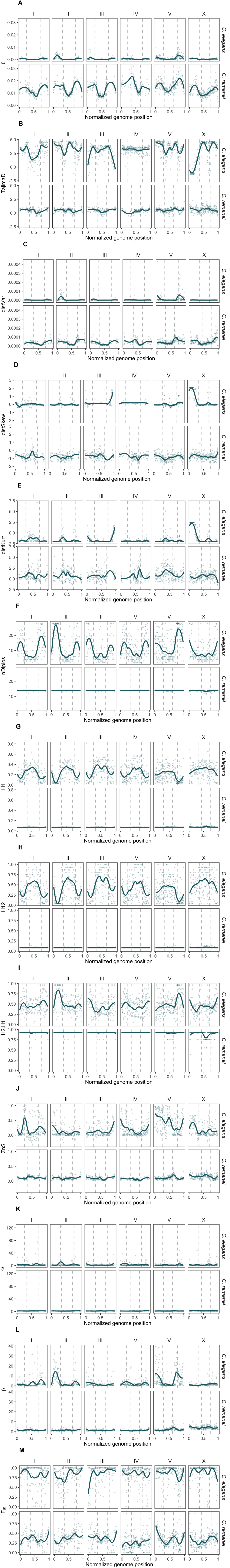

### Supplementary Figure S3

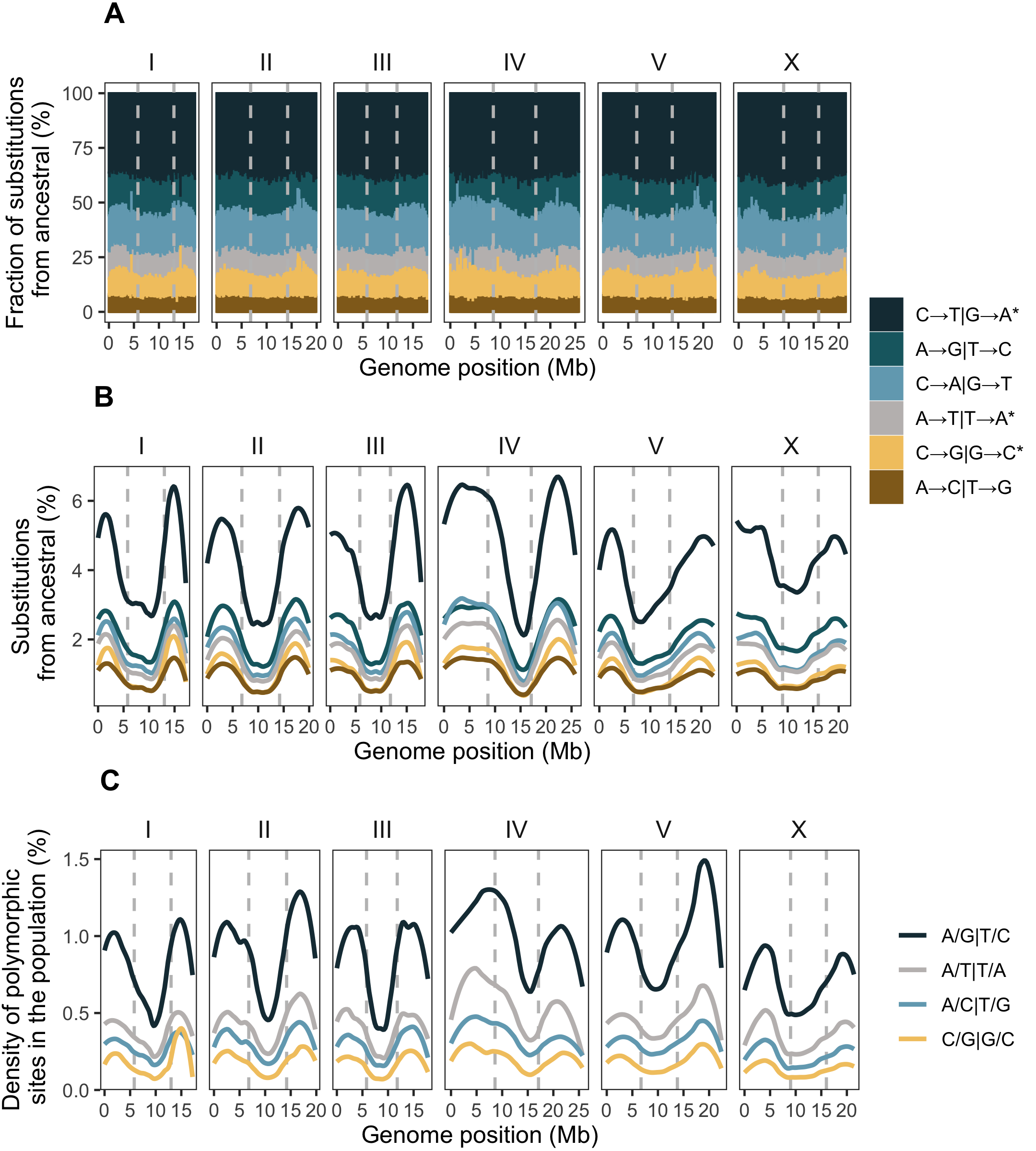

### Supplementary Figure S4

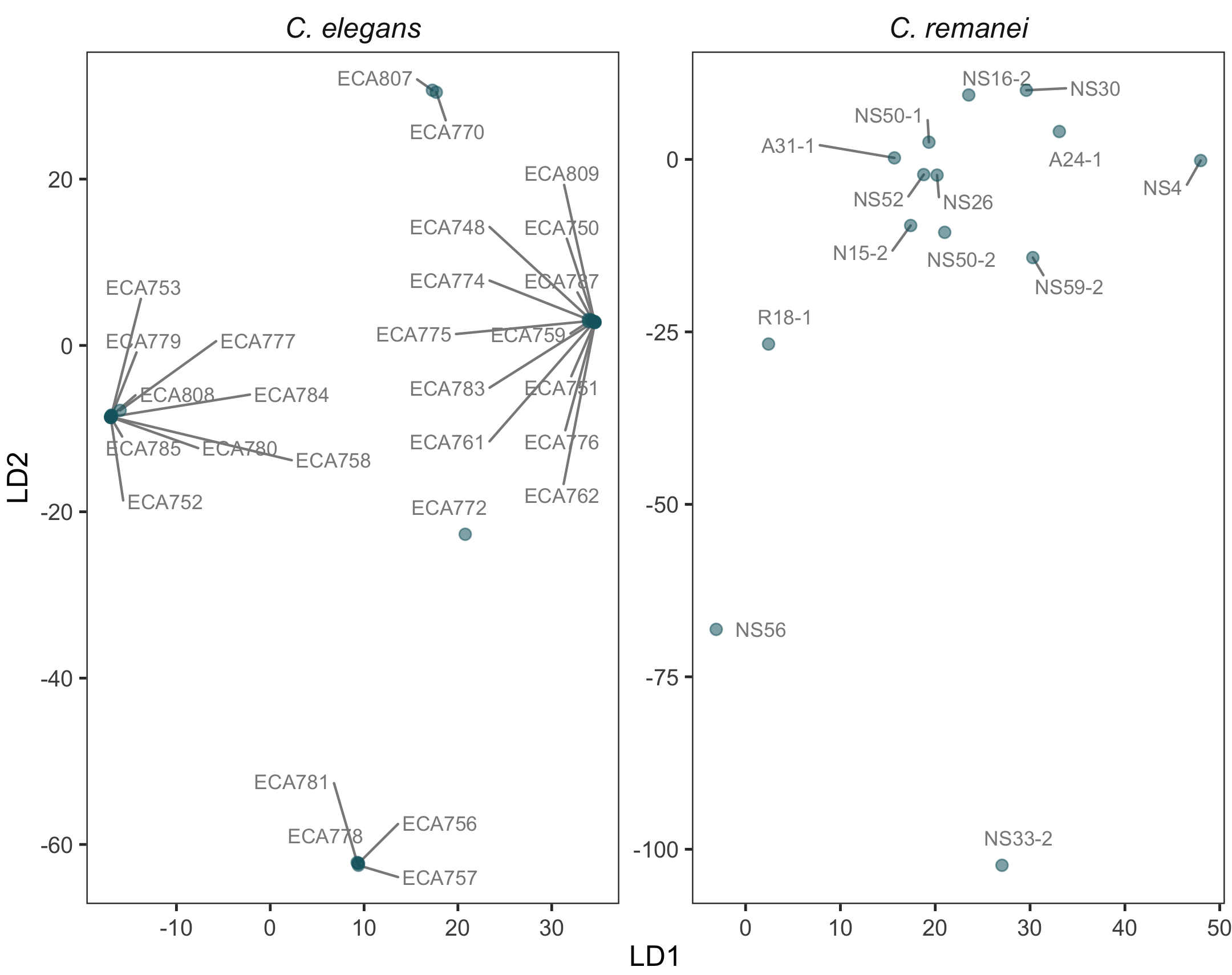

### Supplementary Figure S5

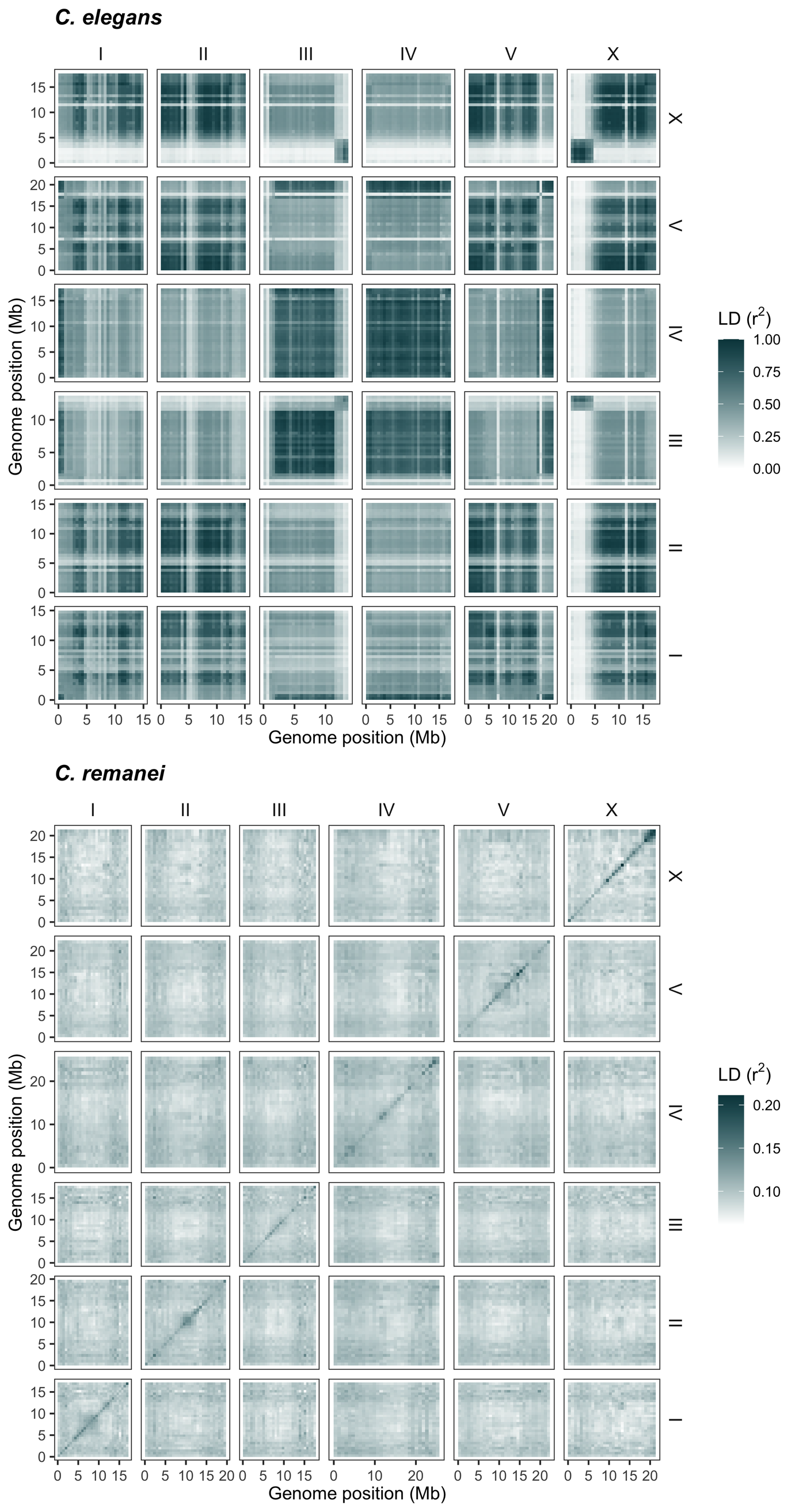

### Supplementary Figure S6

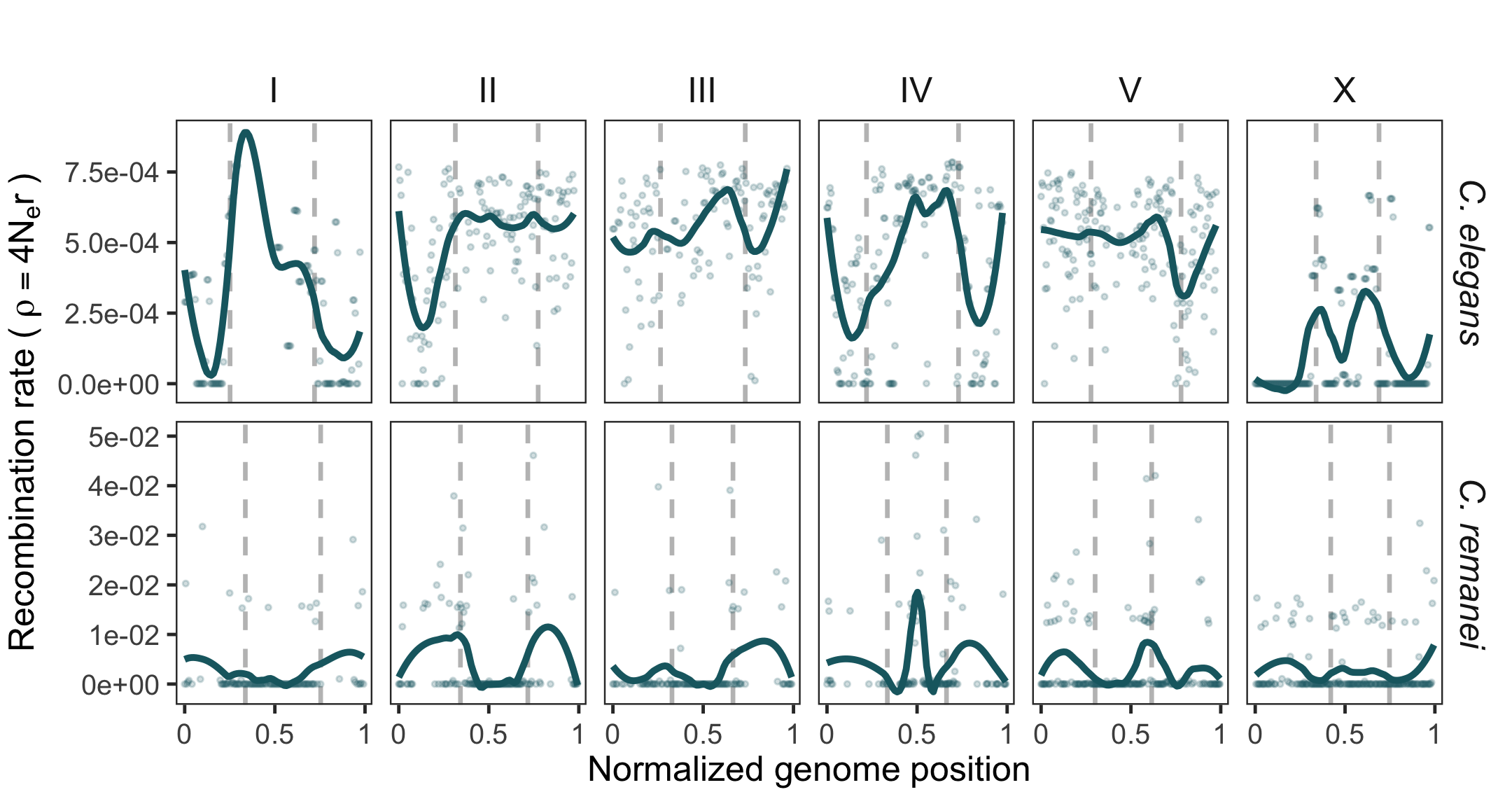

### Supplementary Figure S7

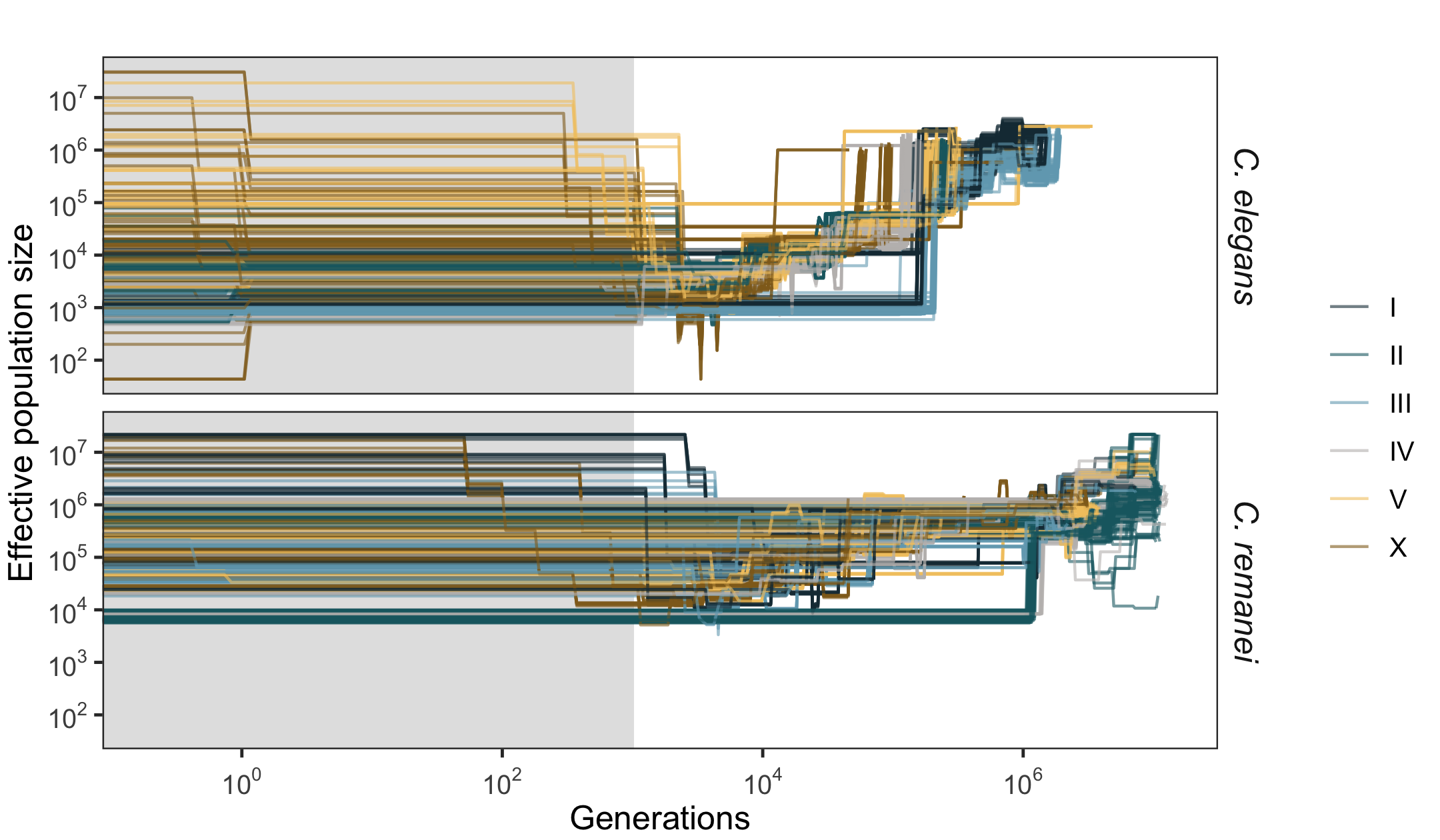

### Supplementary Figure S8

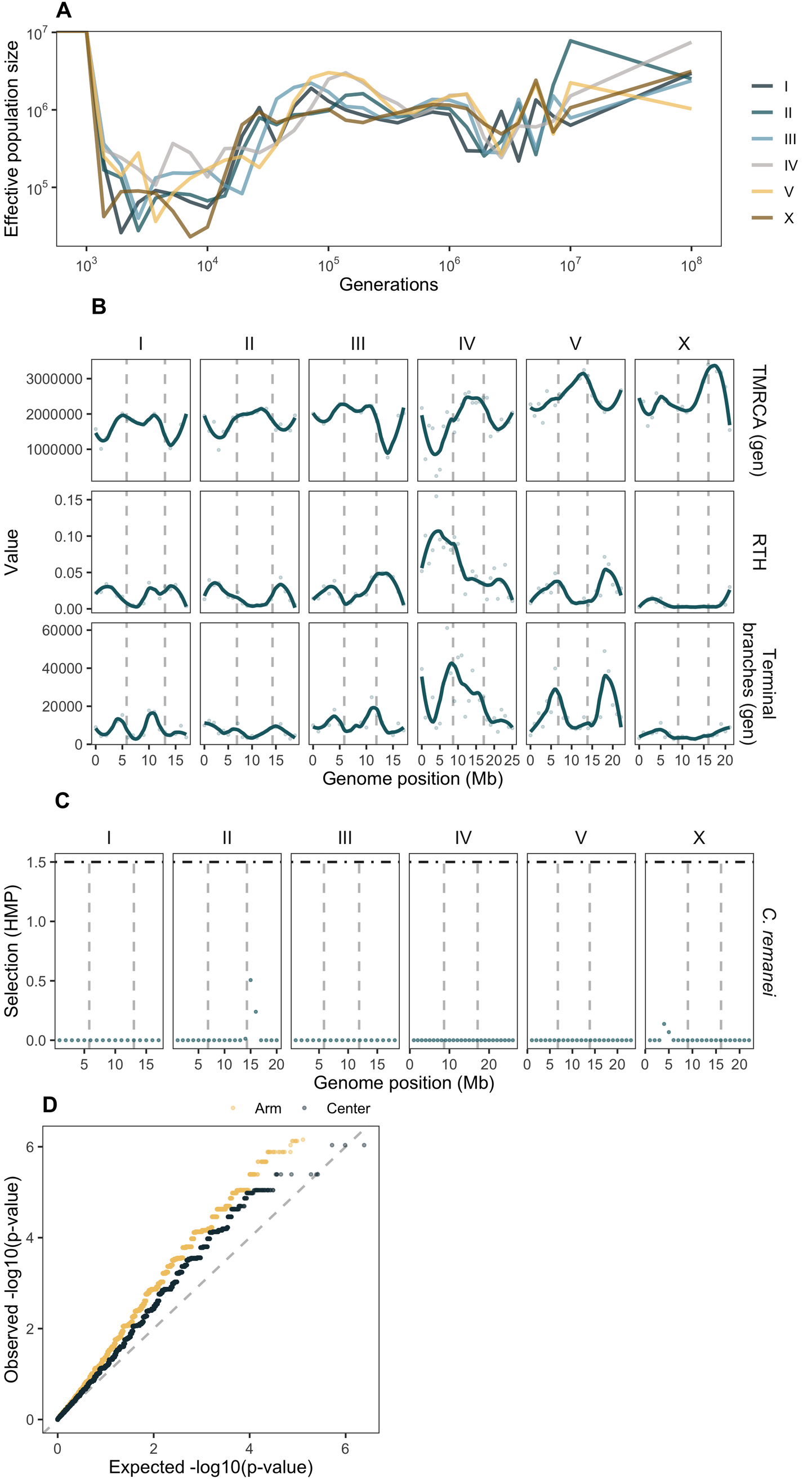

### Supplementary Figure S9

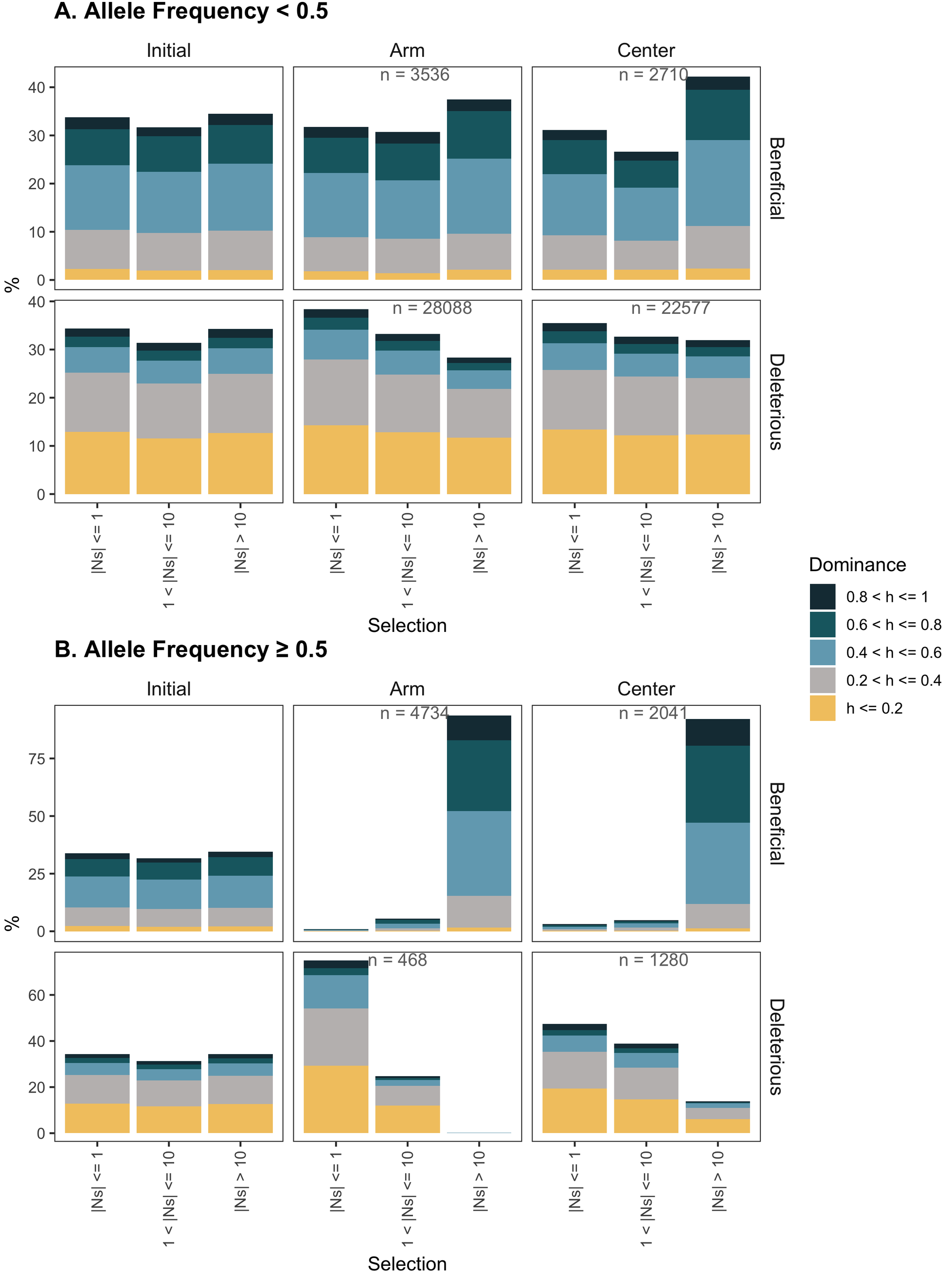

### Supplementary Figure S10

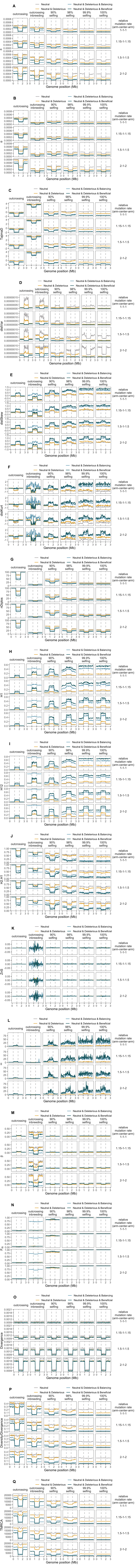

### Supplementary Figure S11

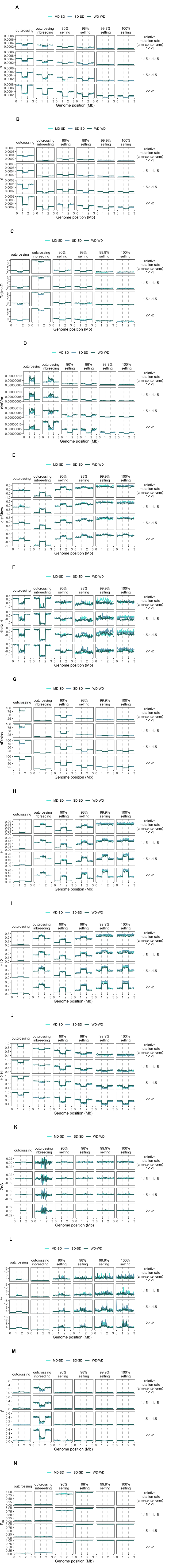

### Supplementary Figure S12

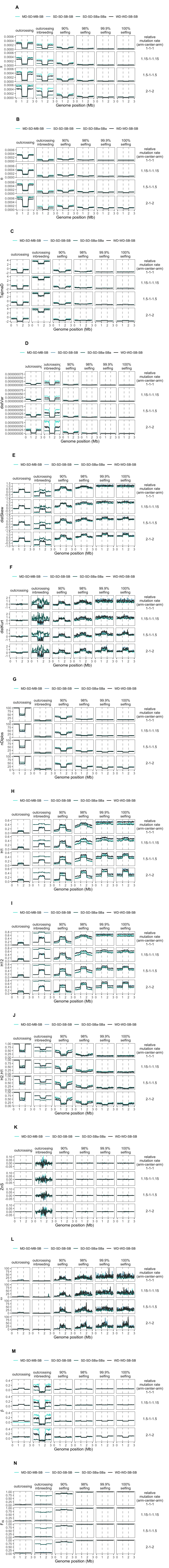

### Supplementary Figure S13

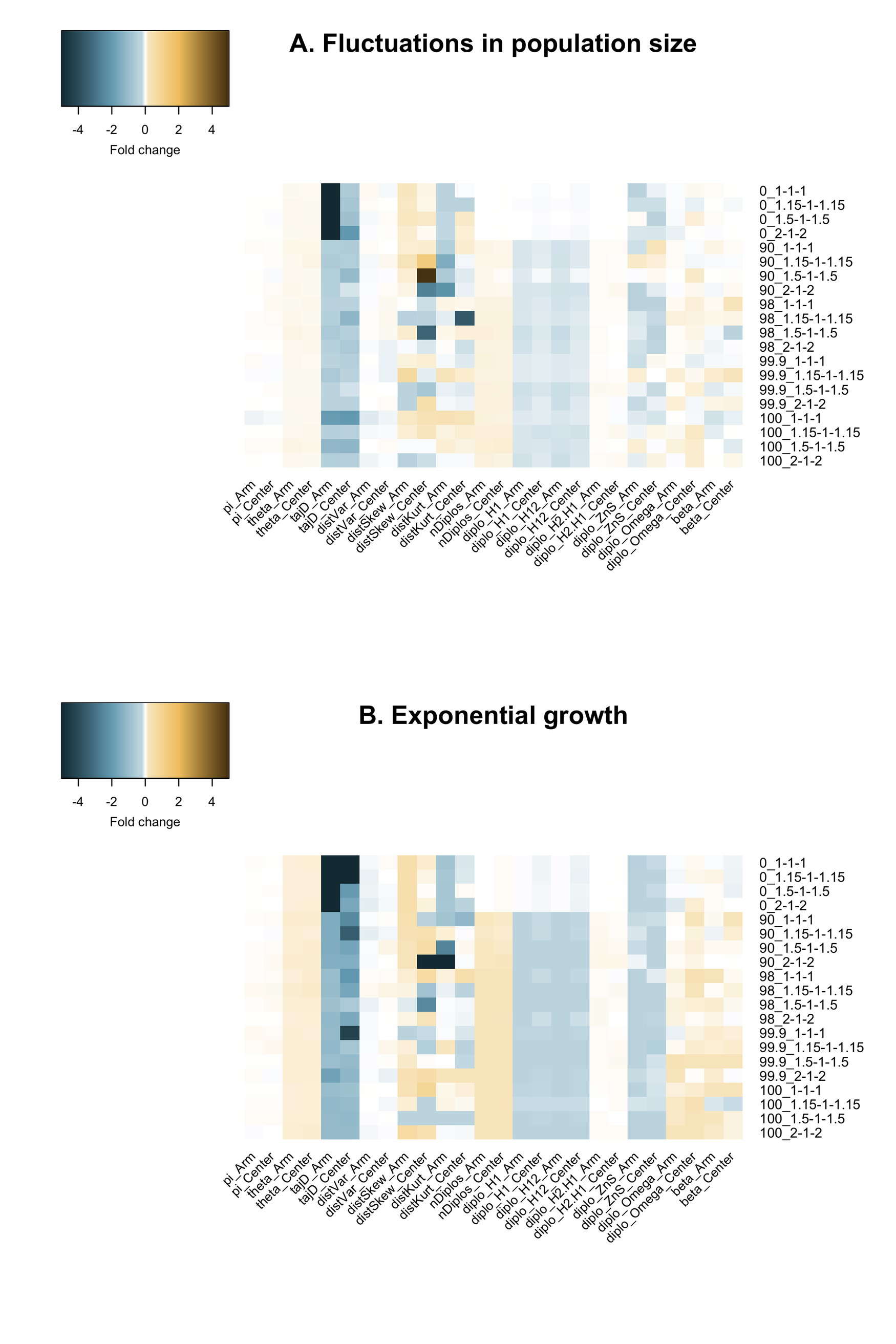

### Supplementary Figure S14

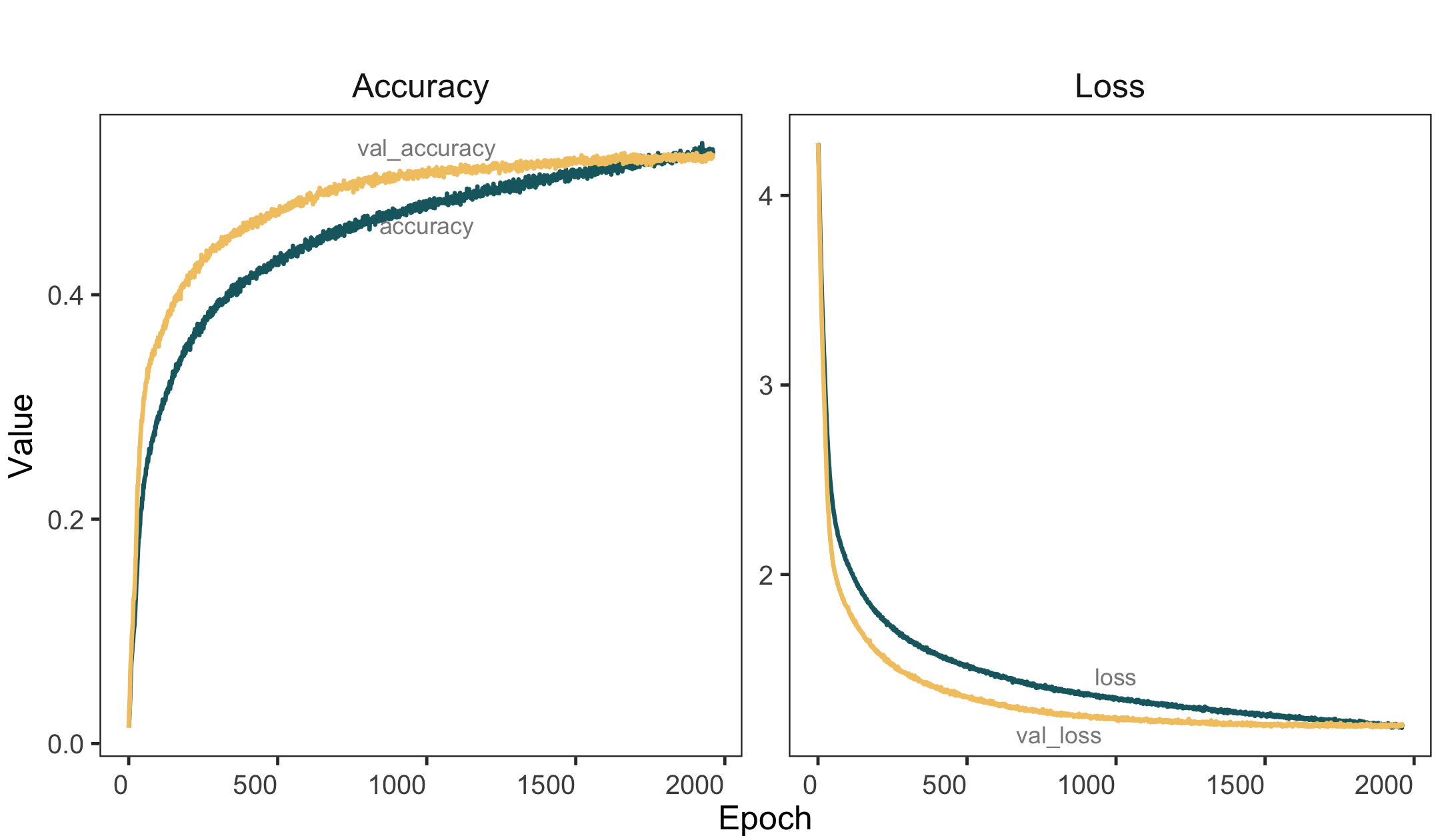

### Supplementary Figure S15

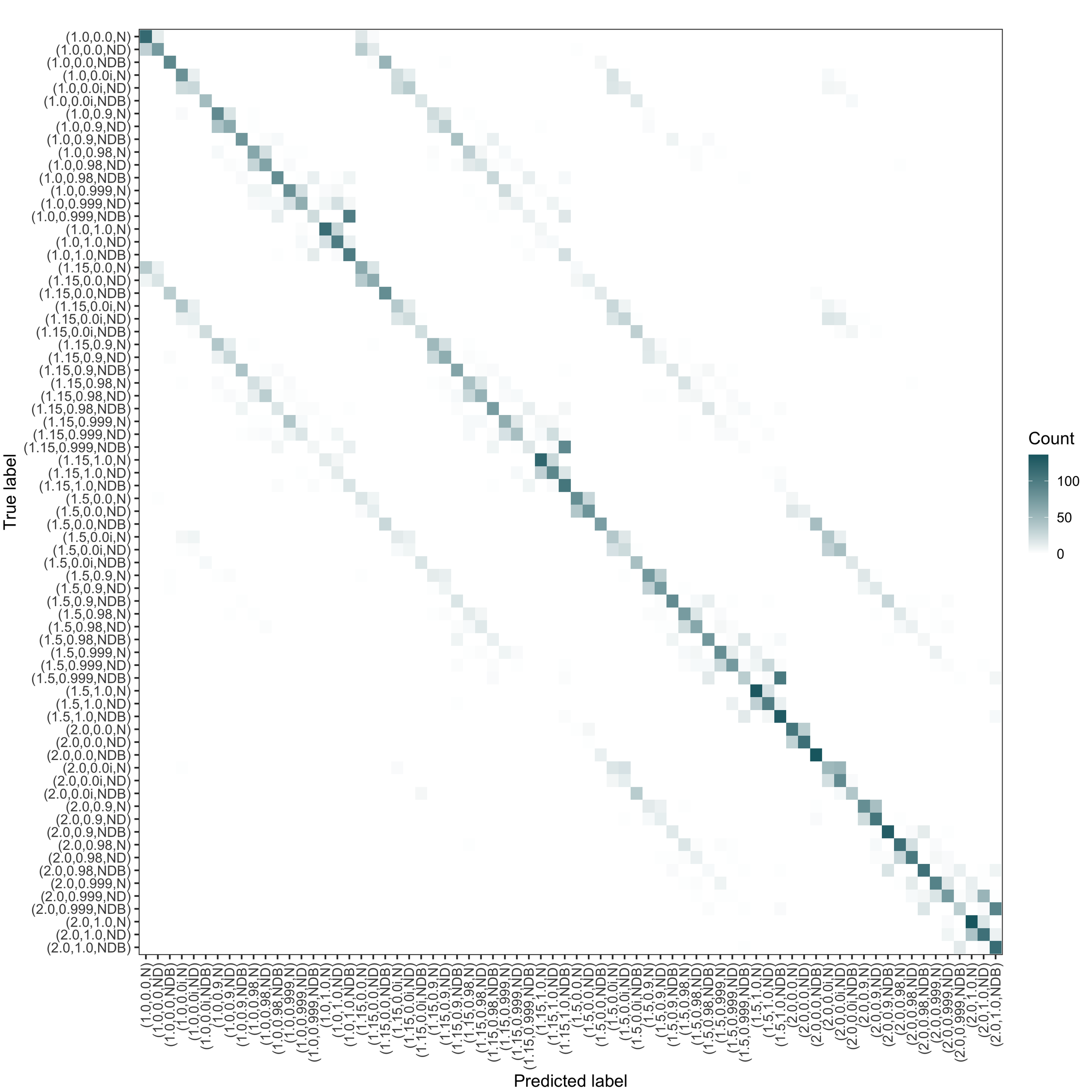

### Supplementary Figure S16

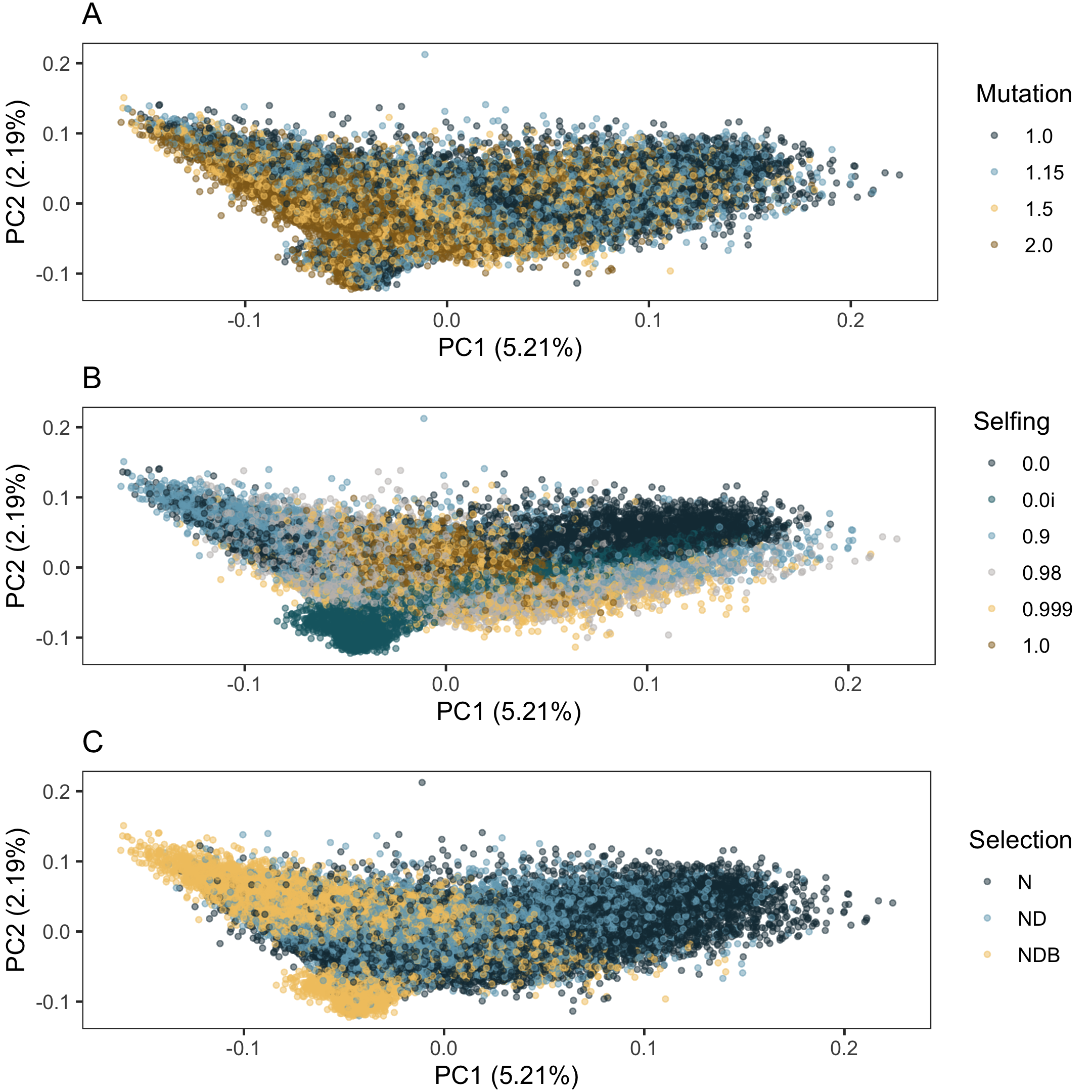

### Supplementary Figure S17

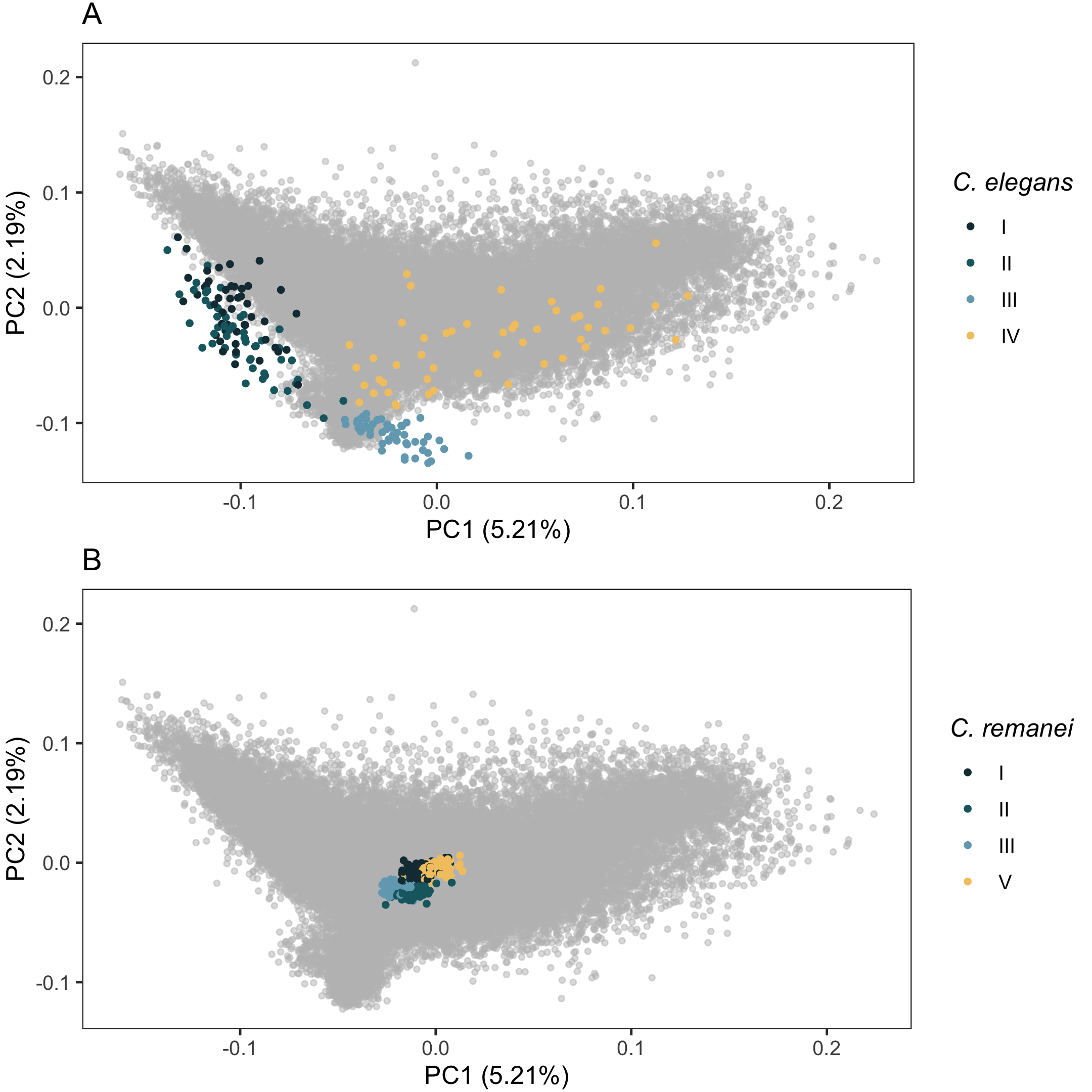

### Supplementary Figure S18

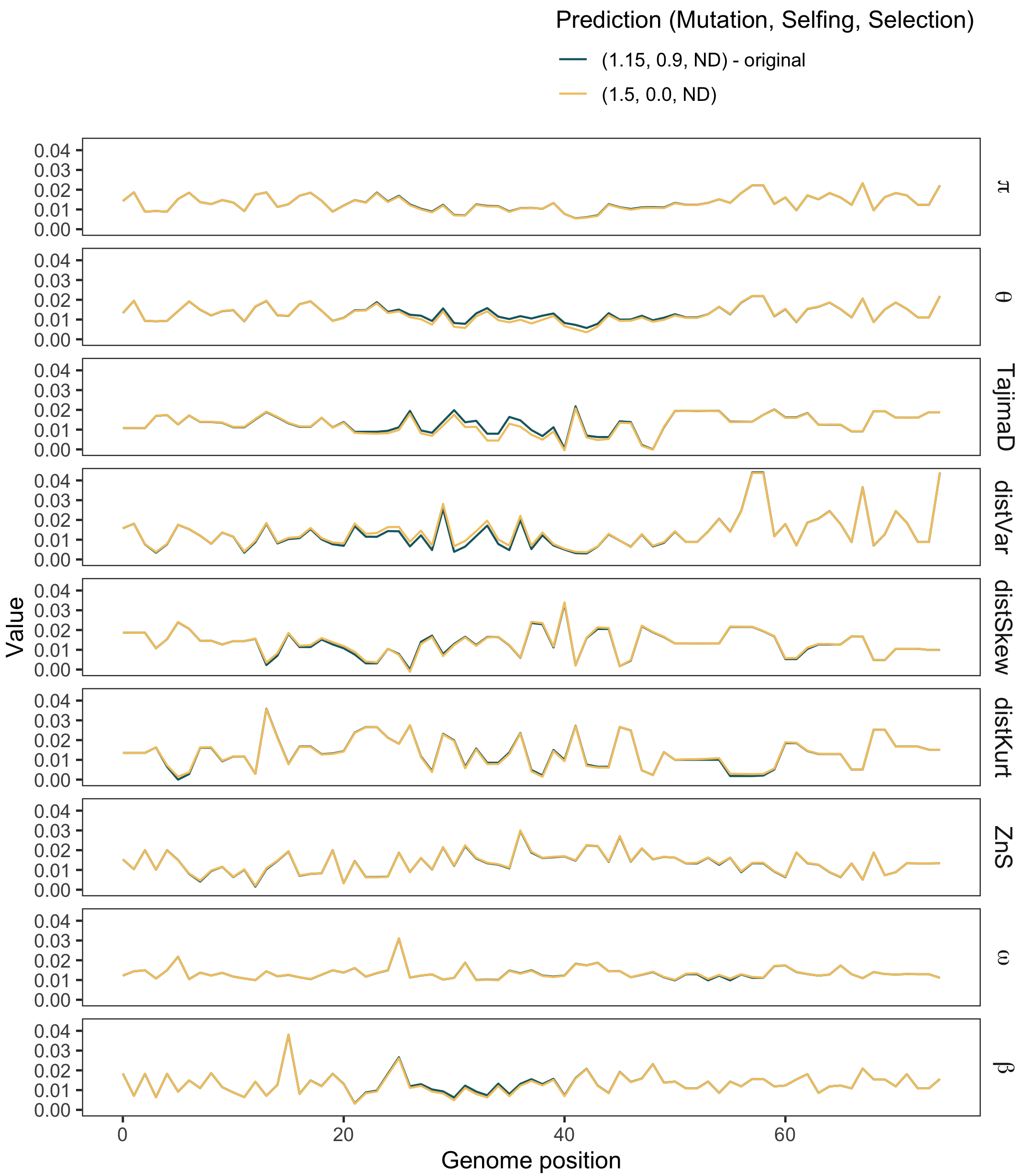

### Supplementary Figure S19

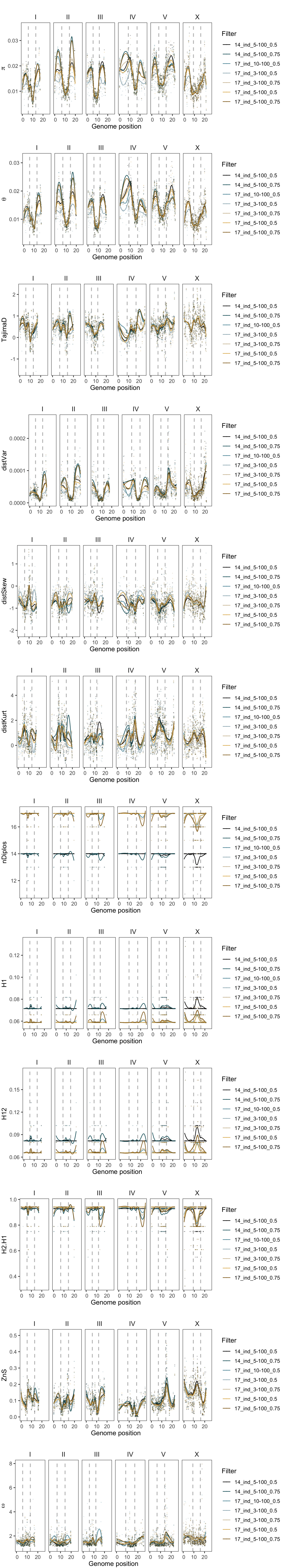

### Supplementary Figure S20

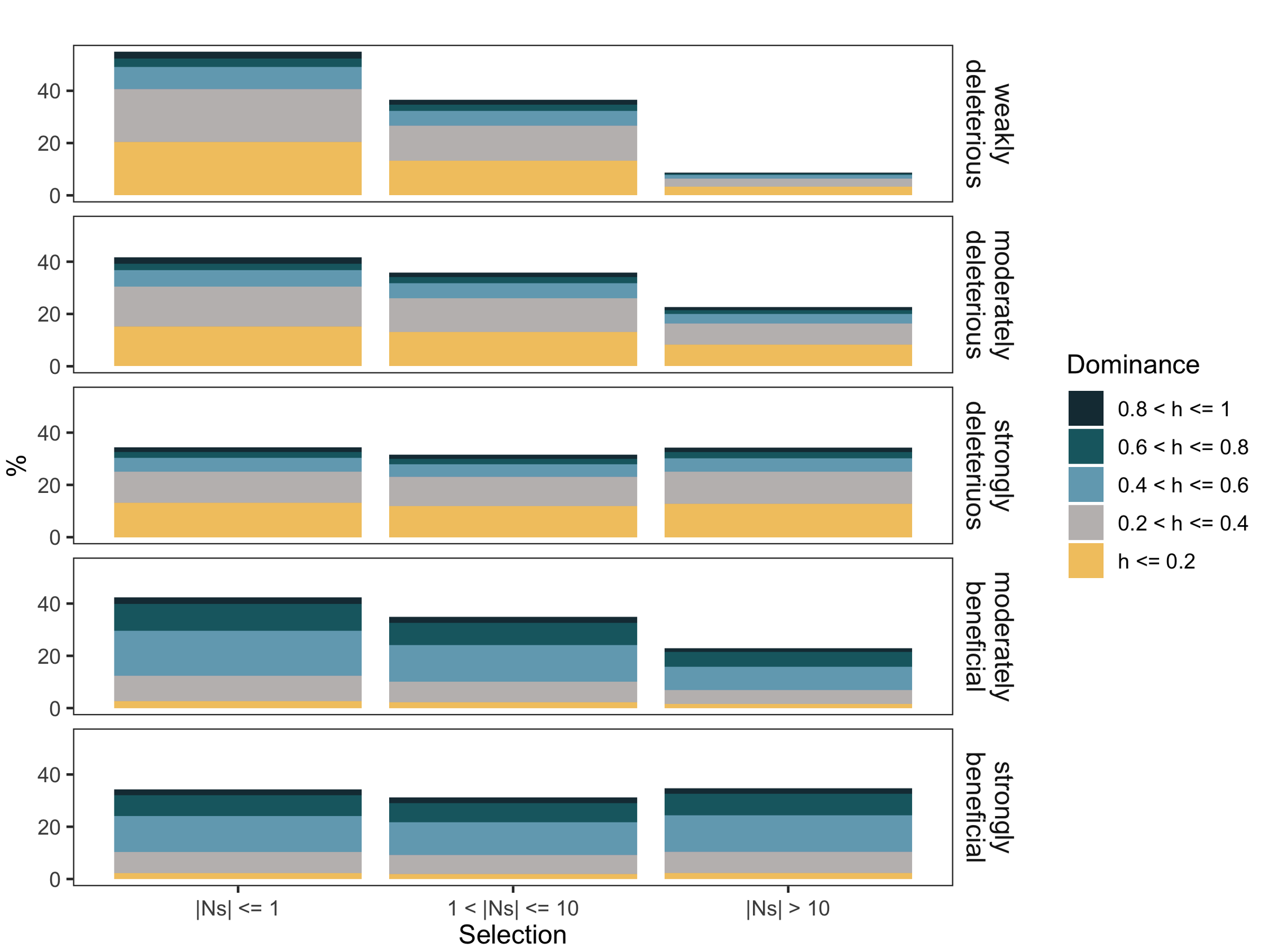
